# Evidence for the Existence of a Bacterial Etiology for Alzheimer’s Disease and for a Temporal-Spatial Development of a Pathogenic Microbiome in the Brain

**DOI:** 10.1101/2022.08.28.505614

**Authors:** Yves Moné, Joshua P. Earl, Jarosław E. Król, Azad Ahmed, Bhaswati Sen, Garth D. Ehrlich, Jeffrey R. Lapides

## Abstract

**Background:** Over the last few decades, a growing body of evidence suggests a role for various infectious agents in Alzheimer’s Disease (AD) pathogenesis. Despite diverse pathogens (virus, bacteria, or fungi) being detected in AD subjects’ brains, most research has focused on individual pathogens and only a few studies investigated the hypothesis of a bacterial brain microbiome. We profiled the bacterial communities present in non-demented controls and AD subjects’ brains.

**Results:** We obtained post-mortem samples from the brains of 32 individual subjects, comprising 16 AD and 16 control aged-matched subjects with a total of 130 samples from the frontal and temporal lobes and entorhinal cortex. We used full-length 16S rRNA gene amplification with Pacific Biosciences sequencing technology to identify the bacteria.

We detected bacteria in the brains of both cohorts with the principal bacteria comprising *Propionibacterium acnes* (recently renamed *Cutibacterium acnes)* and two species each of *Acinetobacter* and *Comamonas* genera. We used a hierarchical Bayesian method to detect differences in relative abundance among AD and control groups. Because of large abundance variances we also employed an unconventional analysis approach that utilized Latent Dirichlet Allocation, often used in computational linguistics. This allowed us to identify 5 classes of samples, each revealing a different microbiome. Assuming that samples represented infections that potentially began at different times, we ordered these classes in time, finding that the last class exclusively explained the existence or non-existence of AD.

**Conclusions:** The AD-related pathogenicity of the brain microbiome seems to be based on a complex polymicrobial dynamic. The time ordering revealed a rise and fall of the abundance of *Propionibacterium acnes* with pathogenicity occurring for an off-peak abundance level in association with at least one other bacterium from a set of genera that included: *Methylobacterium, Bacillus, Caulobacter, Delftia,* and *Variovorax. P. acnes* may also be involved with outcompeting the *Comamonas* species, which were strongly associated with non-demented brain microbiome, whose early destruction could be the first stage of the disease. The statistical results are also consistent with a leaky blood brain barrier or lymphatic network that allows bacteria, viruses, fungi, or other pathogens to enter the brain.

## INTRODUCTION

More than a century ago, Oskar Fischer [1,2] and then Alois Alzheimer [3] independently described, the two histopathological hallmarks of a neurodegenerative disorder which is now called Alzheimer’s disease: amyloid-β plaques and neurofibrillary tangles [4]. Alzheimer’s disease (AD) is the most common neurodegenerative disease in the elderly, accounting for an estimated 60% to 80% of cases of dementia. AD patients are affected by memory loss and a progressive decline of cognitive abilities (thinking, language, behavior changes) [5]. The majority of AD cases are sporadic, late-onset forms of the disease occurring after the age of 65 years with only asmall percentage of cases (around 5%), mostly familial, presenting earlier [6].

AD is characterized by neuroinflammation, extracellular deposition of amyloid-β (Aβ) peptides into plaques in the brain parenchyma and intraneuronal neurofibrillary tangles (NFT) that ultimately lead to a loss of neurons and synapses. Aβ deposition has been considered as the main cause of the disease leading to the ‘amyloid cascade hypothesis’ as a model of AD pathogenesis [7–9]. Aβ peptides are produced through the abnormal processing of the Aβ precursor protein (APP) by the sequential action of β- and γ-secretases. This amyloidogenic processing produces Aβ peptides differing in length, including the highly pathogenic and aggregation-prone Aβ42 (42 amino acids) and the less neurotoxic Aβ40 (40 amino acids) [10–13]. Aβ peptides aggregate into oligomers, fibrils and plaques in the extracellular space. Aβ is also involved in the formation of NFT by induction of hyperphosphorylation of the tau protein (a microtubule-associated protein) via the kinase Fyn [14–17]. Although the amyloid cascade hypothesis has guided much of AD research for the last several decades, multiple observations challenge this model. First, the amyloid cascade hypothesis is based on the study of the genetic mutations observed in the rare early onset forms of AD, and clinical trials targeting Aβ accumulation have not resulted in any noticeable success. Moreover, the quantitative level of Aβ does not correlate with the amount of cognitive decline and a substantial proportion of healthy elderly subjects (10-30%) show significant amyloid deposition [18–20] (see [7] for counter arguments) upon autopsy following death from other causes.

In recent years, a growing body of evidence has suggested a role for various infectious agents (virus, bacteria, fungi) as well as the innate immune system and neuroinflammatory pathways in AD pathogenesis leading to the emergence of alternative models variously called the ‘pathogen hypothesis’ (or ‘infectious hypothesis’) and ‘antimicrobial protection hypothesis’ [21–25]. Diverse pathogens have been detected in the brains of AD patients. Viruses particularly from the Herpesviridae family have been long suspected to play a role in AD (reviewed in [12,26]). Herpes simplex virus type 1 (HSV1) has been found to be active in brains from non-demented elderly as well as in AD patients and to be localized within amyloid plaques [27]. A retrospective cohort study from Taiwan showed that subjects with HSV infections may have a 2.56-fold increased risk of developing dementia and that anti-herpetic treatment of HSV infections was associated with a decreased risk of dementia [28]. Recent findings suggest that Herpesviridae infections could contribute directly to amyloid deposition [29,30], nonetheless thepotential role of Human Herpesvirus 6 and 7 in AD pathogenesis [31] remains controversial [32–36].

Another body of work has associated bacteria with an etiological role in AD pathogenesis. The presence of spirochetes including the Lyme disease agent, *Borrelia burgdorferi,* and the periodontal *Treponemal spp.* pathogens have been repeatedly identified in post-mortem AD brains. Moreover, tertiary syphilis produces a dementia, general paresis, with a neurohistopathology complete with Aβ amyloid and NFT and associated behavioral changes essentially identical to AD [37,38]. Other bacterial species including *Chlamydia pneumoniae, Porphyromonas gingivalis,* and *Propionibacterium acnes* (recently renamed *Cutibacterium acnes*) have also been linked with AD [39–45]. *C. pneumoniae* is an intracellular respiratory bacterial pathogen that was proposed to cause sporadic late-onset AD (reviewed in [40]). *In vitro* studies have shown that *C. pneumoniae* is able to infect human astrocytes and to promote amyloidogenic APP processing [41] and murine models of *C. pneumoniae* CNS infection have recapitulated the cardinal features of AD [42].

The vast majority of such microbial survey studies in AD have relied on molecular diagnostics in which the bacterial DNA is directly detected, either by a PCR-based methods [44,46] or *in situ* hybridization (FISH) [38] – as opposed to cultural methods owing to the demonstrated difficulty in culturing bacteria associated with chronic infections and biofilms [47–52] and the greatly improved sensitivity and specificity of nucleic acid-based methods [53–55]. Most recently, species-specific, pan-domain molecular diagnostics have become available for bacteria [56–60]. These assays provide for unbiased surveys without the need for investigators to *a priori* decide what taxa to survey.

Epidemiological studies have raised a possible association between periodontitis and AD [43,61]. Among the periodontitis-related pathogens, *P. gingivalis* is a keystone pathogen for both chronic periodontitis and systemic sequelae. Dominy et al. [44] have detected *P. gingivalis* DNA and gingipains (arginine or lysine specific cysteine proteases and major virulence factors in *P. gingivalis)* in postmortem AD brains and in cerebrospinal fluid (CSF) of living AD patients.

Moreover, a recent *in vitro* study by Haditsch et al. [45] have shown the neurotoxicity of the gingipains and that *P. gingivalis* can invade and persist in neurons and that the infected neurons display AD-like neuropathology including an increase in tau phosphorylation ratio. Preliminary microbiome studies using next-generation sequencing of the variable regions of 16S ribosomal rRNA gene (V3, V4) have also identified several bacterial species in both AD brains and non-demented controls [61,62]. Emery et al [61] have found higher bacterial loads in AD brains and a higher proportion of Actinobacteria, especially *P. acnes,* whereas the study of Westfall et al [62] showed no difference in bacterial populations between AD and control subjects but variations in microbial composition between hippocampal and cerebellum regions in AD subject’s brains. Microbiome studies have also detected several fungal genera as being more prevalent in AD brains *(Alternaria spp.*, *Botrytis spp.*, *Candida spp.,* and *Malassezia spp.)* [63].

The potential involvement of microbes as etiological agents of AD has been strengthened by the evidence that the Aβ peptide has potent antimicrobial properties. Soscia et al. [64] demonstrated *in vitro* that the Aβ peptide possessed antimicrobial properties. The antimicrobial activity of Aβ is comparable to the well-known human antimicrobial peptide (AMP) LL-37. The protective effect of Aβ against bacterial infection has been shown in a murine model where it was demonstrated to mediate entrapment of microbes by oligomerization and fibrillization of Aβ [65]. The demonstration that Aβ is an AMP has led to the antimicrobial protection hypothesis. In this model, Aβ deposition is a defensive mechanism against infection and AD pathology results from a chronic innate immune inflammatory response to a recalcitrant bacterial biofilm leading to the accumulation of Aβ deposits and ultimately mediating neurodegeneration.

In this study, we take advantage of the Pacific Biosciences (PacBio) long-read DNA sequencing technology we previously developed [59,60,66] to sequence the full-length bacterial 16S rRNA gene and to profile the bacterial communities to the species level in AD-affected and non-demented age-matched brains.

## MATERIALS AND METHODS

### Biological Material and Sequencing

#### Brain tissue samples

Frozen postmortem human brain samples were obtained from the University of Arkansas for Medical Sciences (UAMS). All the samples were neuropathologically evaluated by the provider. All Alzheimer’s disease cases were given Braak stages IV-VI. The control cases designated as age-matched controls (controls) were described as non-demented. The average postmortem interval was 8hr. The data contained 130 samples from 32 individual subjects about half of whom had Alzheimer’s Disease (“AD”). For most subjects, we had at least one sample from the entorhinal cortex and the frontal and temporal lobes. We had no underlying histological information from the sample sites with regard to AD diagnoses. To prevent contamination, the samples were handled in a laminar flow hood with proper personal protective equipment (lab coat, mask, gloves and protective eyewear).

#### DNA extraction

Total DNA was isolated from frozen brain biopsies using the DNeasy Blood & Tissue Kit (Qiagen) according to the manufacturer’s recommendations with slight modifications. Biopsy material was incubated overnight at 56 °C with 570 μl ATL tissue lysis buffer with 30 μl Proteinase K in a Lysing Matrix E tube (MP Biomedicals LLC), homogenized by SPEX 1600 MiniG (SPEX SamplePrep) for 10 min at 1500 Hz, and centrifuged 1 min × 13,000 rpm. DNA was eluted with a 200 μl AE elution buffer. DNA quality and quantity were analyzed by agarose gel electrophoresis and Nanodrop 2000 spectrophotometer (Thermo Fisher Scientific) respectively.

#### Full-length 16S rRNA gene amplification

The taxonomic composition of bacterial communities in the post-mortem human brain tissues were analyzed using the Pacific Biosciences (PacBio) single molecule real-time (SMRT) sequencing technology (Pacific Biosciences, Menlo Park, CA, USA) to obtain the full-length 16S ribosomal RNA (rRNA) gene sequences as previously described [59,60,66]. Briefly, the full-length 16S rRNA gene was amplified using the universal 16S rRNA bacterial primers 27 F (5’-GRAGAGTTTGATYMTGGCTCA) and 1492 R (5’-TACGGYTACCTTGTTACGACTT). Both the forward and reverse 16S primers were tailed with the universal sequences (5’-GCAGTCGAACATGTAGCTGACTCAGGTCAC and 5’ - TGGATCACTTGTGCAAGCATCACATCGTAG, respectively) to allow for multiplexed sequencing and a 5’ block (5’NH2-C6) was added according to the recommendations of Pacific Biosciences. The primers were synthesized and HPLC purified by Integrated DNA Technologies.

Barcoded 16S rRNA amplicons were obtained via a 2-step PCR. All the PCR reactions were performed in 96 -well plates. The first PCR round was performed using 10 μl of total DNA (approximately 1-2 μg of DNA) as template, the universal 16S rRNA bacterial primers 27F and 1492R described above (0.2 μM each), and 1X Hot-Start GoTaq DNA Polymerase Master Mix (Promega) in 50 μl final volume. Cycling conditions were 94 °C, 3 min; then 35 cycles of 94 °C 30 s, 54 °C 30 s, 72 °C 2 min, and a final extension step at 72 °C for 5 min. The amplified products were then analyzed by agarose gel electrophoresis to check the quality and the size of the amplicons. The second PCR round was performed in a 50 μl reaction volume containing 2 μl of a unique primer pair of Barcoded Universal F/R Primers (Pacific Biosciences, 100-466-100), 10 μl of 16S rRNA amplicons from each sample, and 1X Hot-Start GoTaq DNA Polymerase Master Mix (Promega). Cycling conditions were 94 °C, 3 min; then 20 cycles of 94 °C 15 s, 64°C 15 s, 72 °C 2 min, and a final extension step at 72 °C for 5 min. PCR products were cleaned with AxyPrep MagPCR (Axygen) according to the manufacturer’s protocol with a volume ratio (bead suspension to PCR product) of 2:1 and eluted in 50 μl of water. Cleaned barcoded 16S rRNA amplicons were quantified using AccuClear Ultra High Sensitivity dsDNA Quantitation Kit (Biotium) on BioTek™ FLx800™ Microplate Fluorescence Reader. Based on quantification results, barcoded amplicons were then pooled in equimolar concentration into multiplexed sets of 2 to 18 samples per pool.

#### Pacific Biosciences Sequel System sequencing

Sequencing libraries were constructed from each pool of barcoded amplicons using the SMRTbell Express Template Prep 1.0 kit (Pacific Biosciences, 100-259-100) according to the manufacturer’s instructions [67]. Multiple SMRTbell libraries were then multiplex sequenced in a single SMRT Cell 1M on a Pacific Biosciences Sequel System.

#### Generation of demultiplexed CCS reads

The raw subreads generated by the Sequel sequencing run were converted into circular consensus (CCS) reads and demultiplexed using the command-line version of the Pacific Biosciences’ workflow engine pbsmrtpipe (v1.3.3) or pbcromwell (1.2.5) within the SMRT Link v7 or SMRT Link v9 software respectively. Differences were minor. CCS reads were generated using the following parameters: minimum number of passes = 3, minimum predicted accuracy = 0.99, minimum subread length = 1000. CCS reads were then demultiplexed by their barcode into FASTQ files.

#### OTU clustering and taxonomic classification

Full-length 16S (FL16S) sequences were then clustered into Operational Taxonomic Units (OTU) and assigned to species level taxonomic classification using the Microbiome Classification by Single Molecule Real-time Sequencing (MCSMRT) pipeline designed by Earl et al. [59]. The MCSMRT pipeline was specifically built to: (i) process PacBio CCS reads, (ii) de novo cluster high-quality FL16S sequences into OTUs, (iii) taxonomically classify each read and assign confidence values at each taxonomic level, and (iv) quantify the abundance of each OTU. Only the CCS reads (hereafter reads) with a number of passes greater than 5 were clustered into OTUs using a 3% centroid-based divergence level.

### Analytical Methodologies

#### Introductory Comments

##### Analysis Models

Our focus for the data analyses was to find one or more of the following patterns in the data: (1) individual microbes which were either correlated or anticorrelated with AD, or (2) combinations of microbes that were correlated or anti-correlated with AD, given the number of bacteria observed. In other words, we were not just interested in whether a bacterium was intrinsically pathogenic, but also whether pathogenicity derived from a polymicrobial interaction of two or more microbes acting within ecosystems. We understand that microbes other than bacteria could be involved but did not attempt to observe fungi, viruses, or other microorganisms. That will be the subject of future work.

##### Data Challenges: Lack of Functional and Temporal Information

Our objective was to determine if the bacteria we detected suggest a causal relationship with AD and attempt to construct a bacteria-based etiology of the disease. Our experiments, however, only identified the presence and abundance of bacteria, and presence is not a surrogate for functionality. Further, all of our measurements are at a particular point in time, after death, making it difficult to explore the time dependence of the illness without additional assumptions about the nature of the data.

In addition to these challenges, our sequencing technique created the possibility that our experiments generated more information than was needed. First, the sequencing sensitivity enabled the detection of large numbers of taxa, some of which could have similar behavior and function. In other words, the dimensionality of the data was potentially too high. Second, the abundance resolution in each bacterial dimension may not be relevant for either individual bacterial or ecosystem behavior. For example, we do not know if having 10% of a particular bacterium in a sample is any different than 40% in terms of pathogenicity. The latter view is suggested by the population abundance standard deviations of many orders of magnitude for samples from the Alzheimer’s and control subjects, as well as the large overlap between the abundance distributions for particular bacteria. In earlier unpublished work, we found that reducing the abundance resolution with a logarithmic binning was a useful way to begin to explore patterns in data both with simple and rigorous methods without filtering out effects from small abundances.

Reducing the dimensionality of the data requires finding taxa with similar or equivalent behavior. The literature, however, provided limited functional information for analysis at any level of the phylogenetic hierarchy. The literature does not provide any guidance on biologically relevant abundances either.

To deal with the lack of explicit temporal information in the data, we made assumptions about the evolution of the disease. Knowing that AD develops over a long time, we surmised that the brains of AD subjects contained bacterial infections from various evolutionary stages of the disease and that our samples therefore may have selected infections from a multiplicity of stages of the disease. We did not know *a priori* if our sample size mixed infections from multiple stages of the disease.

##### Spatial sampling

It is well known that Alzheimer’s disease affects multiple regions of the brain which are manifested by large-scale morphological changes. We assumed that the disease effect was homogeneous over areas that are large compared to the sample size (about 1 mm across). Therefore, we hoped that a small number of samples might represent the disease’s effect on the brain. In the future, a larger number of samples should help establish the validity of this under-sampling justification. Even so, while it would appear that this decision limited the spatial resolution of the experiment, we found ways to combine the samples to estimate the large-scale spatial distribution of our bacterial measurements.

Further, we know that there are bacterial communities that are organized and function on scales smaller than our sample size. These bacteria may form complex ecosystems of interacting species and these ecosystems may interact with, or at least affect, spatially adjacent but differentiated ecosystems. The differences in these ecosystems may be defined by niches that provide specific nutrients or other environmental factors conducive to their growth.

As our physical sampling scale is tens to hundreds of times larger than the smallest (cellular) scale, the sample abundances we measured averaged out the details of the densities of ecosystems and their microscopic distributions at the cellular scale. Here too, our methodologies permitted us to find patterns in the spatial distribution of ecosystems at the cellular scale.

The methodologies described below also allow us to speculate on how the bacteria enter the brain. It may be possible to distinguish, for example, whether they enter through a limited number of failures in the blood brain barrier and subsequently multiply and spread or whether they enter from numerous random failures over a large region accompanied by a minimum of spreading.

##### Two Analytical Approaches

We took two different analytical approaches to look for single species and multi-species microbial relationships with AD. These methods were alike in some ways, but they also had a major difference. Thus, any similar findings between the two methods would provide mutual orthologous support for their respective findings and would serve as strong evidence that the findings were inherently reproducible.

To find relationships among individual bacterial species and AD, we first investigated the differences in relative abundance of individual taxa between AD and control groups. We used a hierarchical Bayesian modelling approach based on a Dirichlet-multinomial model (DMM). This procedure was supervised as it used information about whether or not the samples came from AD subjects.

To find relationships between combinations of bacterial species and AD, we used an approach described below [68] that first found relationships among the bacteria without using information about the disease state of the subject at all, i.e., it was unsupervised. This approach enabled us to group the bacteria and their abundances into different classes and then relate the classes to AD.

We used the classes derived from the unsupervised step to infer the existence and approximate microscopic spatial distribution of distinct ecosystems at the human cellular scale. Further, we estimated the macroscopic spatial distribution of mixtures of these classes on a brain lobe scale.

To understand the origins of possible pathogenicity, we needed to also include the disease state of the subject. In each approach, we had to be careful to treat sample and subject differently in order not to assume the bacteria in a particular sample are pathogenic just because they originate from a subject with AD. Each analytical approach handled this challenge differently. The first approach distinguished sample from subject in its statistical model. The second approach used the bacterial classes computed for each sample of a subject to find relationships between bacterial classes and AD in a second level of analysis.

Last, to understand the etiology of the disease, we used the commonalities in bacteria between classes of samples to order them in time from earlier classes that were not correlated with AD to later ones that were.

In summary, overall, our analysis operates on four spatial scales and one time scale. There is the cellular ecosystem scale, which is not directly observed, the sample scale, the brain region/location scale which is under-sampled (several samples per brain), and the subject scale which provides disease information. The temporal scale relates to disease stage.

##### Statistical Properties of the Data

We present some simple statistical results below to help to suggest why we chose particular analytical approaches. These simple statistics do not answer our scientific questions but emphasize where the challenges are.

Table 1 shows the top 30 genera ordered by their abundances in the data set. The list contains both species and genera where we have broken out species for several high abundance genera. In this paper, we will often refer to *Propionibacterium*, *Acinetobacter*, and *Comamonas* as the principal bacteria mainly because of their overall abundance and prevalence, but in the case of *Comamonas* because of its abundance and prevalence within a single class not associated with AD.

**Table 1:**
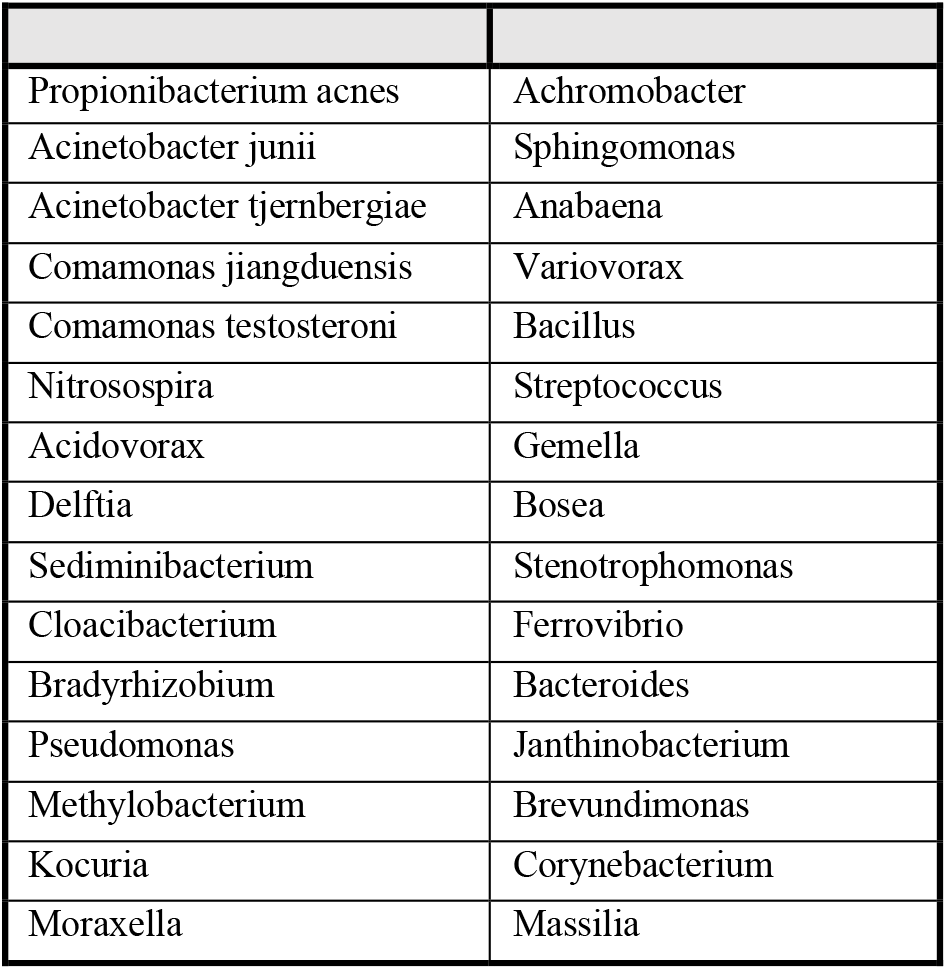
Top 30 Genera/Species by prevalence in order top to bottom, left to right.

We show a couple of comparisons of the abundance distributions for two of these in AD and control samples in Figure 1. While there is a hint of difference in the average abundances between the AD samples and the controls, the wide variances apparent in the figure render the differences statistically insignificant. This pattern is similar for all of the bacterial species that occur frequently in the samples.

**Figure 1:**
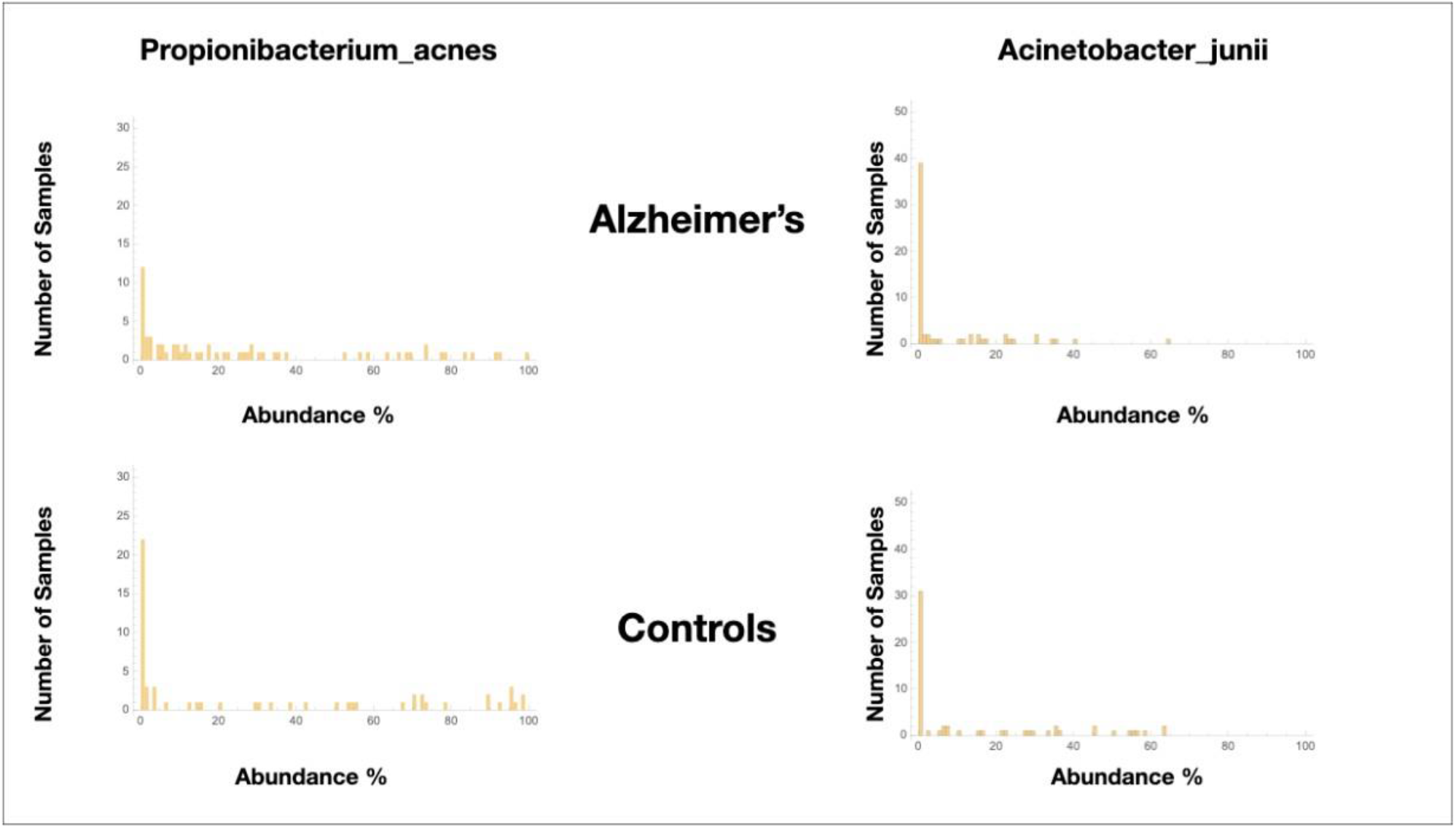
Comparisons of AD & control abundance distributions for Propionibacterium acnes and Acinetobacter junii.

Another typical characteristic that we observed was the sparseness of the data, meaning that most of the observed bacteria do not occur in most of the samples and if they do, they do not have the same abundance. This could mean that the bacteria have little to do with AD or that functional redundancies across bacteria must be discovered to reveal bacterial pathogenicity.

A number of bacteria have high abundances only in a few samples, e.g.,*Methylobacterium.* Using standard arguments, we could have chosen to filter these out because of their low occurrence but it is hard to dismiss these bacteria because they have high abundance and, generally speaking, high abundance is more likely causal than low abundance. We considered that these were contaminants but eventually found that together they exhibited patterns that could be a critical factor in the etiology of AD.

##### Discrete Statistics of the Data

We described above various challenges in analyzing this data set. In order to get a better sense for the data and potential biologically meaningful patterns it harbored, we decided to generate a view of the data with greatly reduced abundance resolution. Mindful of the possibility that some bacteria of low abundance may have a disproportionate effect on pathogenicity, we chose to logarithmically bin the data. This is not meant to be a suggestion that this is the best way to analyze the data. It is a starting point to begin to get a sense of it.

Specifically, we defined a set of contiguous abundance bins in the 0.0% to 100.0% range and labeled them with integers. The bin sizes which we chose are shown in Table 2. We then mapped the abundance data into descriptive discrete objects formed by appending the numerical bin label to the microbe name. The result of the binning was to transform a row of abundance data from a table whose rows correspond to samples and whose columns correspond to microbe name into a list of microbial objects.

**Table 2:**
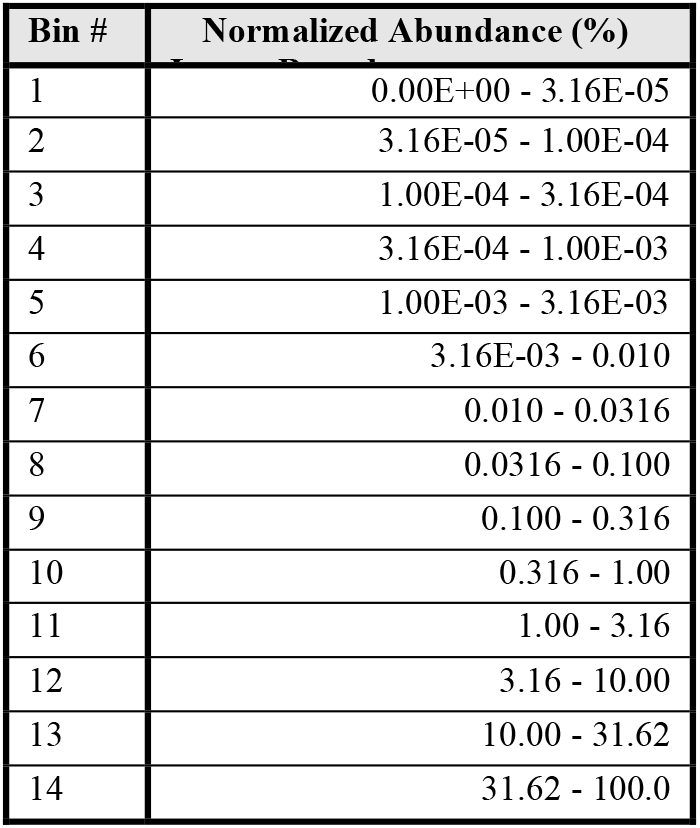
Abundance bins in percent.

| Bin # | Normalized Abundance (%) |
| --- | --- |
| 1 | 0.00E+00 - 3.16E-05 |
| 2 | 3.16E-05 - 1.00E-04 |
| 3 | 1.00E-04 - 3.16E-04 |
| 4 | 3.16E-04 - 1.00E-03 |
| 5 | 1.00E-03 - 3.16E-03 |
| 6 | 3.16E-03 - 0.010 |
| 7 | 0.010 - 0.0316 |
| 8 | 0.0316 - 0.100 |
| 9 | 0.100 - 0.316 |
| 10 | 0.316 - 1.00 |
| 11 | 1.00 - 3.16 |
| 12 | 3.16 - 10.00 |
| 13 | 10.00 - 31.62 |
| 14 | 31.62 - 100.0 |

This scheme allowed us to count the number of times an object was found among the AD samples compared to the controls. Later, we will describe how we used this as the initial input data for one of our algorithms which was, in fact, the initial motivation for transforming the data in this manner. The bin width was somewhat arbitrary and somewhat guided by experience in microbiome analysis and was eventually amended for use with our algorithm.

We compiled object statistics in Table 3 where we compared object occurrence in samples from subjects with and without AD. Note the correlations with AD among certain objects, in particular *Propionibacterium acnes-13, Acinetobacter junii-* 13 and *Acinetobacter tjernbergiae-13.* It is important to note that these are not the maximum abundances nor close to 100% abundance, indicating the presence of other microbes in the sample. These are clues to the abundance dynamics that will be more fully explored below. *Comamonas jiangduensis* presents a curious situation in that its objects are associated with the no-AD disease state of the controls.

**Table 3:** Object statistics, comparison of samples from Alzheimer’s and controls subjects.

| MICROBIAL OBJECT | Alzheimer's | Controls | MICROBIAL OBJECT | Alzheimer's | Controls |
| --- | --- | --- | --- | --- | --- |
| Propionibacterium acnes-14 | 20 | 23 | Acidovorax-8 | 4 | 3 |
| Propionibacterium acnes-13 | 17 | 6 | Acinetobacter junii-10 | 2 | 5 |
| Acinetobacter junii-13 | 14 | 8 | Comamonas jiangduensis-14 | 0 | 7 |
| Acinetobacter tjernbergiae-13 | 11 | 6 | Sediminibacterium-13 | 4 | 2 |
| Acinetobacter junii-14 | 4 | 13 | Pseudomonas-9 | 4 | 2 |
| Cloacibacterium-12 | 9 | 6 | Comamonas testosteroni-11 | 4 | 2 |
| Propionibacterium acnes-12 | 9 | 4 | Sediminibacterium-11 | 3 | 3 |
| Acidovorax-13 | 8 | 4 | Streptococcus-12 | 2 | 4 |
| Acinetobacter tjernbergiae-10 | 1 | 10 | Nitrosospira-13 | 4 | 1 |
| Cloacibacterium-11 | 3 | 7 | Moraxella-10 | 4 | 1 |
| Acidovorax-11 | 3 | 7 | Delftia-14 | 4 | 1 |
| Propionibacterium acnes-11 | 6 | 3 | Acinetobacter junii-11 | 4 | 1 |
| Acinetobacter tjernbergiae-12 | 6 | 3 | Acidovorax-14 | 4 | 1 |
| Acinetobacter tjernbergiae-14 | 5 | 4 | Sediminibacterium-10 | 3 | 2 |
| Streptococcus-11 | 4 | 5 | Nitrosospira-14 | 3 | 2 |
| Sediminibacterium-12 | 6 | 2 | Bacillus-10 | 3 | 2 |
| Corynebacterium-11 | 6 | 2 | Acidovorax-12 | 3 | 2 |
| Comamonas testosteroni-10 | 6 | 2 | Streptococcus-10 | 2 | 3 |
| Delftia-11 | 5 | 3 | Stenotrophomonas-10 | 2 | 3 |
| Moraxella-11 | 3 | 5 | Propionibacterium acnes-10 | 2 | 3 |
| Corynebacterium-10 | 3 | 5 | Kocuria-10 | 2 | 3 |
| Bradyrhizobium-10 | 3 | 5 | Streptococcus-9 | 1 | 4 |
| Acinetobacter junii-12 | 3 | 5 | Pseudomonas-8 | 1 | 4 |
| Pseudomonas-12 | 2 | 6 | Pseudomonas-11 | 1 | 4 |
| Cloacibacterium-10 | 2 | 6 | Nitrosospira-11 | 1 | 4 |
| Novosphingobium-9 | 5 | 2 | Lactobacillus-10 | 1 | 4 |
| Moraxella-12 | 5 | 2 | Anaerococcus-8 | 1 | 4 |
| Moraxella-9 | 4 | 3 | Comamonas jiangduensis-13 | 0 | 5 |

Last, while the statistics are weak, there is little indication that low-abundance microbial objects are correlated with disease state.

##### A note of caution

The statistics quoted in the previous paragraph and in Table 3 show how often the object came from a subject who had or did not have AD. When an object occurs more often in AD subjects, this does not necessarily mean that the bacterium and abundance it represents is pathogenic. We will show below that many of these are likely not pathogenic.

#### Method for Individual Bacteria

In this section, we describe ways to explore differences in individual bacterial abundances between AD and control subjects.

##### Data filtering

As samples vary in total read number and low-yield, samples could be noisy. The samples with less than 100 total reads were removed from the dataset. Four blank extraction controls (no sample input) were processed in the same way as the true biological samples to allow identification of any contamination from reagents or during sample processing. Potential contaminant OTUs were detected based on their occurrence in biological samples vs. negative controls using a prevalence-based method (IsNotContaminant function) from the R package Decontam [69]. To qualify as contaminant, an OTU had to have a score ≥ 0.5 or a higher mean relative abundance in the negative controls than the biological samples (Supplementary Table S1). Contaminant OTUs were then removed from the dataset. The phyloseq R package [70] was used for handling OTU counts, taxonomy and sample metadata.

##### Exploratory data analyses

OTU counts were normalized using the centered log-ratio (clr) transformation to account for the compositional structure of the data [71]. Given an observation vector of D OTUs in a sample, X = [x1, x2, …, xD], the clr transformation for the sample is calculated as follows:

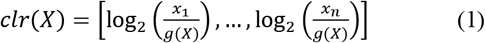

where

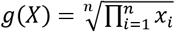

is the geometric mean of X. A pseudocount of 1 was applied to zero entries in the OTU count table before taking the log to the base 2. A positive clr value for a given OTU indicates a relatively higher amount than the overall composition mean and a negative value indicates a relatively lower amount. A principal component analysis (PCA) of the clr-transformed data was performed.

##### Differences in relative abundances

To test for differences in the relative abundances of individual OTUs between AD and control sampling groups, the OTU count data were analyzed using a hierarchical Bayesian model based on the Dirichlet and multinomial distributions as described in Harrison et al. (2020) [72]. The Dirichlet-multinomial model (DMM) is relevant for the compositional structure of microbiome data. This model allows the sharing of information (parameters) among the samples within sampling groups and the estimation of the relative abundance of each OTU while propagating the uncertainty in those estimates [72–74].

DMM estimates the multinomial parameters that describe the proportion of each OTU in a sample (vector 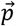) and the Dirichlet parameters that describe the proportion estimates of each OTU for the entire sampling group (vector 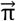).

To take into account that some samples were from the same individual subject (non-independent samples), the vector 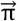 is informed by an additional Dirichlet distribution with 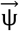 that describes the relative abundance of OTUs within each subject and the intensity parameter τ.

The model was specified as follows:

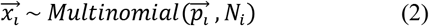

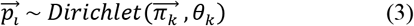

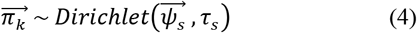

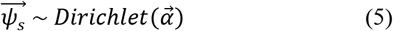

with priors,

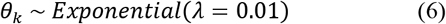

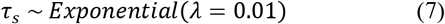

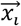 is the observed count of a particular OTU in the sample *i*. 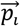 is the vector of the estimated OTU proportion and *N_i_* is the total counts in each sample *i*. 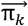 is the estimated proportion of each OTU in the sampling group k and θ_*k*_ is the intensity parameter of the Dirichlet distribution. 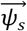 describes the expected relative abundance of OTU within each subject *s* and *τ_s_* is the intensity parameter of the Dirichlet distribution. The prior for the 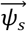 parameter is a Dirichlet distribution with equal prior probability for each OTU. Exponential distributions have been used as the prior for θ_*k*_ and *τ_s_*.

To quantify the differences in relative abundance between AD and control groups, the posterior probability distribution (PPD) for the OTU of interest in the control group was subtracted from the PPD of that OTU in the AD group. Following convention, if 95% of the PPD for difference does not overlap zero, there is high certainty that the OTU of interest differs in relative abundance between AD and control.

DMM was specified in the Stan probabilistic programming language through the Python interface Pystan (version 2.19.1.1) that implements the Hamiltonian Monte Carlo No-U-Turn sampler (HMC-NUTS) algorithm. For each of four chains, 3500 iterations were used with 1500 burn-in and a total of 4000 samples were drawn (thin=2). Convergence was assessed using the Gelman-Rubin statistic.

#### Methodfor Combinations of Bacteria

##### Development of New Methods

In the following paragraphs, we describe how we identified sets of microbes that may be related to Alzheimer’s disease using Latent Dirichlet Allocation (LDA). We enhanced the standard LDA approach in various ways and used graph theoretic techniques for optimization and visualization of the results.

We also tried to analyze the data using Principal Component Analysis (PCA) but found it wanting for various reasons discussed below. A detailed examination and comparison of this and other methods is beyond the scope of this work.

##### Multi-scale analysis

The algorithm operates entirely at the sample scale as the microbial objects that are its input are sample scale entities. We will explain later how the statistics of the results of computations with this algorithm can be used to infer information about the spatial distribution of the bacteria at the cellular scale and brain scale and how their abundances evolve in time. All of this will lead to a simple theory of the etiology of Alzheimer’s disease within a bacterial model.

##### Summary of Latent Dirichlet Allocation (LDA)

There are, of course, many ways to reduce dimensionality but it is important to use a technique that has the capability of leading to biologically meaningful conclusions. We decided to adapt a technique from computational linguistics that can group words into meaningful topics and to summarize documents by these topics so that they are easily discerned by human readers. LDA [75] does this by classifying words, i.e. assigning classes (topics in the literature) to each word. Documents are statistically summarized by the fraction of their words assigned to each class. Words are summarized by the fraction of times they are assigned to each class in the entire set of documents. Classes (topics) are summarized by the fraction of times each word in the set of documents is assigned to a class. The object and class summaries are how LDA reveals that a particular word may have more than one meaning. Class statistics are also how LDA reveals how a set of words could have a common meaning by forming a topic.

So, we reasoned that LDA might be able to group bacterial abundances in a way that reveals biological meaning. To adapt this approach for microbial data, we needed to find a suitable alternative to the words of the linguistic analysis. We chose to define a sample’s words by the objects we introduced earlier, namely a concatenation of microbe name and the abundance bin its measurement fell within. This has the value of imbuing the object with information about both its microbial function (i,e, its name) and its relative abundance in a sample, both of which might be needed to reveal biological meaning.

These microbial objects are then assigned one of a preset number of classes using the LDA algorithm which we describe at a high level in the following paragraphs and in Figure 2. Dimensionality reduction occurs because the number of classes is much less than the number of unique microbial objects. The result describes each sample by the fraction of its objects that are assigned to each class, termed the sample distribution. Objects are described by the fraction of their assignments to each class for the whole data set, which is called the object distribution. Each instance of these distributions is a vector whose number of components is equal to the number of classes chosen. Classes can be summarized by the fraction of times each object is assigned to a class. When an object distribution has more than one non-zero component, it suggests that it has multiple meanings. When many objects occur frequently in one class, it suggests a relationship, common meaning, or similar functionality among the objects.

**Figure 2:**
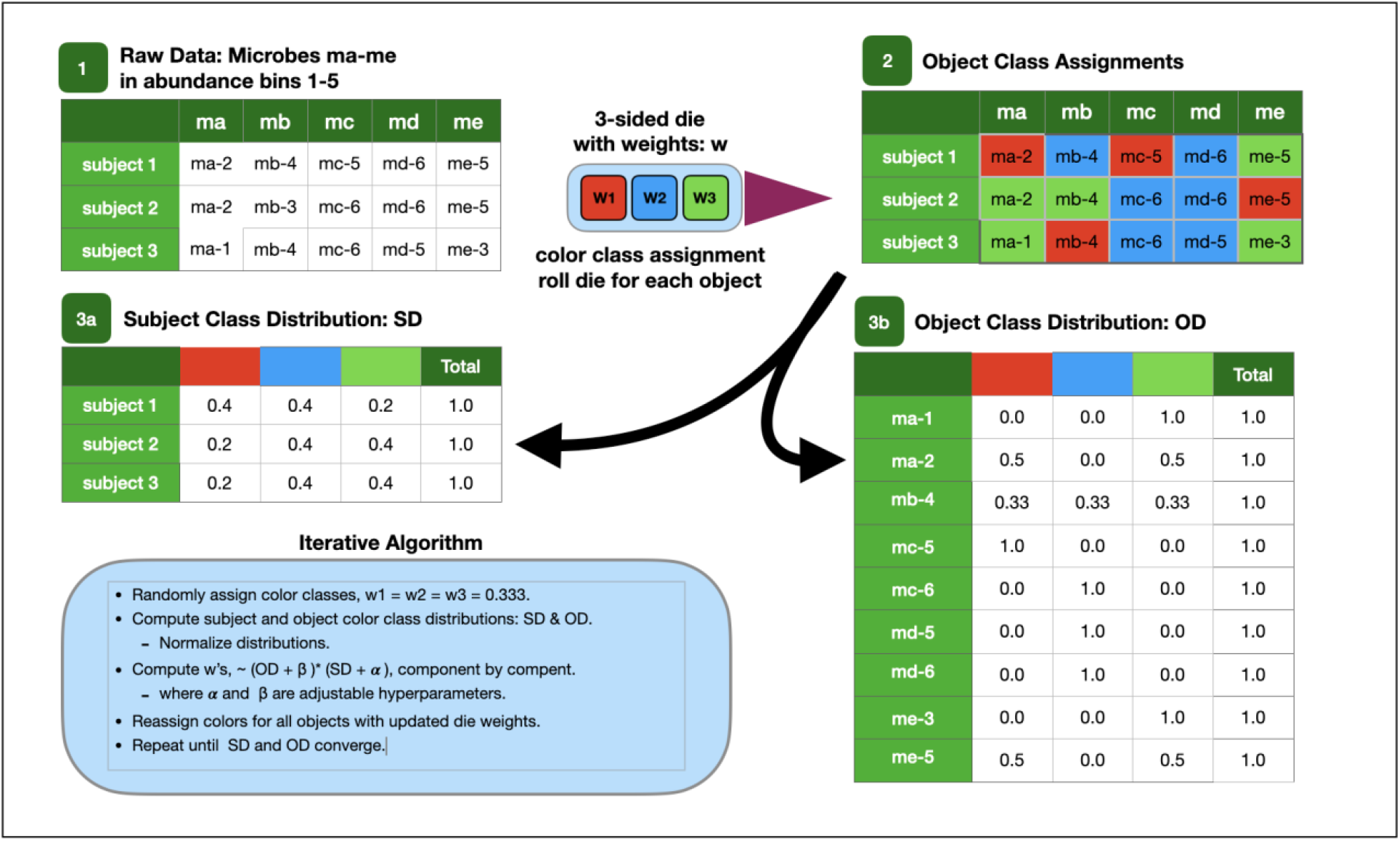
Cartoon of LDA Algorithm.

The results data can be tabulated in two tables. The sample distribution table has a row for each sample and its columns are labeled by class. Its entries are the number of times the sample’s objects were classified with a particular class. The rows can be normalized to show the fraction of objects classified in a class. The object distribution table has a row for each object with columns labeled by class. One microbe may have multiple rows corresponding to different abundance bins. The entries are the number of times objects were classified with a particular class for the entire data set. The object class distribution is computed by normalizing across the rows to show the fraction of times each object is assigned to a class. If the same count data were normalized by column instead, the composition of each class is revealed by microbial object. This is a rigorous definition of the class microbiome for the sample. Sometimes we use approximations by identifying microbial objects from samples with a common maximum class component. It should always be remembered that all of these distributions are computed from the same counts of class assignments of the objects.

In order to implement this approach, we must compute the class of an object because objects are measured but class is computed. LDA assumes that the class of an object is determined first by the chance that each class is present in the sample and second, for each class, what the chances are of measuring a particular object. In other words, the class of an object is determined by two probability distributions. The former distribution is given by the sample class distribution and the latter by the object class distribution. The probability of the class of the object is roughly the component-by-component product of both distributions. Unfortunately, we do not know these distributions *a priori*. In other words, this is one of those situations where we need the answer in order to compute it.

One way to solve this problem is through iteration. We chose a Markov chain Monte Carlo (stochastic) approach [68]. The computation begins by randomly assigning a class to each object of each sample. This allows starting sample and object distributions to be calculated with their normalized values interpreted as probabilities for the stochastic computation.

For each object, we recompute the two probability distributions, multiply them component by component and renormalize. The components form the weights of an unfair die which is used as a random number generator whose probabilities contain complete information of the object and sample. When the die is rolled, it yields a class which is used to update the class assignment of the object. This sequence of stochastic computations forms a Markov chain since each depends on the last. The process iterates through all the data multiple times until the two distributions stop changing significantly, i.e. until they converge. The result is hopefully a stable and repeatable class distribution for each sample class distribution (SD) and for each object class distribution (OD).

There is one other way to look at what we are doing that helps to distinguish this method from other dimensionality reduction approaches. We note that one of the distributions, the sample distribution, is computed only from the sample while the other, the object distribution, is computed from all samples. We can call the sample distribution a local distribution because it only uses class assignments from the sample and the object distribution a global distribution because it uses class assignments from all the samples. Using this terminology, we see that the class of each object is determined by both a local and global contribution. As long as the experiment was adequately designed so that local and global information is adequately sampled, the scheme can work. Other dimensionality reduction schemes, e.g. t-SNE and UMAP [76,77] tend to focus on sample attributes, e.g. making sure that sample closeness in the n-dimensional input space is preserved in the lower dimensional output space. Local schemes like these may have trouble allowing objects (or measurements) to have multiple meanings although the common objects within the samples of a sample cluster in low dimensions could represent common meaning the way a class does in LDA. The rigorous definition of cluster may involve a second computational step or often, eyeballing. In contrast, the LDA cluster is intrinsically defined by class.

LDA finds co-occurrences within samples that suggest something going on in one sample is going on in another. This is its local capability, but it goes further because of the global contribution. For example, if microbe A occurs with a microbe in set B in one sample and microbe A occurs with another microbe in set B in another sample, then LDA finds those patterns too by classifying the samples in similar ways. The latter suggests that samples such as these might be evidencing a common functionality.

A heuristic derivation of the complete classification formulas is shown below that ties to the rigorous Bayesian derivations in the literature [68,75,78–84]. We also describe several extensions that collectively we call MLDA that enhance repeatability.

Last, the most important idea to retain about this method is this. If a class structure can in fact be found, ignoring it is tantamount to averaging over it. Consequently, this implicit average risks averaging out the very evidence that is sought. It is equivalent to ignoring confounding variables in an analysis. This is an issue in the individual bacteria method above that will be further discussed.

The patterns found by LDA are sometimes difficult to understand so we developed graphical visualization techniques to assist us. These are also used extensively to refine algorithms, check for convergence and repeatability and to optimize adjustable parameters.

##### Type I Graphs

This type of graph, where the nodes are samples, was designed to display classification results, sample similarity, metadata values and metadata statistics. A glance enables you to get a sense of the quality of the classification and see the presence of statistical fluctuations in the classification. The graph helps to reveal gross features of the classification which may relate to the emergent features of the ecosystem biology. The graphs were drawn using Wolfram Mathematica [85,86].

###### Nodes

Each node is a sample.

###### Color

The LDA computations result in each sample being described by C components, where C sis the preset number of classes the LDA algorithm used. Each component is labeled by a color. A node’s color corresponds to the component that is the MAXIMUM of the sample’s components. From here on, when we refer to: a color class or the color of a sample or object, we are referring to the maximum component of the distribution. The color of a node should not be confused with an exclusive classification for the node. While each node is, in fact, described by a mixture of C components, the ubiquitous existence of color clusters suggests that the exclusive classification suggested by the colors is an approximation that is justified. We use the concept of color to approximate microbiomes. Here it can be thought of as the set of objects that occur in samples of a given color.

###### Node Size

Nodes were enlarged (other graphs below) if a sample contained one or more specific microbial objects of interest. This visualization is used frequently to explore the class location of objects of the same microbe but differing abundance bin.

###### Node Shape

The shape of the node displays the SUBJECT metadata value - diamonds for AD, circles for controls. Typically, we may note the diamond fraction statistic next to a color cluster. This AD statistic is the number of diamonds in the cluster divided by the total number of nodes in the color cluster. In our data, we have roughly 50% of the samples from AD subjects and 50% from controls. So, if the class means something for AD, the diamond statistic should be way over 50% if there is a correlation with AD or way less than 50% if the class is anti-correlated with AD. The fact that this is not the case is something we address.

###### Edge

Edges were defined by node pair similarity. In general, many types of similarities can be used but we used a coarse measure, the dot product. In this case, the similarity is the product of each pair of components summed together. To define the edges in a graph, we used a similarity range that contained the highest values of node pair similarities because these nodes were the most alike. Forming a histogram of all possible similarities from pairs of nodes, this range would correspond to the right-hand tail of the histogram. We found that even when ranges spanned a small piece of this tail, the entire set of nodes was likely to be included among the selected edges.

###### Node Position

The features above define the topology of the graph — how the nodes were connected [85]. An embedding algorithm is used to position the nodes in 2D, or 3D space. The algorithm finds the equilibrium position of the nodes when the nodes and edges are given physical properties that both repel and attract the nodes. The repulsion is computed by assuming that each node possesses the same electrical charge, and the attraction derives from representing each edge as a spring. This algorithm is known as spring-electrical embedding [86] and the resultant graphs are called force-directed graphs. It is possible to have springs whose spring constants are a function of similarity; however, we used a simple binary method. If nodes were connected, they used springs with the same constant, an adjustable parameter. Node clustering is driven by the edge spring. This algorithm positions the nodes in 3D space and the images we present are a projection of the 3D arrangement onto a 2D plane. Because nodes that are the most similar are connected by springs, samples that are the most similar are pulled together in clusters.

###### Outliers

Typically, nodes that are relatively far away from a cluster compared to other nodes of the same color indicate a statistical fluctuation in the LDA class assignment. The underlying class distribution has a maximum close to another component which should be the maximum, so it ends up with the ‘wrong color’ and because it is not similar to the other nodes, it is positioned far away.

###### Class Number Optimization

The embedding algorithm helped to optimize the class number input parameter. From experience, we knew that microbiome data formed homogeneously colored clusters mainly because samples tend to be dominated by one class. If the class number is set to high, the graph will display small satellite clusters near the main clusters, often not tightly clustered or repeatable from run to run. If it is set too low, clusters will be formed with samples that are too dissimilar. These are often not tightly clustered and can be multi-colored when two smaller components are merged that sometimes become the dominant class. Proving that tightly clustered, homogeneously colored clusters represent the best classification is beyond the scope of this paper but we will assume it since it helps to make sure that samples that have similar composition end up in the same class. Keep in mind that the disease state is not used in the classification computation. Heterogeneous colored clusters suggest a lack of class dominance within samples and a lack of class structure. Small, rarefied satellite clusters of samples are prone to class hopping between runs and therefore suggest diminished repeatability.

##### Type II Graphs

The type II graph is the dual of the sample graph [85,86]. It contains the same object class assignment information as the sample graph. These graphs utilize the classification statistics of microbial objects where the objects are nodes as opposed to the samples of type I graphs. Color is assigned in the same way as in the type I graph but with the maximum class in the object class distribution. The nodes are also positioned using the right tail of the similarity distribution using dot products between object distribution pairs.

Node shape is not used since object nodes do not have a unique disease state. This is because objects contain information from the entire data set including both AD and control subjects. Further, a node’s color does not mean that it only occurs in samples of that color. For example, the A-13 node is in the orange class but there are many instances of A-13 being assigned to the green class. The type II graph is a visualization of the microbiome where an object’s peak component is labeled by color and its similarity to other objects is shown by how proximate it is to other objects.

##### Parameter Setting and Optimization

It will become clear that there are many adjustable parameters in the analytical methodology we used for combinations of bacteria. These include LDA input parameters (e.g. number of classes), Monte Carlo parameters (e.g. number of iterations), parameters that describe the LDA results in graph visualizations (e.g. spring constant), data abundance binning and phylogenetic level summation(naming). This constitutes a large parameter trade space for which we do not have a rigorous optimization methodology. There are several criteria we used to guide our choice of parameters. The last set is treated in more detail in the following section.

###### 1) Convergence of the sample and object class distributions

After some number of iterations in the LDA Monte Carlo computation, we require individual sample class distributions and object distributions to change little after a preset number of iterations. See **Figure 5** and Convergence Monitoring below.

**Figure 3:**
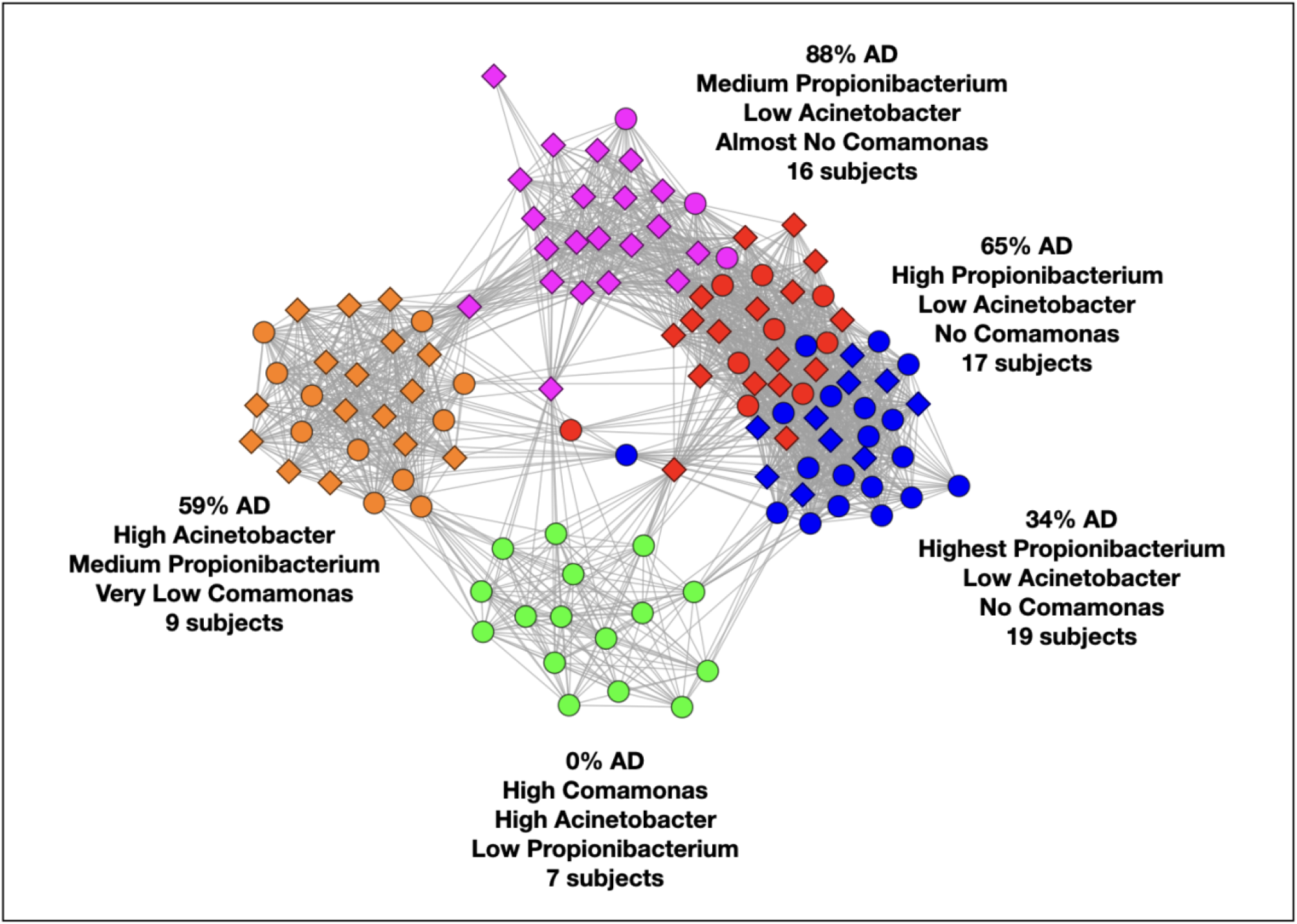
Type I graph. Results from summation of 5 runs. Nodes are samples. Colors are maximum classes. Principal bacterial genera and abundance levels indicated for each color.

**Figure 4:**
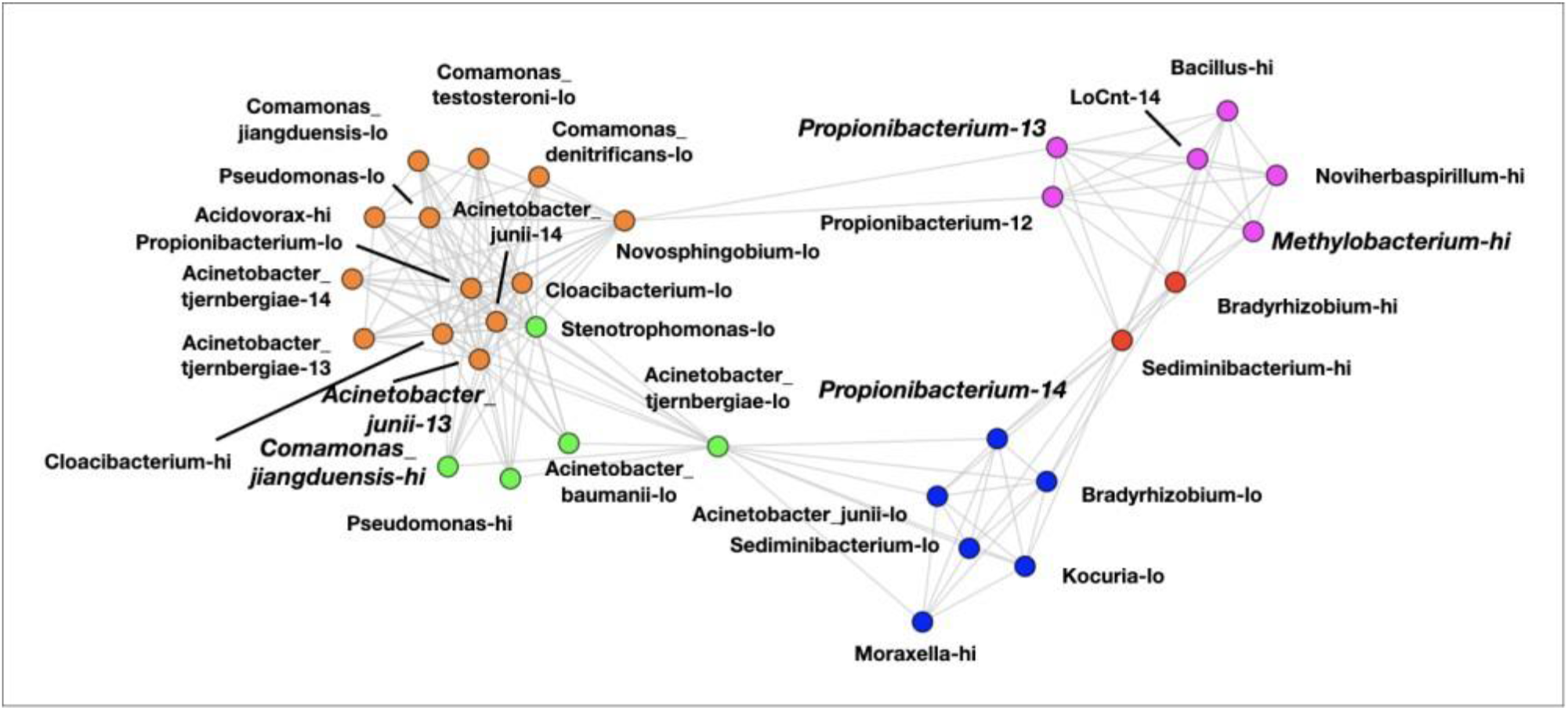
Type II Graph displaying objects with entropies ≤ to 0.7. Example from one run.

**Figure 5:**
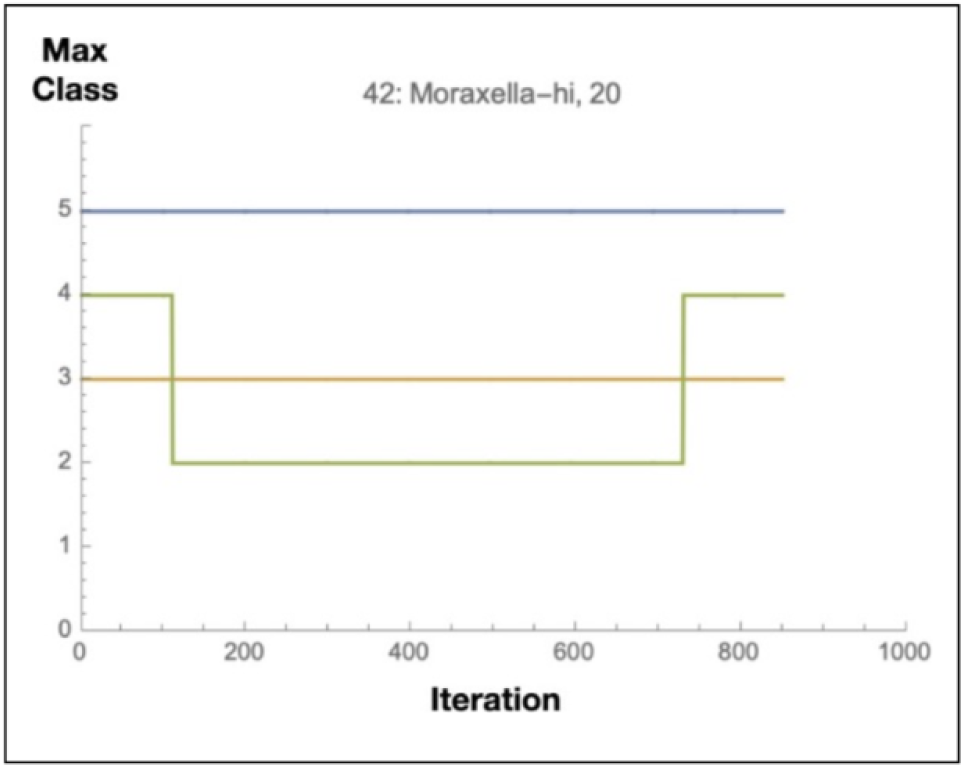
Convergence monitoring.

###### 2) Repeatability

We require that the same samples end up with the same color classes defined above after repeated runs. This is critical because it is possible that there could be convergence without repeatability. A rigorous repeatability evaluation method is detailed below.

###### 3) Sample and Object Graph Quality

The graphs, described above, are a holistic comparison of every pair of samples or objects. The abiding feature of the graphs is that similarly colored nodes cluster together which randomly generated data does not do. In unpublished results of about 7,000 subject’s microbiomes, we saw this pattern ubiquitously. Without proof we used this feature to adjust parameters to get the tightest well -defined homogeneously colored clusters. It seems reasonable to suppose that tight, homogeneously colored graphs represent repeatable computations because the opposite will have ill -defined (multi-colored) and small features that are prone to statistical fluctuations and thus lack of repeatability.

Even with non-optimized parameters, LDA can provide insights about the data, so we explored the parameter space by running the LDA algorithm hundreds of times both to optimize the parameters and get a sense for patterns in the data. We focused on whether objects and samples had repeatable class component maxima (colors) or whether component values were roughly equal. If class means something, the former occurs while if the classification is independent of class, the latter occurs. This is particularly important in the binning and naming adjustments of the next section.

It is important to realize that we are using these techniques to find qualitative patterns in the data leading to insights and the formation of new hypotheses about the biology of AD. Consequently, rigorous optimization and justification of particular parameter values was not attempted, relying instead on a trial-and-error approach and the general criteria above.

Overall, once the criteria were met, we did not continue optimizing parameters, but froze them and tried to discern if the sample and object classification patterns revealed underlying biology. Further validation of this method is important, but we emphasize that there is already an extensive literature on LDA’s ability to find classes (topics in the literature) in documents by finding words that co-occur within documents. Of course, LDA can’t tell the difference between a microbial object and a word.

##### Abundance Binning, Microbe Naming and Object Merging

To utilize the LDA algorithm, the data needed to be converted to objects. This required a phylogenetic level (naming) to sum the OTU data to and an abundance binning structure. The raw data consists of OTU counts labeled by subject, sample, and bacterium. We removed measured contaminants which is described in a subsequent section. The *Propionibacterium* and *Acinetobacter* genera occurred the most frequently in the samples, so we summed these to the species level. Although not as prevalent, we also decided to do the same for *Comamonas* because of its prevalence within the controls. The rest were summed to the genus level. These counts were then normalized to relative abundances within each sample. We did experiment with all genus level summation without much difference in results. The latter three genera are hereafter referred to as the principal bacteria and are sometimes abbreviated, P, A and C.

We binned these counts using the logarithmic binning (14 bins from 10^-5^ to 100%) shown earlier implicitly avoiding the assumption that small abundances were not important. While we did not know if this binning was optimal, we knew from experience that it could reveal microbiome structure, so we began here with the intention of adjusting it to maximize the repeatability of the computations. Discrete measurement objects for each sample were created by concatenating microbe name and bin number (1-14), e.g. *Methylobacterium*-14, as already described. Note that the number of sample objects are not necessarily the same for each sample, nor do we include zero abundance objects.

With this scheme, 83% of the objects occurred 5 times or less and 96% occurred 10 or less times in the data set. *Propionibacterium*, *Acinetobacter* and *Comamonas* species objects made up most of the objects with 10 or more counts. See Table 4. The average number of common objects between sample pairs, the overlap, equaled 0.43, an indication of the sparseness of the data.

**Table 4:** Results of object merge transformations.

| Microbial Object | Occurrences | Microbial Object | Occurrences |
| --- | --- | --- | --- |
| Propionibacterium acnes-14 | 43 | Pseudomonas-hi | 13 |
| LoCnt-hi | 26 | Propionibacterium acnes-12 | 13 |
| Acidovorax-lo | 26 | Moraxella-hi | 12 |
| Moraxella-lo | 25 | Stenotrophomonas -lo | 11 |
| Streptococcus-lo | 23 | Nitrosospira-lo | 10 |
| Propionibacterium acnes-13 | 23 | Delftia-hi | 9 |
| Cloacibacterium-lo | 23 | Comamonas jiangduensis-lo | 9 |
| Pseudomonas-lo | 22 | Acinetobacter tjernbergiae-14 | 9 |
| Propionibacterium acnes-lo | 22 | Acinetobacter tjernbergiae-12 | 9 |
| Corynebacterium-lo | 22 | Acinetobacter baumannii-lo | 9 |
| Acinetobacter junii-13 | 22 | Streptococcus-hi | 8 |
| Acidovorax-hi | 22 | Methylobacterium-hi | 8 |
| Acinetobacter tjernbergiae-lo | 21 | Kocuria-lo | 8 |
| Cloacibacterium-hi | 20 | Kocuria-hi | 8 |
| Comamonas testosteroni-lo | 19 | Comamonas denitrificans-lo | 8 |
| Sediminibacterium-lo | 17 | Bacillus-lo | 8 |
| Acinetobacter tjernbergiae-13 | 17 | Acinetobacter junii-12 | 8 |
| Acinetobacter junii-14 | 17 | Achromobacter-lo | 8 |
| Sediminibacterium-hi | 16 | LoCnt-14 | 7 |
| Novosphingobium-lo | 16 | Achromobacter-hi | 7 |
| LoCnt-lo | 16 | Stenotrophomonas -hi | 6 |
| Delftia-lo | 15 | Noviherbaspirillum-hi | 6 |
| Bradyrhizobium-lo | 15 | Meiothermus-hi | 6 |
| Acinetobacter junii-lo | 15 | Bradyrhizobium-hi | 6 |
| Nitrosospira-hi | 14 | Bacillus-hi | 6 |
| Comamonas jiangduensis-hi | 14 | Methylobacterium-lo | 5 |

LDA can perform analyses where there is little overlap between pairs of samples but functions better the more overlap there is, and our binning and naming scheme left the data too sparse. The first runs suggested that the data could support about 5 classes. Since LDA finds within-sample object co-occurrences across many samples, we thought higher overlap would improve repeatability. Doing this objectively required filtering and grouping the data differently, specifically, reducing the microbial name specificity or the abundance resolution or both. We refer to this as object merging since different named and abundance-labeled objects are mapped to the same name and abundance label. In many microbiome studies, this is accomplished by summing higher up the phylogenetic tree, but this is a blunt instrument that would make it more difficult to see the biology if it were there. Our sequencing capability provided sub-species fidelity and we wished to retain as much information as possible. Summing higher in the phylogenetic tree turned out to be unnecessary to achieve repeatability.

Objects contain information about both the bacterium’s identity and its abundance in a sample which is a crude way of measuring their function, while not a detailed molecular biological characterization. When LDA results characterize a sample by class, the suggestion is that two samples in the same color class, even when they do not contain exactly the same bacterial objects, may have similar processes going on within them. In other words, the result is suggesting a commonality that could represent a redundancy of bacterial function.

Still, transformations that group different-named bacteria and that reduce abundance resolution could average out the biology. The goal is to transform the data to get reasonable repeatability but not go beyond what is needed. Of course, we did not know *a priori* whether we had enough data to achieve reproducibility without averaging out the biology nor where the line of optimal binning and naming was. The best approach was to achieve repeatability and hope the results revealed biology.

The next several paragraphs summarize how we arrived at a naming and binning structure that produced repeatable LDA results.

To begin with, we noticed that *Propionibacterium*, *Acinetobacter* and *Comamonas* species objects had class structure with one or two dominant classes that was apparent from the earliest runs. Individual species of *Propionibacterium* and *Acinetobacter*, particularly *P. acnes* and *A. junii* are examples. *Comamonas*, while its species objects have fewer total counts, had high counts per class in one class, increasing *their* statistical significance. So, we preserved the abundance resolution and naming from our starting point for *Propionibacterium* and *Acinetobacter* and preserved the name for *Comamonas* while reducing the abundance resolution. For other objects, the overall idea was that either by summing over lower abundances or lower occurring objects (name), we could get enough statistical significance to see a significant class structure emerge.

Objects with abundances over 12 had class structures where one component dominated the class distribution. Below 12, it was less clear mainly because objects had too few occurrences (counts). There were several high abundance but low count objects that occurred frequently in the magenta and red classes but whose samples jumped from one of these classes to the other with repeated runs, perhaps contributing to the repeatability problem. Last there were many objects with 5 or less occurrences.

In this scheme to reduce abundance resolution and naming specificity, we never cherry-picked by name or used disease state. We only used count cutoffs and abundance cutoffs applied to the entire data set. Because 12 was the abundance limit above which we could see structure with *Propionibacterium* and *Acinetobacter* we used this cutoff as we gradually realized that two abundance bins was what worked to get repeatability for other objects. We summed over the objects of questionable significance because of counts and filtered out very low count objects. This was not done in one step but through multiple runs of trying to optimize repeatability and graph quality.

It is useful to break the objects into four categories to describe the transformation from the original binning and naming: High-Count-High-Abundance (HiCnt-hi), Low-Count-High-Abundance (LoCnt-hi), High-Count-Low-Abundance (HiCnt-lo), and Low-Count-Low-Abundance (LoCnt-lo). The high-abundance cutoff was 12 or more and the high-count cutoff was about 10. The detailed parameters described below are a little different, but the following summary is easier to understand. Only resultant objects with 5 or more counts were retained. Please note that even when an object is mapped to a new name and abundance, it is still classified and contributes exactly the same to the LDA statistics but from a better-defined object.

HiCnt-hi: Consisted of mainly P and A objects. Kept initial abundance resolution over 12 and retained names.

LoCnt-hi: Consisted of many differently named objects mainly found in samples from red and magenta classes. Renamed to LoCnt. retained abundance 14 label with abundances of 12 or 13 mapped to hi.

HiCnt-lo: There were very few of these objects. Name retained. Abundance mapped to lo.

LoCnt-lo: Many different bacteria. When these were P or A, the name was retained, otherwise the object name was mapped to LoCnt and abundance to lo. 16 of these were kept.

LDA tends to defy standard statistical intuition though. While there may be a tendency to filter out low-occurring (low-count) objects, retaining these objects within a sample can reduce sample class distribution statistical error. This is why a trial-and-error approach, slowly adjusting parameters to get repeatability in terms of sample membership in each color class and cluster was an appropriate way to avoid classification bias.

The results are shown in Table 4. The specific transformations are detailed in Object Merging: Details below.

Overall, we reduced the number of objects from 218 to 69 with the abundance transformation and then to 52 with the name transformation. With this scheme, 2% of the objects occurred 5 times or less and 40% occurred 10 or less and we were able to increase the overall overlap statistics from 0.43 objects per sample to 0.86 objects per sample. The number of pairs with no overlap went from about 5,000 to 3,500 out of totals of about 7,500 (~120*120/2). Among pairs with non-zero overlap our scheme improved the overlap from 1.30 to 1.63. The three renamed objects had maximum classes as follows: LoCnt-lo was blue and both LoCnt-Hi and LoCnt-14 were magenta. See Table 4. These changes improved the repeatability (detailed definition below) into the 80-90% range.

##### Object merging: Details

After a great deal of LDA experimentation, we arrived at this scheme as described above.

###### Identify the following subsets of objects

Subset 1: Include any *Propionibacterium* (P), *Acinetobacter* (A) or *Comamonas* (C) species and genera from other objects with abundances ≥ 12 and occurring three or more times (specific objects for use in abundance transformation).

Subset 2: Include any species or genera not in subset 1 that occur ≥ 5 times (all objects of specific microbes for use in abundance transformation).

Subset 3: Include objects with counts > 1 and not P or A and £ 4 occurrences - (specific objects for use in count-based name transformation).

###### Perform the following transformations: for each of the subsets

1a) For subset 1, P & A species objects remain the same for abundances ≥ 12. All other abundances are mapped to -lo.
1b) For the rest of subset 1, if their abundance is 14, no change. For abundance 12 and 13 objects, the abundances are mapped to -hi and the remaining are mapped to -lo.
1c) For the objects whose microbes are not in subset 1 ≥ 12 is mapped to -hi, £ 11 to -lo.
2) For subset 2, for objects with abundances of 14, the object is not changed; if abundance = 12 or 13, mapped to -hi otherwise it is deleted.
3) Using the results of 1 and 2, for objects in subset 3, names are mapped to LoCnt. resulting in members of subset three being mapped to ‘LoCnt-14’, ‘LoCnt-hi’ and ‘LoCnt-lo’.

##### Deriving the LDA Classifier

Following is a heuristic derivation of Latent Dirichlet Allocation (LDA) classifier that hopefully provides a more intuitive sense of the algorithm. Rigorous derivations can be found here [68,75,78–82].

Overall, it is a stochastic approach that involves randomly assigning classes to the objects with a C-sided unequally weighted die where C is the number of classes used. In other words, it is a random process using the multinomial distribution. We will describe how to compute the probability weights of the die which are a function of the sample and object distributions introduced above. This set of die weights is also formally referred to as a classifier.

First, we derive the formula we used to compute the classifier weights and then we describe the iterative algorithm that uses the classifier to arrive at convergent values for the two distributions. In subsequent sections we describe additional practical details of the algorithm and improvements to it for this particular data set that, combined, we call MLDA.

Without resorting to fundamental probability theory, we offer the following heuristic derivation of the classifier. We assume that the classifier formula will depend in some way on the two distributions. Our approach is to find the simplest reasonable combination of the distributions. In this way, we arrive at a result that has an easy intuitive appeal.

The simplest expression that one can imagine for the weights involves the sum of four terms, two linear terms, a quadratic term and a constant. Each of the terms is a vector with length C. Keep in mind that the die roll assigns the class for one microbial object. All of the vector multiplications below are Hadamard or element-wise products where vector components from each vector are multiplied. The product of two 5-component vectors yields a 5-component vector.

They are as follows:

Term 1: It seems reasonable to suppose that the class of a particular object in a sample should be proportional to the class count distribution for the entire SAMPLE. The proportionality constant is a constant vector (vector of equal constants).

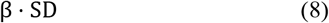

where SD means sample distribution.

Term 2: Since the object may occur in many samples, there should also be a term that reflects its classification in all of the samples which is the class distribution of the particular object being classified. The proportionality constant is a constant vector here as well.

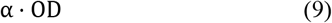

where OD means object distribution.

<u>Term 3</u>: There should also be a term that is the component-by-component product of these two distributions. The meaning of this is seen to be the joint distribution of the two.

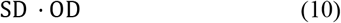

where for now we assume no proportionality constant.

<u>Term 4</u>: A constant, K.

Combining the four terms we obtain:

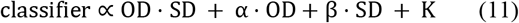

We can express equation (10) as the product of two terms.

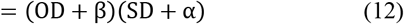

where the constant is seen to be:

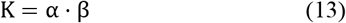

This way of expressing the relationship suggests the form of the denominator, D. For a frequency, we need an expression that is unit-less. Since the numerator is in units of counts, the denominator needs to be in counts as well. A reasonable linear form is have the object count term be divided by the total number of all objects assigned to each class, ∑_*all*_ O, plus a constant and the sample term be divided by the number of objects in the sample, NS, plus a constant. (Neither should include the object being classified which is required by probability theory and is also the case for the above distributions).

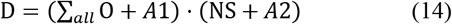

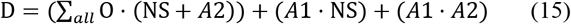

In practice, the first two of these terms dominate with the count product being the largest. We have found that the calculation converges with the Alzheimer’s data sets using the sum over objects term alone and normalizing. So in summary, the classifier is expressed in the following way:

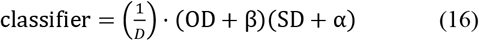

We can compare this result to equation 5 of [68] which is a rigorous result and see that it is the same result.

where P is the classifier and the *n’s* correspond to the distributions in the above formulas with

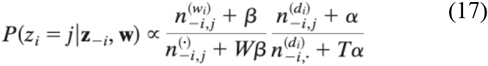

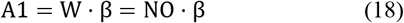

and

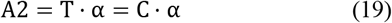

W is the number of unique objects in a data set which we call NO, and T is the number of classes in the computation which we call C in this paper.

The dot indicates that the distributions are to be calculated without using the object that is being classified. This is the case you obtain when the distributions are estimates of conditional probabilities which is technically what they are. The final expression is then:

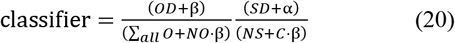

The components of the classifier, a vector with dimension equal to the number classes one is trying to classify into, are the weights of the die that we described above. This is the stochastic formula we use to classify an individual object. So, we will be classifying an individual object using the class distribution of the sample it is in and the object distribution of all the other occurrences of the object in the data set. We used the estimation rules for α and β presented in Griffiths and Steyvers [68] but also explored their parameter spaces. They suggest that *α* be 50/C. We adjusted to a lower value for α of 0.5 and retained the value of β suggested, 0.1, to obtain good graphical clusters.

##### Basic Latent Dirichlet Allocation (LDA) Algorithm

In general, since we do not know the distributions *a priori*, we begin the assignment computation by randomly assigning classes to all of the microbial objects in all of the samples using a die with equal weights. Then we cycle through the entire data set over and over (iterations) recomputing the assignments. Since the die weights are functions of the distributions which are obtained from the assignments, every roll of the die changes the die weights a small amount. Eventually, as the iterations proceed, the distributions stop changing and converge. It is a bit more complicated than this and we describe other features that are necessary for convergence below. Generally, we used 1000 iterations.

##### Modified LDA (MLDA)

Popular implementations of LDA algorithms use the sample and object distributions obtained after completing a number of iterations sufficient for these distributions to converge. For our data set, and in many others too, this approach does not result in repeatable distribution computations. We added the following procedures to improve repeatability.

###### Accumulation, Thinning and Burn-in

Rather than using the terminal distributions as the result, we accumulated the count distributions over the course of the iterations. To guard against correlations between iterations, we included results from one iteration after n iterations in the accumulation, typically 5. Sometimes this is called thinning. These two procedures were suggested by Heinrich [81,82].

When the classification begins, the distributions are far from convergence and would distort the accumulation if included, so we begin the accumulation after some number iterations, typically 100, determined after convergence monitoring (see below).

###### Randomization

The order of sample classification was randomized with every iteration. We did not randomize the object classification order within sample although it may be useful.

###### Classifier Mixing

After the burn-in we add additional averaging to the classifier beginning at 150 iterations which is performed for every object classification. This is done by computing the object and sample count distributions and then averaging in a small amount of the accumulated distribution, typically 4%, scaled by the ratio of counts in the current distribution to the accumulated distribution. The averaging is delayed until 150 iterations to allow for some accumulation to occur after burn-in.

###### Classifier Filtering

We filtered the classifier to include only the top three components, turning the classifier into a three-sided die.

###### Run Summing

The last layer of averaging added together multiple runs, usually five. We found that we could not just extend the length of a run to improve the convergence. After long runs, the classifications can destabilize and increase their entropy. We discovered that we could obtain better repeatability by summing multiple runs.

###### Mapping Classes before Run Summing

Since the computation begins with a random assignment of classes to the objects, the class labels (1, 2, 3, …) or their equivalent colors, change from run to run, requiring that they be mapped to a common labeling scheme before summing. We compared the samples by color class with assistance from the graphs to accomplish this.

If the computation and graph of individual runs are fairly repeatable, we should end up with the same clusters within statistical fluctuation but having different colors (labels). If our computations are not repeatable, we will see splitting of classes where, for example, the samples in cluster from one run end up with two or more other different clusters. On the other hand, if most of the samples from a particular class in one run correspond to most of the samples in another class in another run, we can see corresponding topologies and confirm it by checking which samples are in each cluster. In practice, run summing stabilizes the minority of samples that jump between color classes.

##### Convergence Monitoring

We monitor convergence in both a coarse and fine way. The coarse method is to compute the average entropy of the sample distributions and the object distributions. We normalize the entropy of the standard entropy by dividing by the entropy of a constant distribution for a vector of length C. This has the effect of making the entropy vary from zero to one.

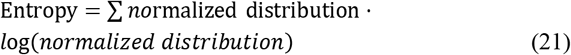

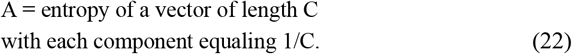

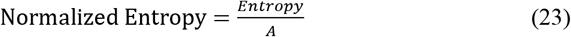

The entropy of the sample and object distributions tends to decline rapidly to values around 0.5 or below. Stable classifications will have lower entropy but low entropy does not always mean a stable classification so we also do the following.

For fine monitoring, we grab the maximum component label for each sample and each object. We make sure that the maximum components remain the same for 30-50% of the last iterations. See **Figure 5**. Iterations are plotted beginning after burn-in. If we look up the final component values of this object, we find that blue is 0.8, orange is 0.06, and green is 0.12. The graph shows that the main component, blue, is stable from after burn-in until the end. If there were hopping at the left end, we would have increased the burn-in. The two others are smaller components and so we expect some instability. In this way we checked the stability of all samples and objects. Generally, we strove to have the main component be very stable, giving us confidence that there were no issues with burn-in. In setting burn-in and other iteration parameters, we also made sure to be well beyond where entropy stopped declining.

##### Repeatability

For the graph to be a useful representation of reality, it must be reasonably repeatable. These are stochastic computations, so we do expect some variability between runs. Operationally, we defined repeatability in the following way. We performed multiple runs then added each sample’s count distributions together. We assessed the repeatability by comparing the populations of each color cluster of the summed graph with the populations of the color clusters of the individual runs that comprised it. The populations of the summed graph matched those from individual runs in the 80-90% range for each run in each color cluster.

##### Background removal and parameter values

We removed the OTUs present in the negative controls that we measured when implementing the second method. This removed a Staphylococcus epidermidis OTU that was not removed for the first method’s analysis. We further explore that when we compare the results of each method.

We did have a concern about contamination by *P. acnes* and *Acinetobacter* given their prevalence on the human body and in the environment respectively. We do not believe there to be a problem for three reasons. First, we presented findings in Data Filtering above that they were not. Second, they did not appear in the negative controls. Third, contaminants should be independent of class and not exhibit a class structure with peaked component maxima so the appearance of large class maxima in the objects of these bacteria argues against them being contaminants. Another way of saying this is that a contaminant object’s class distribution should have a high entropy close to 1.0. Most of the objects of *P. acnes* and *Acinetobacter* had entropies below 0.65. While it could be possible that some sort of systematic process outside of the subjects is responsible for their class structure, perhaps during the postmortem interval before the samples were extracted, the evidence that will be presented in the results argues strongly against such an explanation.

## RESULTS

### Individual Bacteria

After data filtering (low-yield samples and contaminant removal), 548 OTU and 108 samples remained. Infrequent OTUs (present in less than 20% of the samples) and low abundance OTUs (relative abundance ≤ 0.005%) were grouped into a composite feature named OTU others. After this step, the dataset contained 108 samples and 247 OTU (including OTU others). OTU were assigned to 229 species, although most of the species correspond to a single OTU, 14 species were assigned up to 3 OTUs.

At the phylum level, the major components (i.e., those with higher average relative abundance) were Proteobacteria (control = 47.35%, AD = 46.35%), Actinobacteria (control = 35.65%, AD = 30.62%), Firmicutes (control = 10.80%, AD = 15.17%), and Bacteroidetes (control = 5.44%, AD = 6.11%). Three OTU showed a broad prevalence across samples and were present in more than 50% of samples. They were assigned to the species *Propionibacterium acnes* (control = 82.69%, AD = 91.07%), *Acinetobacter junii* (control = 67.31%, AD = 55.36%) and *Staphylococcus epidermidis* (control = 55.77%, AD = 60.71%). Twenty-three OTU were present in more than 10% of the samples and 93 OTU were observed in only one sample.

The PCA on the clr-transformed OTU counts did not reveal any notable clusters of samples related to the disease status or biopsy sites (Figure 6), except for 14 control samples from 6 subjects that clustered together at the bottom of the PCA space. Only 32% of the variance was explained by the two first components.

**Figure 6:**
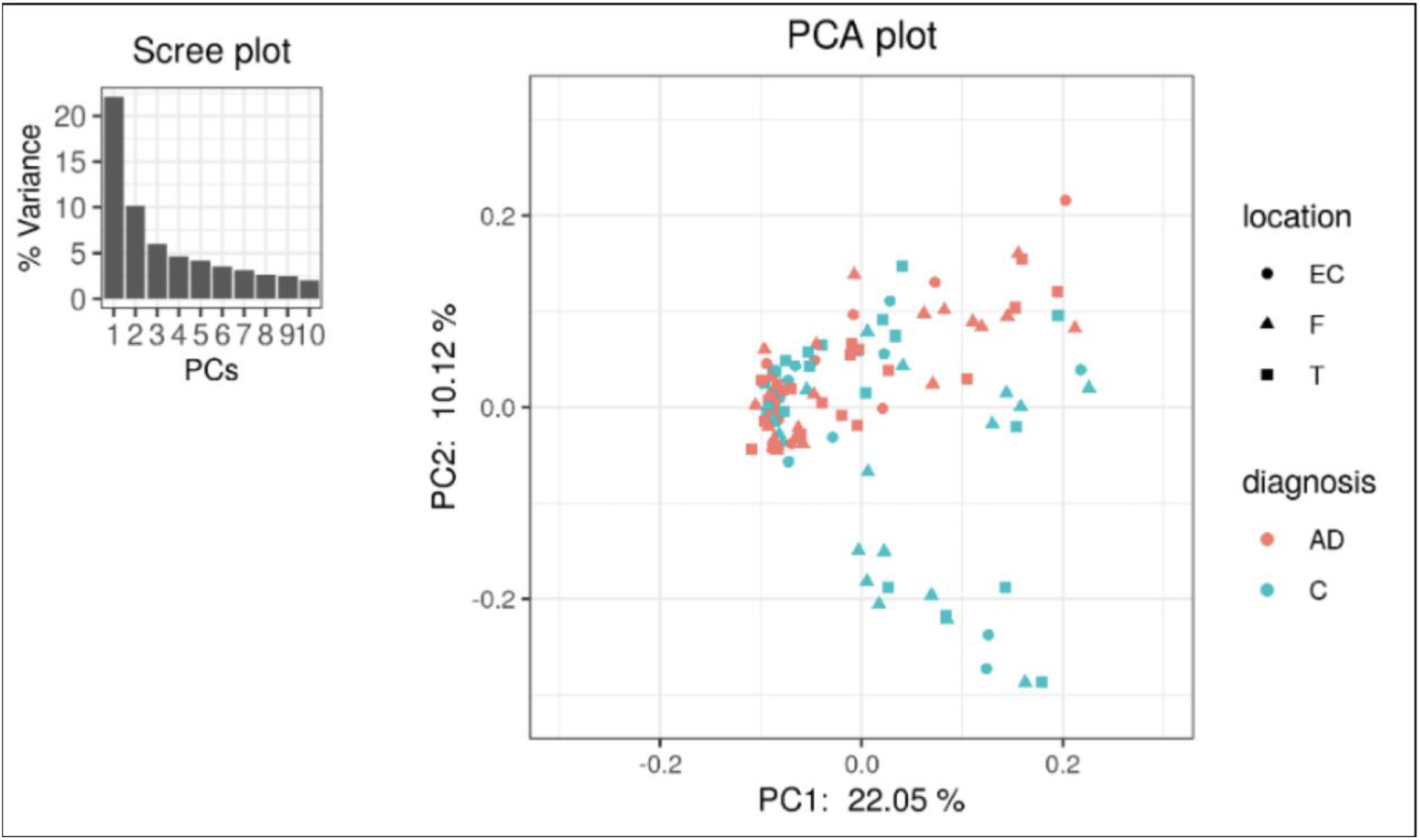
PCA was performed on clr-transformed composition. Each colored point represents a sample. Points are colored by diagnosis and shaped by biopsies location (EC: entorhinal cortex, F: frontal lobe and T: temporal lobe).

The heatmap of the top 80 most variable OTU, where the OTU and the samples were grouped by hierarchical clustering, shows that most of the samples were dominated by the same OTUs but did not evidence any pattern related to AD or control groups (Figure 7).

**Figure 7:**
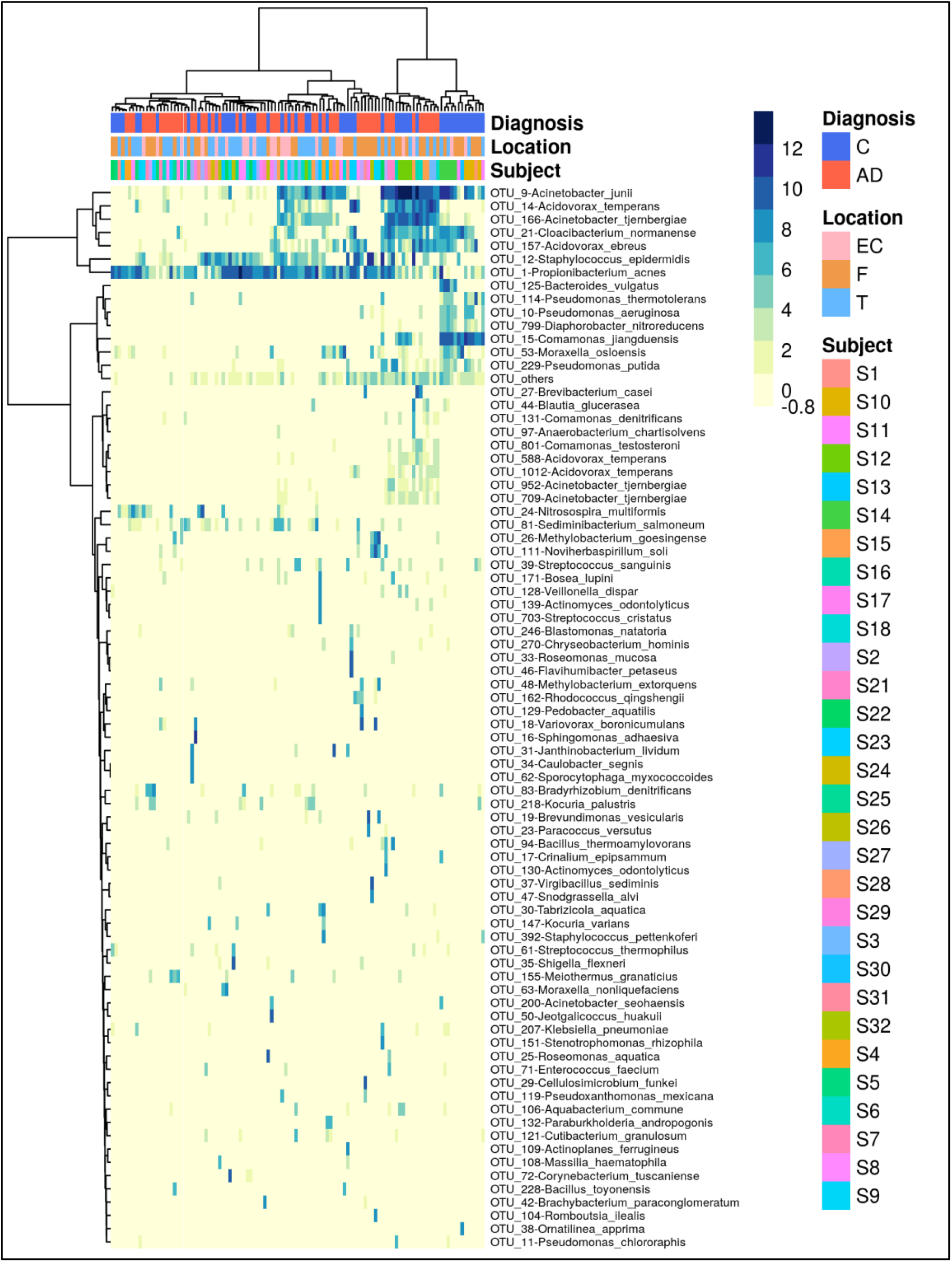
Heatmap that represents the clr-transformed OTU counts (more abundant OTU are darker in color) within each sample of the 80 most variable OTUs. The dendrogram was generated using the Euclidean distance between clr-transformed compositions. Sample’s subject, biopsy brain locations and diagnosis are indicated by the vertical colored strip. AD: Alzheimer’s disease; C: controls; EC: entorhinal cortex; F: frontal lobe; T: temporal lobe.

#### Difference in relative abundance between AD and Controls

Using DMM and assuming sample non-independence due to multiple samples coming from a single subject in the model, we found 12 OTU that shift in relative abundance between AD and control groups (Figure 8). Six OTU are more abundant in the control group: Acinetobacter junii, Comamonas jiangduensis, Cloacibacterium normanense, Pseudomonas putida, Pseudomonas thermotolerans, and Diaphorobacter nitroreducens. *C. jiangduensis, C. normanense, D. nitroreducens*and *P. putida* have low species-level confidence values (Table S2). The most important shift is in *A. junii.* Seven OTU are more abundant in AD group (*Propionibacterium acnes, Staphylococcus epidermidis, Acidovorax ebreus, Acinetobacter tjernbergiae, Acidovorax temperans, Noviherbaspirillum soli,* and *Methylobacterium goesingense*). *A. ebreus, A. tjernbergiae* and *N. soli*show very low species-confidence values (0.2112, 0.1169 and 0.0076 respectively). The most important change was in *P. acnes*. When the non-independence of the samples is ignored, the same results are obtained for *A. junii, C. jiangduensis, C. normanense, A. temperans, A. tjernbergiae, A. ebreus, S. epidermidis* and *P. acnes*, while no shift in relative abundance have been detected for *P. putida, P. thermotolerans, N. soli and M. goesingense* (Figure S7).

**Figure 8:**
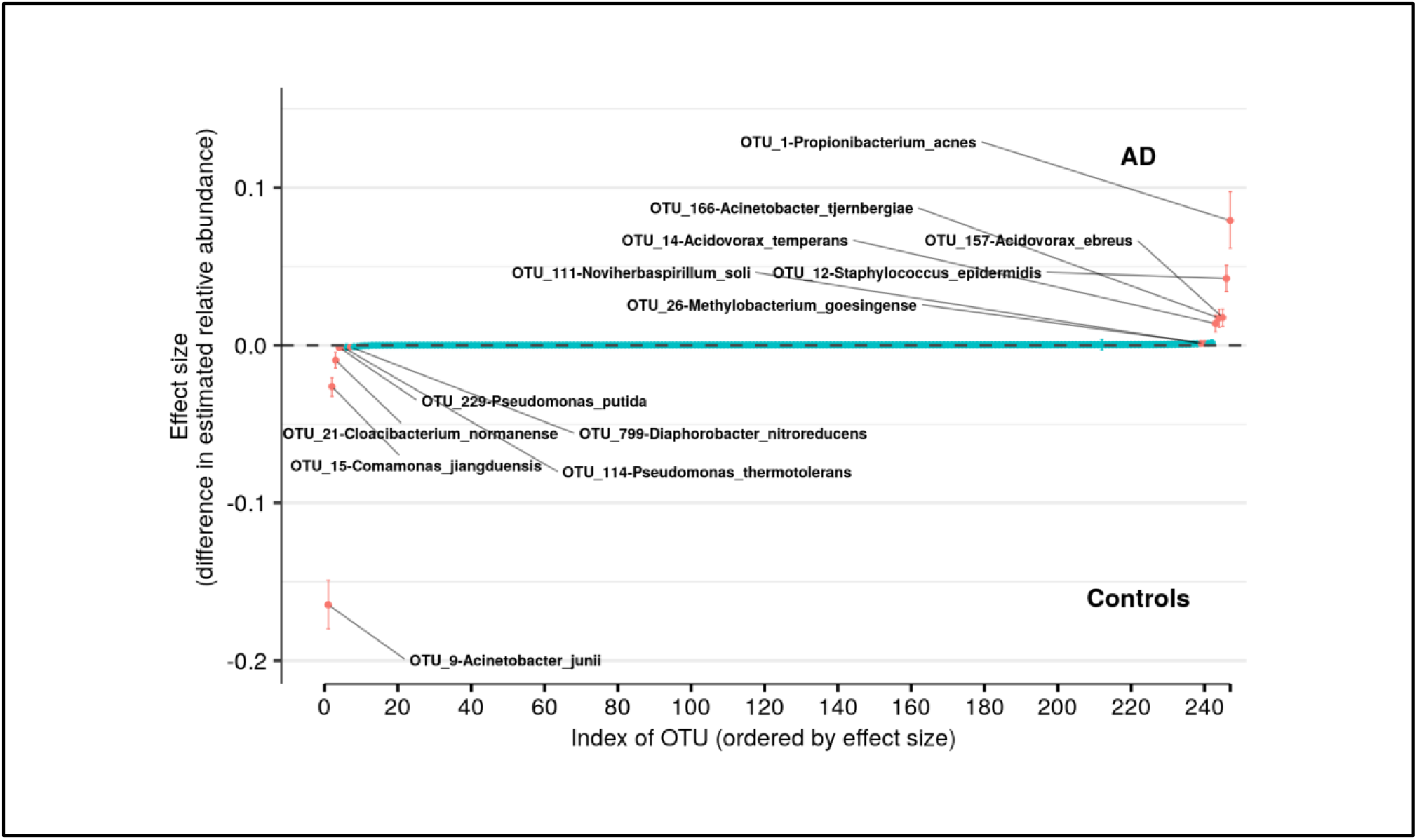
Differences in relative abundance between the Alzheimer’s disease (AD) group and the age-matched control group. The relative abundances were estimated for each OTU from each group through hierarchical Bayesian modeling. The vertical axis shows the difference for the estimated relative abundance of OTU between the AD and control groups. Points are the means of PPD and the whiskers show the 95% equal tail probability intervals of PPD (see Materials and Methods).

### Combinations of Bacteria

#### Introduction

Overall, we organized the results according to four themes: (1) the color classes, their microbiomes and their principal bacteria as revealed by the MLDA classification and graph methods, (2) microbe object abundance statistics that were used to infer the spatial distributions of underlying cellular scale ecosystems and the macroscopic distribution of ecosystem mixtures by class, (3) the relationships between the classes that will be used to determine the temporal order of the classes assuming each class represents different stages of underlying ecosystem evolution, and (4) the occurrence of the classes within each subject that suggests the pathogenicity of the ecosystems within each class. In this section, we focus on the mathematical results without detailed discussion of the ecosystem biology which will come in the discussion section.

##### Theme 1 - Color Classes and their Microbiomes

###### Microbiome Description

We used MLDA to compute five distinct color classes of samples. The results of our computations are summarized in a graph shown in **Figure 3** which is the sum of five different stochastic runs. We chose five classes as the optimum for our computation with the methods described previously.

The set of microbial objects resulting from a statistical summary of microbial objects from the samples in each color cluster will be called the class microbiome in the following paragraphs which approximates the rigorous microbiome. A more rigorous definition of microbiome would be the set of objects above a cutoff in the class column of the object distribution table. There are some differences, but the former is easier to understand because it ties to the data of the samples in the color class and does not contain objects from samples of other classes. See, for example, Figure 4 which shows a graph rigorously computed from the MLDA object results. The observed microbial objects derive from the summation of one or more ecosystems at the cellular scale during the physical sampling process. We will show how to characterize these ecosystems and how they determine the sample measurements in the discussion section.

Further characterizing these five homogeneously colored clusters, two are very distinct, the green and orange, while the other three are merged together: blue, red and magenta. This means the green and orange microbiomes are both different from one another and different from the blue, red, and magenta supercluster microbiomes. On the other hand, the latter are more related to one another since they are closer to one another. Each cluster represents a different underlying microbiome with a different set of microbial objects statistics. The green and orange sets must therefore have very different objects while the blue, red and magenta have objects in common, specifically blue with red and red with magenta although not blue with magenta. As we have pointed out before, common objects between classes could be suggestive of multiple meanings of an object. This is one of the peculiarities and benefits of MLDA, that identical measurements mean different things in different contexts.

The statistical results shown in Table S3, Table S4 and Table S5 show these microbiome summaries for each of the color classes for objects with abundance bin ≥10 (0.3% abundance) and counts ≥5 in Table S3 and ≥2 in Table S4 and Table S5. Different tables show different abundance combinations for particular microbes to show the importance of various microbes. Table S8 shows the same information in a different form. Each object is shown with their approximate microbiome computed from the occurrence count of the object in the sample color class.

It is important to emphasize that the graph is just as important a part of the results as the numbers behind it. While any feature is directly attributable to the underlying class distribution of the nodes, it is exceedingly difficult to see many features of the results and subtleties by inspecting a table of numbers or plotting statistical summaries of specific objects. By viewing the graph, it is possible to see tightness of the clusters, distinctness of the clusters, or nodes that may be dominated by a statistical fluctuation in the classification, etc. These features provide insights into the computational results. We comment on many but not all these features throughout the results section.

For each color class, we also show the fraction of samples that come from AD subjects. Diamond-shaped nodes are from AD subjects and circles from the controls. It does not necessarily follow that this number is an estimate of the pathogenicity of the underlying microbiome which will be emphasized in a number of contexts.

#### A Few Graph Anomalies

In **Figure 9**, we show the sample maximum component distributions for the samples in each class. This gives you a sense for how the maximum components are distributed for each class. It is apparent that most are over 0.4. That is, the components are nowhere near equally distributed. So, when the maximum component falls below this value or when the second component is close to the maximum, sample nodes start being pulled out of their color cluster. From right to left in **Figure 3** we provide the relevant components of the nodes that have been pulled from their clusters. The first color is the sample color followed by one or two that are between ~0.2 (evenly distributed) and ~0.4: red:green, blue:orange, red:blue-orange, magenta:green-orange, magenta:orange, magenta:blue-orange-green. Referring to **Figure 3**, with the exception of the last anomalous node, it is possible to see how the nodes are pulled between the first color and those colors after the colon.

**Figure 9:**
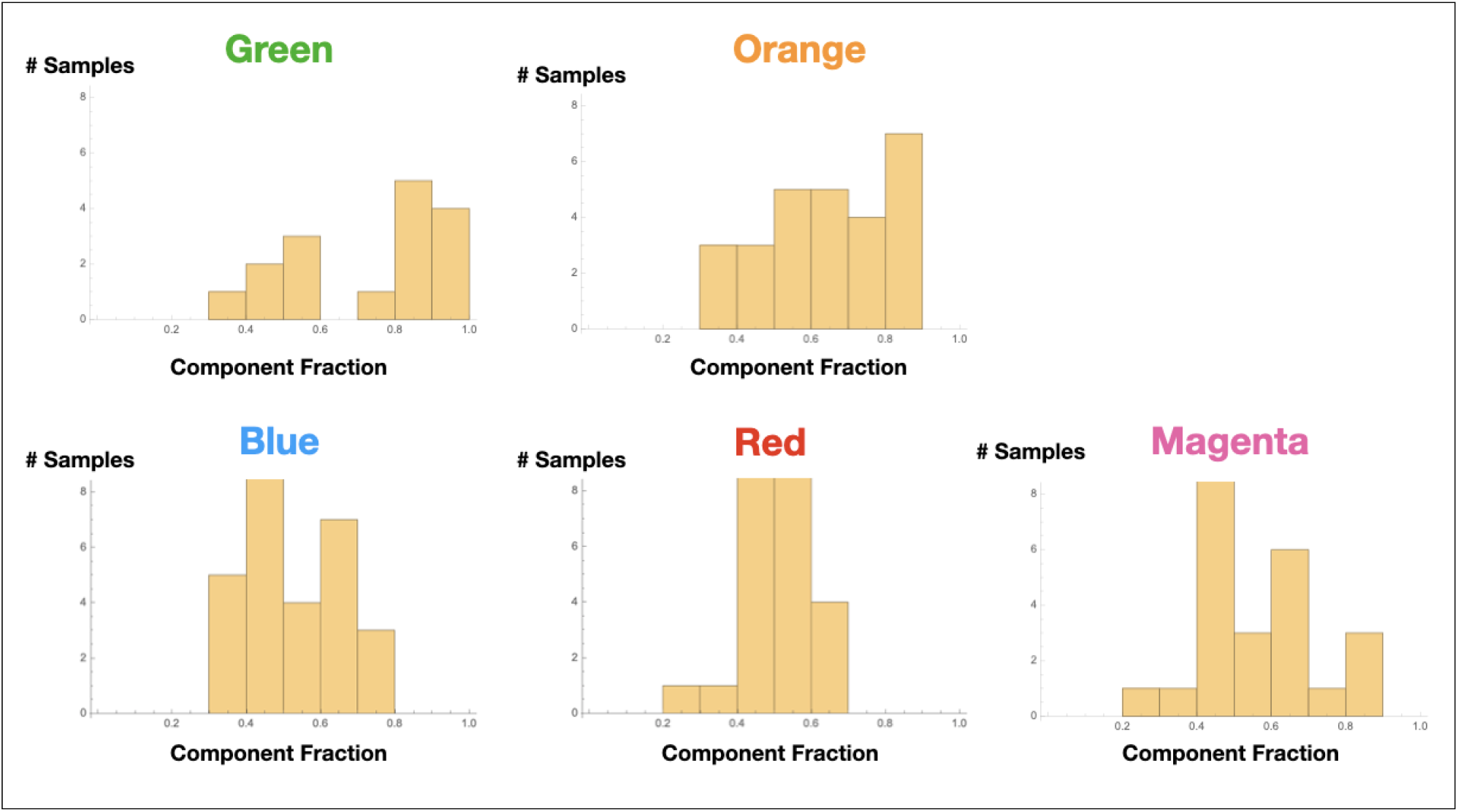
Sample maximum component distributions by class.

#### Principal Microbes and Microbial Objects

In the Figure 10 graphs, we show the samples in which the most abundant species are found, combining their objects in abundance ranges 12-14. The samples that contain these objects are shown by enlarging the sample/nodes containing these microbial objects. *P. acnes* is most prevalent in the blue-red-magenta supercluster with some in the orange. The blue is dominated by the 14 abundance range, the red has a combination of 14 and 13 abundance ranges and the magenta a combination of 13, 12 and 11 abundance ranges. *Acinetobacter junii* is mainly found in the green and orange classes and *Comamonas jiangduensis* is found almost exclusively in the green class. These underlying details can be found in Table S3, Table S4, Table S5, Table S8 and Table 5.

**Figure 10:**
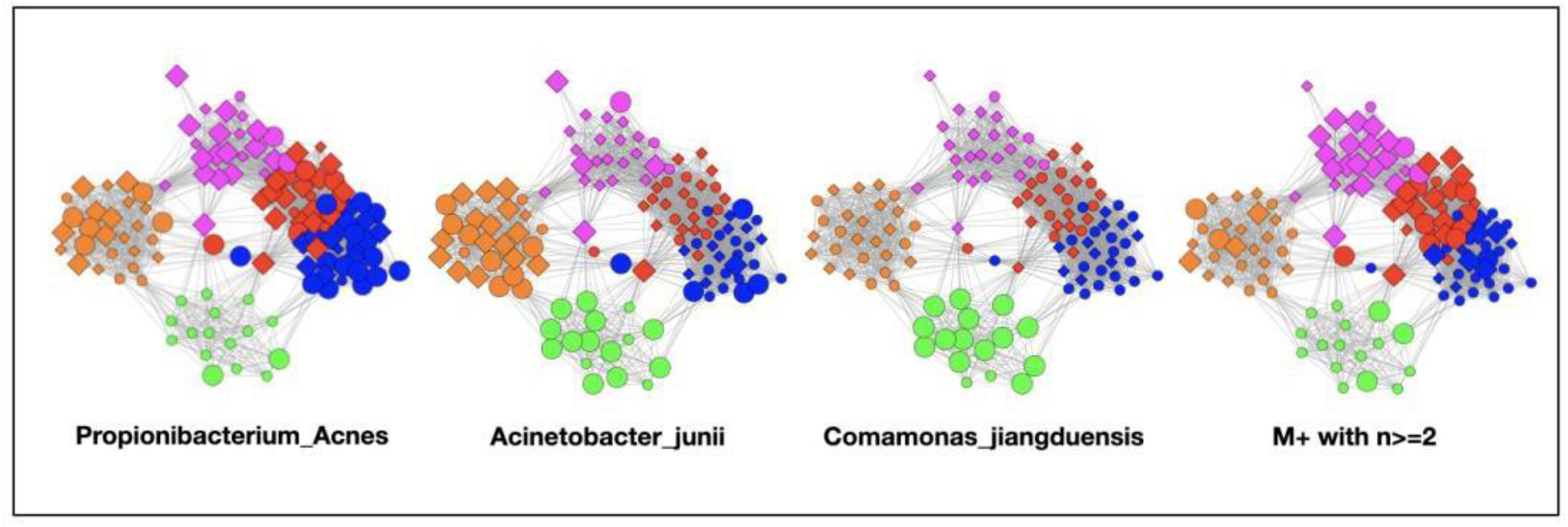
Graphs showing samples with abundances levels 12-14 with enlarged nodes for three main microbes.

**Table 5:** Principal bacteria abundance distributions. Note that the M+ rows are different because they show the occurrence of any of 21 different genera in the M+ set.

|  | Microbe/<br>Abundance Bin | 7 | 8 | 9 | 10 | 11 | 12 | 13 | 14 | tot/<br>tot-class | Abun-<br>dance | Width | Avg | SD | Sum |
| --- | --- | --- | --- | --- | --- | --- | --- | --- | --- | --- | --- | --- | --- | --- | --- |
| Green | *P. acnes* | 1 | 1 | 0 | 1 | 3 | 0 | 1 | 1 | 8/16 | low | wide | 10.6 | 2.3 | 8.0 |
|  | *A. junii* | 0 | 0 | 0 | 1 | 0 | 2 | 5 | 5 | 13/16 | med | narrow | 13.0 | 1.2 | 13.0 |
|  | *C. jianduensis* | 0 | 0 | 0 | 0 | 1 | 2 | 5 | 7 | 15/16 | med | narrow | 13.2 | 0.94 | 15 |
|  | M+ n>=2 | 0 | 4 | 1 | 4 | 3 | 1 | 1 | 1 | 8/16 | low | wide | 10.2 | 1.9 | 15 |
|  | M+ | 2 | 7 | 6 | 6 | 6 | 2 | 1 | 1 | 15/16 | low | wide | 9.7 | 1.7 | 31 |
| Orange | *P. acnes* | 2 | 1 | 3 | 3 | 2 | 3 | 7 | 2 | 23/27 | med | med | 11.1 | 2.2 | 23 |
|  | *A. junii* | 0 | 0 | 0 | 0 | 1 | 0 | 12 | 11 | 24/27 | high | narrow | 13.4 | 0.71 | 24 |
|  | *C. jianduensis* | 2 | 0 | 1 | 2 | 3 | 0 | 0 | 0 | 8/27 | low | wide | 9.5 | 1.7 | 8.0 |
|  | M+ n>=2 | 1 | 4 | 4 | 5 | 5 | 5 | 1 | 0 | 15/27 | low | wide | 10.1 | 1.6 | 25.0 |
|  | M+ | 1 | 7 | 8 | 8 | 7 | 6 | 2 | 1 | 20/27 | low | wide | 10.1 | 1.7 | 40.0 |
| Blue | *P. acnes* | 0 | 0 | 0 | 0 | 0 | 0 | 0 | 27 | 27/29 | high | narrow | 14.0 | 0 | 27 |
|  | *A. junii* | 0 | 0 | 3 | 5 | 2 | 3 | 3 | 0 | 16/29 | low | wide | 10.9 | 1.5 | 16 |
|  | *C. jianduensis* | 0 | 0 | 0 | 0 | 0 | 0 | 0 | 0 | 0/29 | none | none | 0.0 | N/A | 0 |
|  | M+ n>=2 | 0 | 5 | 3 | 11 | 5 | 5 | 1 | 1 | 18/29 | low | med | 10.3 | 1.5 | 31 |
|  | M+ | 0 | 5 | 6 | 16 | 10 | 9 | 4 | 3 | 24/29 | low | med | 10.7 | 1.6 | 53 |
| Red | *P. acnes* | 0 | 0 | 0 | 0 | 1 | 2 | 6 | 14 | 23/26 | high | narrow | 13.4 | 0.84 | 23 |
|  | *A. junii* | 0 | 0 | 0 | 1 | 2 | 0 | 1 | 0 | 4/26 | low | wide | 11.3 | 1.3 | 4 |
|  | *C. jianduensis* | 0 | 0 | 0 | 0 | 0 | 0 | 0 | 0 | 0/26 | none | none | 0.0 | N/A | 0 |
|  | M+ n>=2 | 0 | 1 | 1 | 4 | 9 | 10 | 9 | 7 | 25/26 | med | med | 12.0 | 1.5 | 41 |
|  | M+ | 0 | 1 | 1 | 4 | 9 | 12 | 11 | 10 | 25/26 | med | med | 12.1 | 1.5 | 48 |
| Magenta | *P. acnes* | 0 | 0 | 0 | 1 | 3 | 7 | 9 | 0 | 20/24 | med | narrow | 12.2 | 0.89 | 20 |
|  | *A. junii* | 0 | 0 | 0 | 0 | 0 | 3 | 1 | 1 | 5/24 | med | narrow | 12.6 | 0.89 | 5 |
|  | *C. jianduensis* | 0 | 0 | 0 | 0 | 0 | 0 | 0 | 0 | 0/24 | none | none | 0.0 | N/A | 0 |
|  | M+ n>=2 | 0 | 0 | 2 | 5 | 2 | 5 | 6 | 12 | 21/24 | med | med | 12.4 | 1.7 | 32 |
|  | M+ | 1 | 1 | 2 | 7 | 6 | 9 | 8 | 19 | 24/24 | med | med | 12.2 | 1.8 | 53 |

#### Low Count Microbial Objects

To understand the low-count microbial objects, it is best to look at high abundance and low-abundance objects separately. Because we are spreading low count objects over five classes, it is difficult to find statistically significant patterns among them. In fact, we were unable to find any statistically sound patterns among the low-count and low-abundance objects. We did, however, find a fundamentally important pattern for high-abundance low-count objects that occurs mainly in the magenta and red classes although signs of it can be traced to the other classes too. See Table 5. Specifically, we noticed that samples with *P. acnes* with abundances 11-13 in the red and magenta classes correlated with a set of low-count bacteria with abundance level 14. In most cases, there was only one that occurred per sample. In Figure 11, we show two different ways of defining this set. This is mainly the Low-Count-High-Abundance objects that were renamed to LoCnt-H or LoCnt-14i in the Abundance Binning, Microbe Naming and Object Merging section and whose maximum component was magenta.

**Figure 11:**
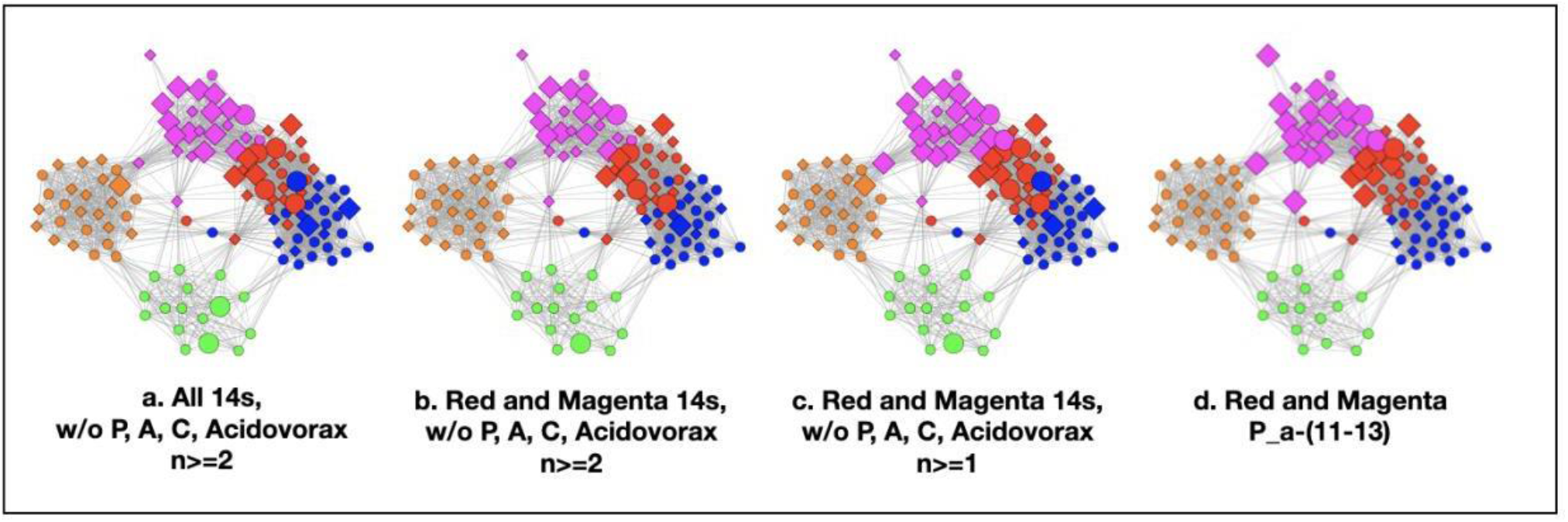
Several definitions of M+ compared to P. acnes (11-13). (a) objects with level 14 that occur 2 or more times in any class, (b) objects of level 14 that occur 2 or more times in magenta or red (c) objects of level 14 that occur 2 or more times and their corresponding objects of level 13 in magenta or red, (d) P. acnes (11-13).

In (a) we show all objects with abundance level 14 that occur at least twice except *P. acnes*, *Acinetobacter spp., Comamonas spp.* and *Acidovorax spp*. *Acidovorax* is not shown here because it is mainly found in the orange class, and we are focused on red and magenta for the results of these paragraphs. This graph shows that the phenomena of the preceding paragraph occur mainly in the magenta and red classes. In (b), we show abundance 14 occurrences from magenta and red wherever they occur and when they occur at least twice. There are some that also occur in blue and green but not many. In (c) we show abundance 14 occurrences from magenta and red wherever they occur.

From here on, we will refer to the abundance 14 objects of (c), occurring in the red and magenta classes, as the M+ set because the most prevalent member of the set is *Methylobacterium* (including level 13 abundances). They include the following 21 genera: *Achromobacter, Bacillus, Blastocatella, Bosea, Bradyrhizobium, Brevundimonas, Caulobacter, Delftia, Ferrovibrio, Janthinobacterium, Kocuria, Massilia, Methylobacterium, Nitrosospira, Rubellimicrobium, Sediminibacterium, Sphingomonas, Stenotrophomonas, Streptococcus, Variovorax, and Virgibacillus*. The 9 objects that occur 2 or more times from (b) are: *Bacillus, Bradyrhizobium, Caulobacter, Delftia, Kocuria, Methylobacterium, Nitrosospira, Sediminibacterium, and Variovorax*. Their objects can also be found in Table S4.

In (d), we show the samples where *P. acnes-*(11-13) objects occur to demonstrate that there are many samples that have both M+ objects and *P. acnes-(11-13)* objects. In fact, of the 28 samples where these *P. acnes* objects occur, 22 contain M+ objects. A couple more have abundance 13 objects from the M+ set. In other words, there is a very large overlap between samples with high abundance M+ objects and *P. acnes-(11-13)* objects. Because of the low occurrence of the M+ objects, it would have been easy to argue they were not important. Their correlation with P-11-13 objects and their ability to improve the repeatability of the MLDA computations when mapped to LoCnt-hi caused us to take notice.

#### *P. acnes* Results at Higher Abundance Resolution

In Figure 12, we show a sequence of graphs where we have selected samples that fall in finer *P. acnes* abundance bins than the logarithmic bins that define the objects. We have set the size by finding the bins that have at least ~15 samples. It is apparent that the successive selections going from high abundance to low abundance show clusters that roughly move from blue to magenta indicating that we have not averaged out the *P. acnes* abundance information by using a binning structure that is too coarse.The fact that the graph can preserve this feature by combining the MLDA and graph embedding is rather remarkable especially since the bin merging reduced the abundance resolution.

**Figure 12:**
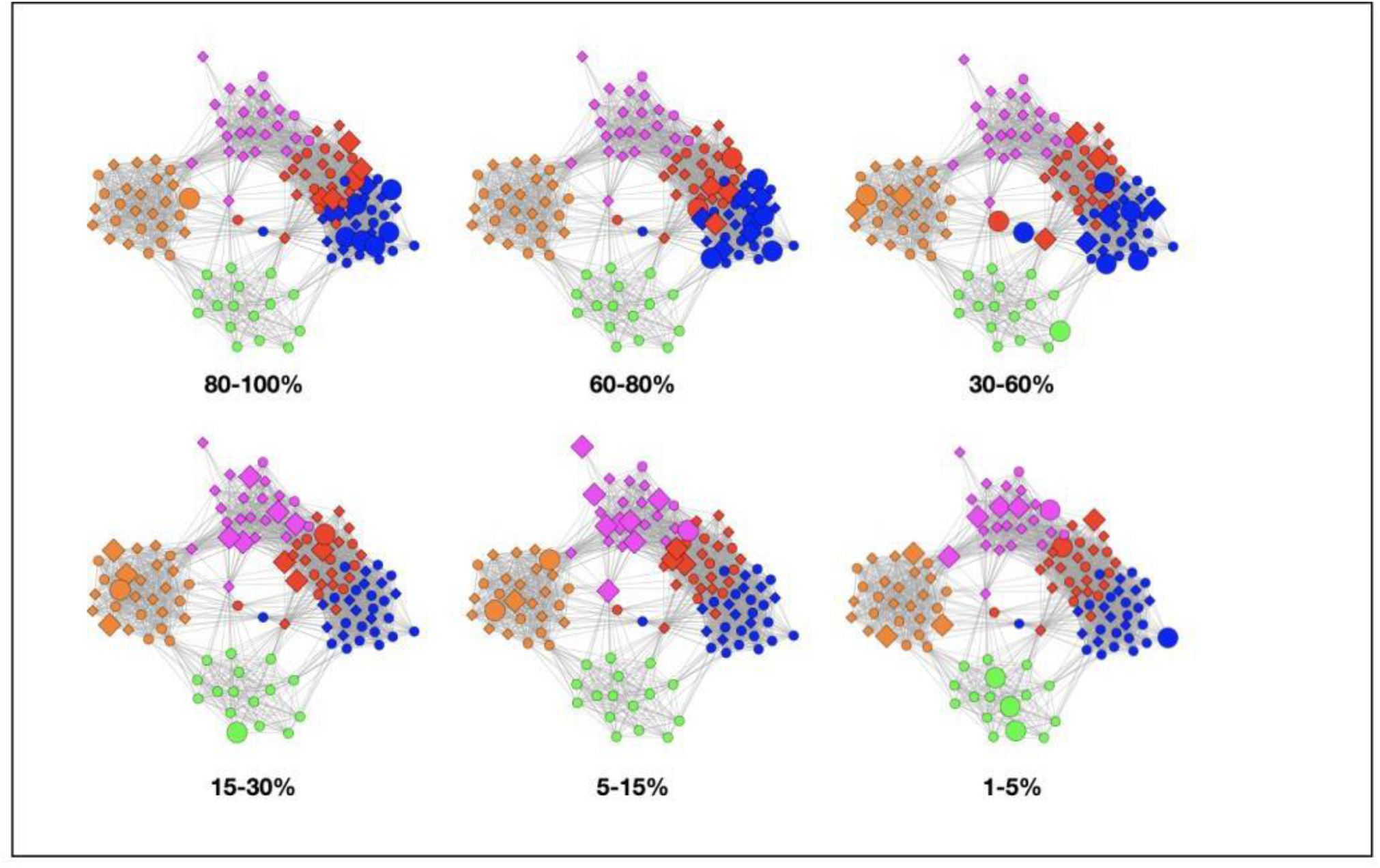
Graph sequence showing P. acnes abundance dynamics.

#### Robustness of Color Clusters

Three of the color classes have a heterogeneous mix of AD and control samples (orange, blue and red). The other two, green and magenta, are nearly homogeneous in disease state, comprising almost entirely samples from either AD or control subjects. We initially thought that we should observe clusters whose disease state was nearly homogeneous meaning that either a microbiome was pathogenic or not. The MLDA results required a more complex explanation but first we needed to establish that they were robust.

We thought that these statistics suggested that there may be within-class differences in the microbiomes of the AD and control samples that could split the orange, blue and red classes into homogeneous color clusters with the right input and object merging parameters. Indeed, there were within-class differences but when we tried to split the clusters by using a larger number of input classes with these adjustments, the clusters would not split. The object differences between samples were just not large enough to support entirely new color classes.

To gain a better understanding of this result, let us describe the node attributes that bring two nodes near one another in a graph. For two samples to be nearby they must have similar classifications. Containing the same objects helps but the nodes also must have common co-occurring objects. Our inability to split the orange, blue and red classes indicates that the attributes that cause these clusters are stronger than the differences between the AD and control subsets of these classes.

The examples of *P. acnes-14* and *P. acnes-13* are illustrative. *P. acnes*-14 occurs 27 times in blue and 14 times in red and is the highest occurring object in both color clusters. See Table 5. These samples end up in different clusters because they occur with other objects too that are not the same. *P. acnes*-13 occurs 9 times in magenta and 7 times in orange. While *P. acnes-*13 is the highest occurring object in magenta, it is only the third highest in orange. Again, these samples end up in different color clusters partly because *P. acnes*-13 occurs with other objects in the two clusters even though it is in common among the 16 samples.

One of the lessons of the MLDA/graphical analysis is that whenever you focus on a particular object, you are always reminded of its relationship to all other objects by its samples’ color and position in the graph. If you forget this by plucking out one object or one microbe and analyze it in isolation, assuming it has a single meaning, it is easy to obtain misleading results and fool yourself. The fundamental reason for this is that you are effectively ignoring confounding variables, namely the classes.

#### Correspondence With Results of Method I

We will look at three aspects of these results in light of the MLDA results. First, we will compare the PCA method’s capability of identifying microbiome clusters, second, we will look at the bacteria that were identified as being correlated or anti-correlated with AD and last, we will look at a contamination issue.

In Figure 13, we show the same chart as Figure 6 except that the nodes are colored by the class colors of the MLDA results. The green and orange samples are distinctly clustered while the blue, red, and magenta are mixed together. While the orange are clustered, there is not an obvious way to distinguish the colors, at least with two PCA components. The MLDA plus graph does show the blue, red, and magenta samples in a super cluster however these color clusters are distinct and defined by the maximum class component whereas they are interspersed in the PCA results. This may be due to the lack of the equivalent of the object factor in the classifier in PCA, but this is the subject beyond the scope of our work.

**Figure 13:**
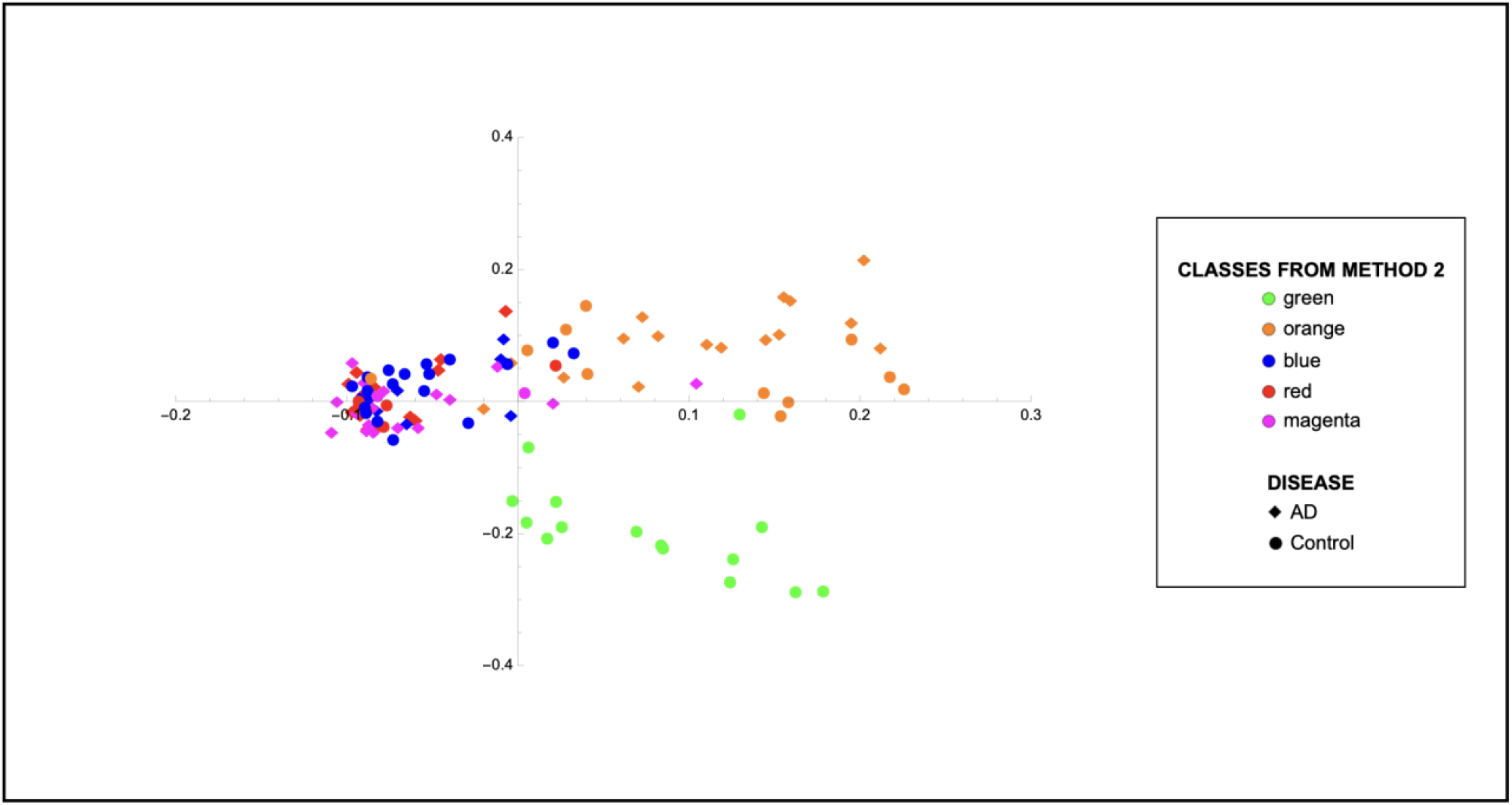
This is the same as **Figure 6** but the nodes are colored by the class colors.

For the following, See Table S1, Table S4 and Table S5. The DMM method reveals *Propionibacterium acnes* as associated with AD. DMM has no class structure, so the method essentially averages over it. As we discuss both in this section and the next, this association is likely due to the prevalence of *P. acnes* in abundance levels 11-13 in the red and magenta clusters being high enough not to be washed out by its presence elsewhere where it is frequently found in the controls.

The method also found *Comamonas jiangduensis* to be correlated with the controls which is also found with the MLDA method. Regarding *Acinetobacter,* one species A. junii was found to be anti-correlated with AD and another A. *tjernbergiae* was correlated with AD. A. junii objects are found in both the green and orange classes, with green not being associated with AD and orange somewhat associated with AD. A. *tjernbergiae* objects are mainly found in the orange class. Overall, since DMM is essentially averaging over the classes these results appear to be consistent with the MLDA results.

Last, but important, the *Sediminibacterium* and *Methylobacterium* species identified by DMM as associated with AD is consistent with the MLDA findings. These two are among the M+ bacterial set that is found in the red and magenta classes. *Methylobacterium* is mainly found in the samples of the magenta class, which is most associated with AD. Other findings of DMM can be reconciled with MLDA by referring to the Table S1, Table S4 and Table S5. Overall, correlation of the results with AD is more complicated than this discussion and requires an understanding of the subject level results.

The first method found that *Staphylococcus epidermidis* was associated with AD. The main OTUs in this species were, however, removed in the background removal process for the second method because all of the OTU’s present in the negative controls were removed even if there was only a small amount as was the case for this OTU. A post MLDA analysis found that S. epidermidis was present in 45 samples ≥ abundance level 11 and in 39 samples ≥ abundance level 12. Since we know what the color classes of the samples containing these objects are, we can estimate an object class distribution based on their occurrence and compute its entropy. The entropies using either an abundance cutoff of 11 or 12 were over 0.95, i.e. a fairly flat distribution, suggesting that they are contaminants and that the findings of the first method are spurious for *Staphylococcus epidermidis.* Its objects with abundances ≥ 12 come from AD samples 59% of the time partially accounting for the DMM result.

In summary, the results of the DMM analysis are what happens when class is not considered in an analysis. As it is a confounding variable, ignoring it can sometimes skew results, though not always.

##### Theme 2 -Microscopic structure and Macroscopic Spatial Distribution

There are two sets of results that provide information about the macroscopic spatial distribution of ecosystem mixtures and microscopic spatial distribution of individual ecosystems within the samples. We describe the results here and review their detailed relationship to ecosystems in the discussion section.

###### Graph Clustering and Macroscopic Structure

A fundamental result of the graph visualization of the MLDA results is the appearance of homogeneously colored clusters. The colors show the maximum MLDA component of the node classification vector. Since there is always a maximum component, we emphasize that the graph embedding algorithm does not just gather the samples with the same maximum component. Only when the MLDA computation has resulted in sample distributions where the maximum component is large, typically over ~0.40 do you get homogeneously colored clusters. In other words, > 40% of the microbial objects have the same classification for each of the samples of a particular color cluster. If the components of the samples were more evenly distributed, where class components were ~0.20, the graph would show a multicolored cluster. **Figure 9** shows that all five classes have samples whose maximum component is 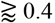.

Even though there are not enough samples per subject for significant spatial correlations computations, the class structure suggests another mathematical view of the results. The color clusters are telling us that even though the samples may come from different subjects, they have strong similarities. This may justify picturing the sampling process as coming from two virtual brains, one with AD and one without AD.

We will argue below that this spatial correlation suggests that large regions of each of these virtual brains have the same microbiome (ecosystem mixture), characterized by class. This conclusion, however, is dependent on a model of the large-scale spatial distribution which will be explored in the discussion.

###### Abundance Distributions of Principal Bacteria and Microscopic Structure

By examining the actual underlying objects in each color class, we can learn even more. In Table 5, we present the abundance statistics of each of the principal objects in each class, specifically *P. acnes*, *A. junii*, *C. jianduensis* and the M+. The principal *P. acnes*, *A. junii*, *C. jianduensis,* and M+ object abundance averages and distributions within a class provide information about the microscopic ecosystem structure, specifically their density and their spatial homogeneity on sample scales. In other words, these statistics provide information regarding the structure within the sample from which we can infer a microscopic structure.

We show the occurrence of each object in each class and then characterize the width of the abundance distribution over bins. The average abundance is related to the density of underlying ecosystems and the width of the abundance is related to the spatial homogeneity of ecosystems within a sample. More details of this interpretation are provided in the discussion. Generally, when the abundance average is ~ 14, we call it high; between 13 and 14, we call medium; and everything else is called low. Also, when the distribution is concentrated in one or two bins, we call it narrow, mainly two or three bins with some peaking, medium, and greater than or equal to 3 bins, we call wide. We will explain in the discussion how to predict the spatial distribution of underlying ecosystem mixtures from these results which is not obvious because it depends on the ecosystem model. The sample data is presented in Table S6.

In presenting the results in this manner, we also need to call attention to an important equivalence principle that we use to understand these results. Further, we emphasize that we are assuming that the microbial objects used in the computations result from summing over physically sampled mixtures of ecosystems, but we do not know much about the ecosystems yet. Therefore, we are assuming that each class microbiome results from a different mixture of ecosystems. The principle is as follows.

##### The sum of the virtual sampling of ecosystems mixtures equals the physical sampling of the sum of the ecosystem mixtures

In other words, we can treat the results of Table 5 as what we would obtain had we been able to individually sample a single ecosystem class mixture. We can then use these results to derive something about the nature of the individual class ecosystem mixtures. This is done in the discussion section.

Below, we describe the principal microbial objects of each class, their class occurrence, average abundances, and the width of the distribution. There are, of course, other objects which can be seen in Table S3, Table S4, Table S5 and Table S8. The dominant bacteria are also made clear in Figure 3.

###### Green

The most prevalent taxa are the *Comamonas* and *Acinetobacter* species shown in Table 5. Their average abundances are medium, and the width of their abundance distributions is narrow. The abundance of *P. acnes* is low, and its width is wide.

###### Orange

The most prevalent bacteria are species of the *Acinetobacter* genus. The abundance of the principal *Acinetobacter* species, *A. junii* is high with a narrow distribution width. There was another species of Acinetobacter, *A. tjernbergiae,* with a significant occurrence, but its green-orange dynamic was different from *A. junii* although it was similar. This is more fully discussed in **The *A. junii-14* to *A. junii* −13 Orange Class Transition** paragraph below. Its involvement, however, in a complex dynamic with *A. junii* is responsible for a significant part of the overlap among samples in the orange class. Orange samples have minimal amounts of *Comamonas* species. The abundance of *P. acnes* is medium with a medium width distribution, occurring in nearly all samples.

###### Blue

*P. acnes* occurs in almost all samples with a very high abundance and a narrow distribution. *A. junii* is present in low abundances with a wide distribution width. *Comamonas spp.* are not present.

###### Red

*P. acnes* occurs in all but three samples with somewhat lower abundance than in the blue class but still with a narrow width. The abundance of *A. junii* is present at low abundance with a wide distribution. There is a high occurrence of the M+ genera particularly with abundance levels 13 and 14 and medium distribution width.

###### Magenta

This class is dominated by the M+ set of genera and they occur in all samples with a medium abundance average and medium width. A significant fraction of the samples has abundance 14 M+ objects. *P. acnes* has a medium abundance and narrow distribution. There is a minimal amount of *A. junii*.

##### Theme 3 - Temporal Order of the Classes

Temporally ordering the classes requires finding relationships between pairs of classes and then determining their temporal order. Below, we describe statistical methods that relate class pairs where it can be argued that the method is finding classes where the underlying ecosystems could evolve from one class to the other. Using these pairs, together with arguments about temporal order, we construct a time-ordered network of the classes.

###### Microbial Abundance Dynamics

A strong statistical relationship between pairs of color clusters, that might indicate a temporal relationship, should involve samples that contain microbial objects whose abundances are the same or differ by one. These are situations where it is likely that one microbe is just beginning to outcompete others or the reverse. To visualize this, we constructed graphs where the samples containing objects of neighboring abundances were enlarged. For example, for a microbe m, we might enlarge samples containing m-11 and m-12. These graphs revealed classes that could evolve into one another. When a relationship is present, you see classes, particularly neighboring classes, containing large populations of enlarged nodes.

Another indication of a relationship is when you see a large population of a particular microbe’s object (i.e., only one abundance) in two nearby classes. In this case, the actual abundance differences between samples containing the object are not enough to cross the boundary of the logarithmic bin but small changes in the abundances of minor objects are enough to change the class of many samples.

Last, if there were not sufficient data to find large numbers of samples with unit or zero changes in abundance, we constructed the following statistic which has lower abundance resolution (i.e., larger abundance standard deviation). Specifically, we looked at correlations between ranges of abundance greater than 1 for specific bacteria to find these weaker relationships.

Overall, we are looking for a small change or no abundance bin change so that it is sensible to conclude that ecosystems underlying the microbiome of one class could have evolved to another.For shorthand, we will refer to these patterns as the microbe having *pronounced dynamics*.

We show results for the highly occurring species of the *Propionibacterium, Acinetobacter* and *Comamonas* genera as well as the low-count high-abundance objects referred to as M+. These had the most pronounced dynamics. It is difficult to see other less frequently occurring object’s populations change as the prevalences are too low to be statistically significant, usually less than 5 samples per class. Thus, while some of these microbes may be important, understanding them will require larger sample sizes.

###### Class Relationship Details

See Figure 14 and Table 5.

**Figure 14:**
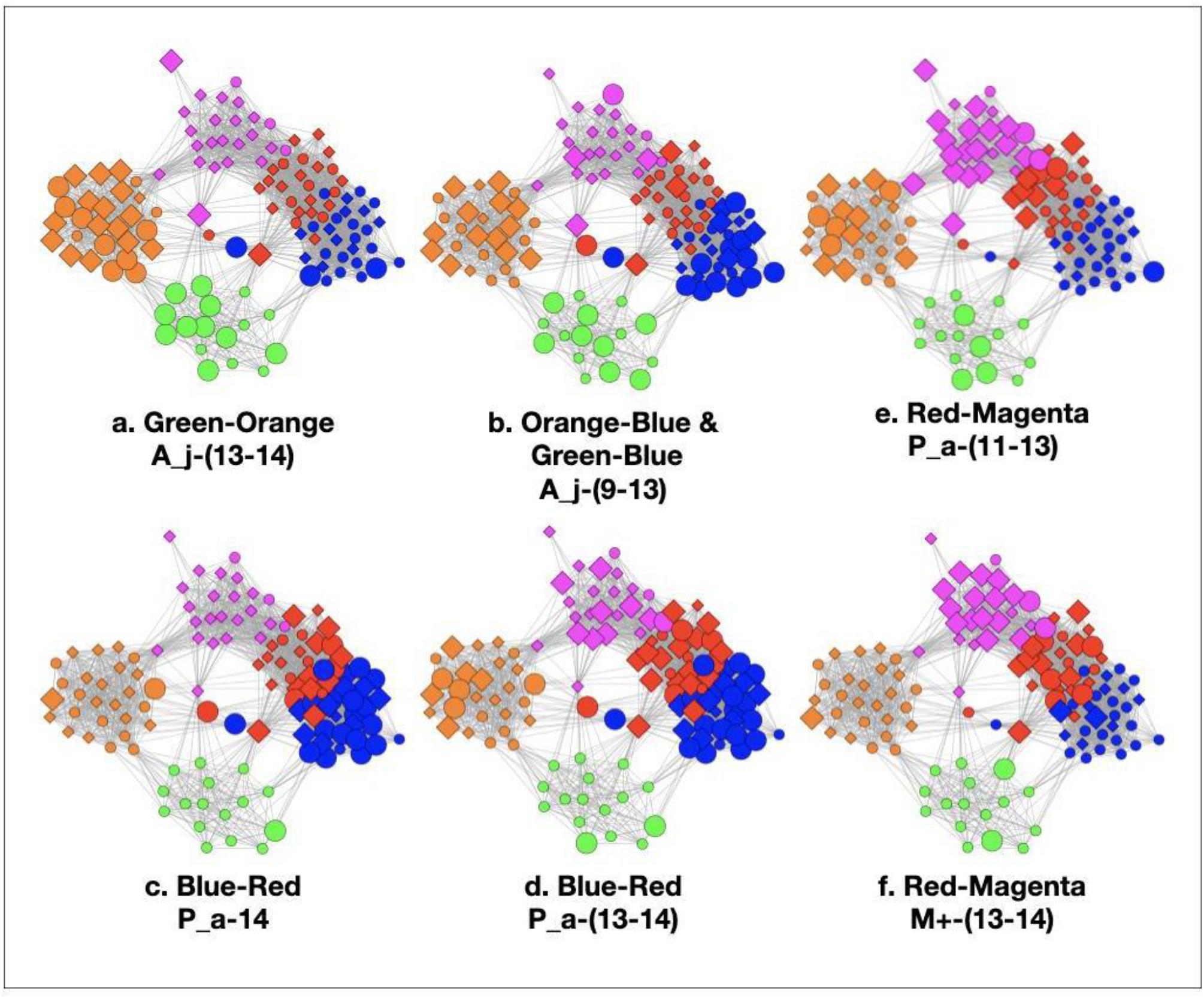
Color pair relationships. (a) Green-Orange, (b) Orange-Blue and Green-Blue, (c) Blue-Red, (d) Red-Magenta, (e) Red-Magenta.

###### Green-Orange

This pair has *A. junii-(13-14).* In this case, both objects are found in both the green and orange samples. This distribution comprises 10/16 green nodes and 23/27 orange nodes. The dominance of these abundance levels in each class suggests a relationship of green and orange. The presence of a peak of *P. acnes* at 11 in green and 13 in orange is another indicator of a relationship but it is greater than one so we will not focus on it. It is to be noted that this difference does not seem to be enough to have resulted in a diminishment of the *A. junii* abundances because of the wide bin structure.

###### Orange-Blue and Green-Blue

These two relationships are a little more difficult to characterize and we will use the wide abundance range statistic for their analyses. By enlarging samples with *A. junii*-(9-13) objects, we see high populations in green, orange, and blue. For the orange-blue, *A. junii* has a peak at 13-14 and blue has a broad wide distribution. The green-blue relationship is similar albeit with a smaller peak at 13-14 in the green. The *P. acnes* abundance also helps clarify these relationships as its average abundance in blue is much larger than in both orange and green. Of these two possible relationships, we are more confident with the orange-blue as the peak for *P. acnes* is 13 in orange and 14 in blue, a change of 1, while there is no pronounced *P. acnes* peak in the green, requiring a very large change to go from green to blue.

###### Blue-Red

This relationship can be seen in Figure 14 (c) & (d). In (c) there is a commonality between samples with *P. acnes-14* in blue and the lower part of the red cluster. This is a change of zero. In (d), we show 14-13 together demonstrating a stronger relationship. We will ignore the orange samples that show up for now because it turns out the *P. acnes* goes up and then down by class. The orange samples are on the ascendant part of the curve and the blue, red, and magenta are on the descendant side. We discuss this in more detail below. To drive home this relationship, the *P. acnes* abundance statistics are nearly the same in blue (twenty-seven 14s) and red (fourteen 14s) with a small population of P-13 objects (six 13s). There is an additional correlation provided by *A. junii* that it is widely distributed in both classes, 9-13 in blue and 10-13 in red such that they overlap. This correspondence, however, is weak as there is only one instance where the bin counts are ≥ 5. At any rate, whether we use the vary by zero or one criterion, we can see a strong blue-red relationship.

###### Red-Magenta

We use three examples to demonstrate this relationship, one with *P. acnes*, one with M+, and the last with *A. junii*. The results are in Figure 14 (e) and (f). For *P. acnes* in magenta, there are nine 13s and seven 12s suggesting a strong relationship to red where the abundance level shifts by one or two from the red distribution of 14s and 13s which can be seen in (c) and (d). The red 13s are mainly in the upper part of the red cluster closest to magenta and the magenta 13s are throughout. There are about the same number of instances of *A. junii* over a wide distribution, 4 in red and 5 in magenta. When we use the distribution of M+- (13-14) bacteria in (f) we can see a striking correspondence between the red and magenta *P. acnes* distributions. While the M+ are found in every class, it is only at abundances of 13-14 that a concentration in two classes is seen. Collectively these data support a strong relationship between red and magenta.

###### Others

We make short arguments for why the following remaining possible pairs should not be included in constructing the time-ordered network. Refer to Table 5.

###### Green-Red and Green-Magenta

There is a large difference in the average *P. acnes* abundance.There are also extensive differences in the total population of *A. junii* between classes and no *Comamonas spp.* were observed in the red and magenta classes.

Orange-Red. Since orange and blue are related, and blue and red are related, there could be a relationship between orange and red, but it is not as strong as with blue because the *A. junii* differences between orange and red are quite large.

###### Orange-Magenta

It might seem reasonable to make this argument as the orange *P. acnes* distribution is similar to magenta’s, both peaking at 13; however, the *A. junii* connection is not as strong as with blue. Additional arguments come below where we argue that over time, *P. acnes* first rises then falls. The orange 13s can then not be associated with the magenta 13s because orange is on the ascending part of the curve and magenta is on the descending part.

###### Blue-Magenta

We are going to rule out this relationship as we have already provided ample evidence that red is an intermediate class between blue and magenta. In short, the *P. acnes* abundance is 14 in blue, between 13 and 14 in red, and 13 and under in magenta.

###### Time Ordering of Classes

Now that we have established relationships between pairs of classes, it is straightforward to order them in time. (See Figure 15). To do this, we need a beginning and an end which is provided by AD statistics and the reasonable assumption that health precedes disease. Earlier, we cautioned about the use of these statistics because it does not necessarily follow that the bacteria in all of the samples of an AD subject are necessarily responsible for the disease. This is most likely not the case, however, for the green and magenta classes.

**Figure 15:**
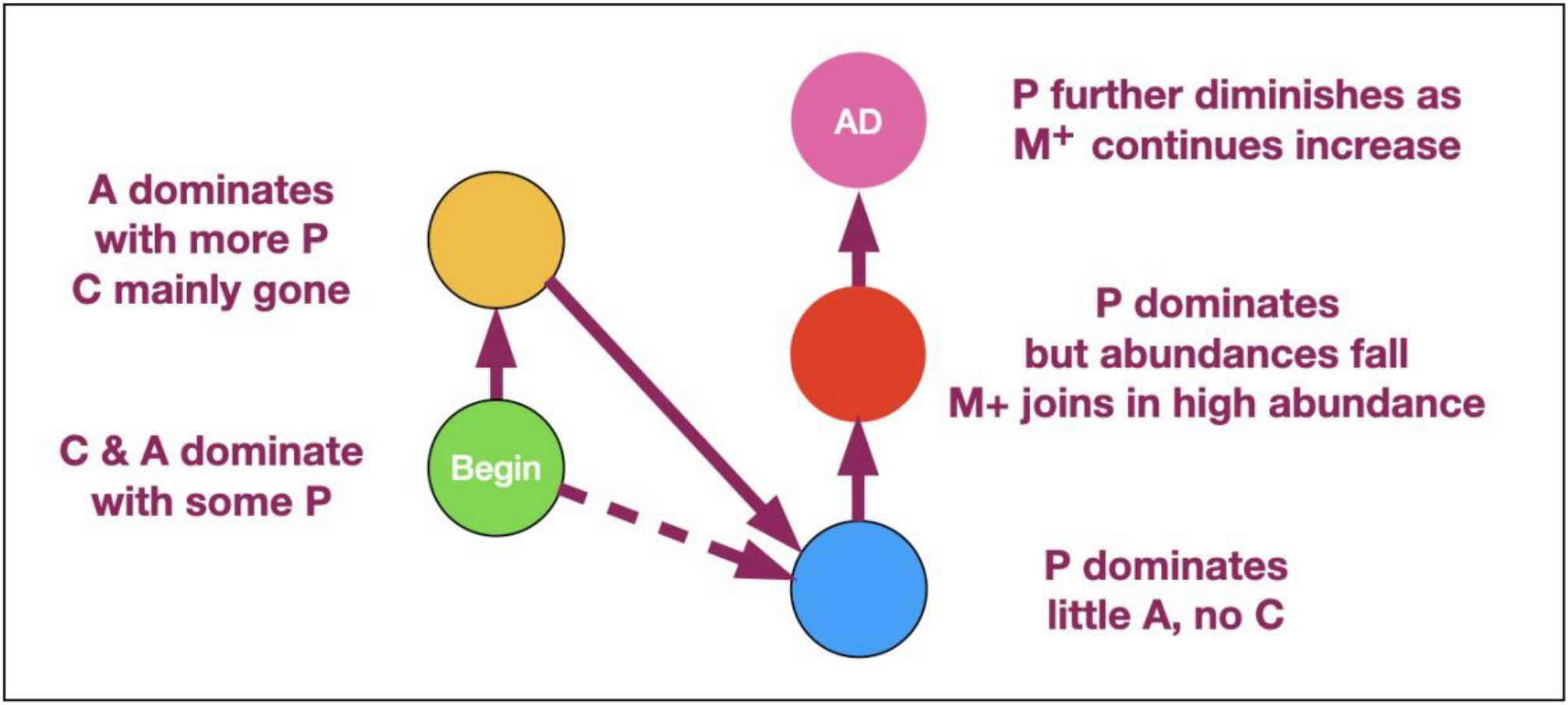
Temporal network of classes.

When the class statistic is either close to 0% or close to 100%, we are looking at situations where the population either never came from a diseased subject (green) or almost always came from a diseased subject (magenta). In these cases, the former would most likely not be pathogenic, or the subject would have AD. As the latter is almost always associated with disease, it is a reasonable hypothesis that it is pathogenic. Given that all the other sample colors are associated with both AD and controls, unless the physical sampling of the subject brains somehow missed a part of the brain that contained other pathogenic ecosystems not in any of our data, the magenta class most likely contains pathogenic ecosystems.

As a reminder, the relationships we are working with are: green-orange, orange-blue, green-blue, blue-red and red-magenta.

Refer to Figure 14. The only strong relationship to green is orange although there might be a minor link to blue (shown as a dotted line). At the other end, the only relationship to magenta is red. The only one we are left with is blue to red which must go in between the former two. Now we have a time-ordered class network. In the discussion section, we will use this network, in part, to determine the structure of the ecosystems that underlie the classes, which we will now call ecosystem mixtures. An ecosystem mixture is essentially the same thing as a class microbiome but with the added information of the ecosystem spatial structure.

###### Microbial Dynamics and *P. acnes* Anti-Correlation

At this point, we observe that the network essentially reveals temporal dynamics for all of the principal bacteria, but most importantly, it has revealed the dynamics of *P. acnes*. It begins in the green class with a low average abundance and a broad distribution. It evolves in the orange to a medium average abundance that is still quite broad. In the blue class, it reaches a high narrow peak. In the red class the abundance begins to fall and broaden. It falls further in the magenta class to a medium level with an even broader distribution.

In the green class, *A. junii* displays a medium average and narrow width. It stays roughly the same in the orange class. It then rapidly diminishes in blue and continues its decline to low abundance for red and magenta. The *C. jiangduensis* dynamic is more pronounced. It begins with a medium average abundance and width in green and then drops precipitously and broadens in orange. It is essentially not present in blue, red or magenta.

On closer inspection, there seems to be another inter-object dynamic, an anti-correlation with *P. acnes* as seen in Table 5. In the green, *P. acnes* is either non-existent, as seen in half the samples or at very low abundances. Conversely this is the class where the *Comamonas* and *Acinetobacter*species have the highest abundances and are the most prevalent. The orange class has the next lowest level of *P. acnes* where it is present in almost all of the samples but with only a small peak in abundance (at 13) observed with seven samples and displaying a very wide distribution ranging from 7-14. Curiously, there is virtually no *Comamonas,* but *Acinetobacter* persists at high levels.

In the blue class, the levels of *P. acnes* are the highest among all of the classes. There is virtually no *Comamonas* and *Acinetobacter junii* levels are low; the latter occurring in only about half the samples with a very wide abundance distribution from 9-13. In the red class, there is a high level of *P. acnes* but *Acinetobacter junii* is further diminished being present in less than a quarter of the samples over a wide abundance distribution of 10-13. Even with somewhat lower levels of *P. acnes* in magenta compared to red, *Acinetobacter junii* is still only present in fewer than a quarter of the samples over a distribution from 12-14.

Thus overall, it appears that when *P. acnes* is not present or is present only at low levels, we observe both *Comamonas spp.* and *Acinetobacter spp*. However, as the abundance of *P. acnes*increases, first the *Comamonas spp.* is lost and then the *Acinetobacter spp*. As the *P. acnes* abundance decreases from the red class to the magenta class, a new nonspecific dynamic enters the mix. The high abundance low count microbes, M+, appear mainly in the upper red and magenta classes. They increase in the magenta class as the *P. acnes* abundance falls resulting in an antiparallel dynamic. There is something very curious here that we will take up again in discussion of the ecosystems and the biology. The *P. acnes* distributions in the magenta class look similar to those in the orange class but there are major differences between the classes otherwise. The orange has a lot of *Acinetobacter spp.* while magenta does not, and the magenta has a lot of M+ whereas the orange class does not. Clearly the ecosystem evolution that goes along with *P. acnes* rise and fall is neither the same nor reversible. Time could provide the explanation. Some of the more prevalent M+ are present at low abundances and prevalence in the earliest stages, orange and green. (See Table 5). Curiously, there is only one instance of Methylobacterium in orange and two in upper red. Given long enough, however, many seem to be able to take over, even if *P. acnes* is present, either by increasing from earlier lower levels, or coming onto the scene later, like Methylobacterium.

Overall, these findings will constrain the underlying ecosystem structure that is presented in the discussion section.

##### Theme 4 - Pathogenicity of Classes

###### Disease State Statistics

The disease state statistics of each class, which summarize the fraction of samples from AD subjects, provide the first clues about whether a particular microbiome is pathogenic or not and we have used these clues to set the temporal order of classes. However, as we pointed out earlier, these numbers are not estimates of the actual pathogenicities of the classes for every class. These statistics summarize the fraction of samples from a class that come from a subject who has AD.

If it were assumed that class AD statistics were an estimate of class pathogenicity, this would be tantamount to assuming a stochastic pathogenicity mechanism where sometimes a microbiome is responsible for AD and sometimes not. It might be reasonable if we were observing AD-control mixes of 80-20 or even 70-30 where we might be able to speculate that they derived from individual differences such as immune system capability. The AD statistics for blue, orange and red were, however, just too close to 50-50 for comfort. Further, since we were unable to split the color clusters through parameter adjustment, we were confident in their robustness and therefore their microbiomes. So, we needed to find a more parsimonious explanation than accepting stochastic pathogenicity. We found one in the subject color class statistics. It is important to remember that the class results emerge from an analysis of bacterial data only. The disease statistics of Table 6 are a statement of results only, i.e. which samples in each class came from AD or control brains and are not computed by fitting an *a priori* hypothesis.

**Table 6:** Percentage of Samples that come from AD subjects by class,

| AD Statistics By Class | Percent |
| --- | --- |
| Green | 0% |
| Orange | 59% |
| Blue | 34% |
| Red | 65% |
| Magenta | 88% |

###### Subject Color Class Statistics

Because the disease state variable is an attribute of a subject, not a sample, it was necessary to relate the subjects’ sample color class distributions to disease state in order to find a relationship between class and AD. In Figure 16, we display the color class of each sample for all the subjects.

**Figure 16:**
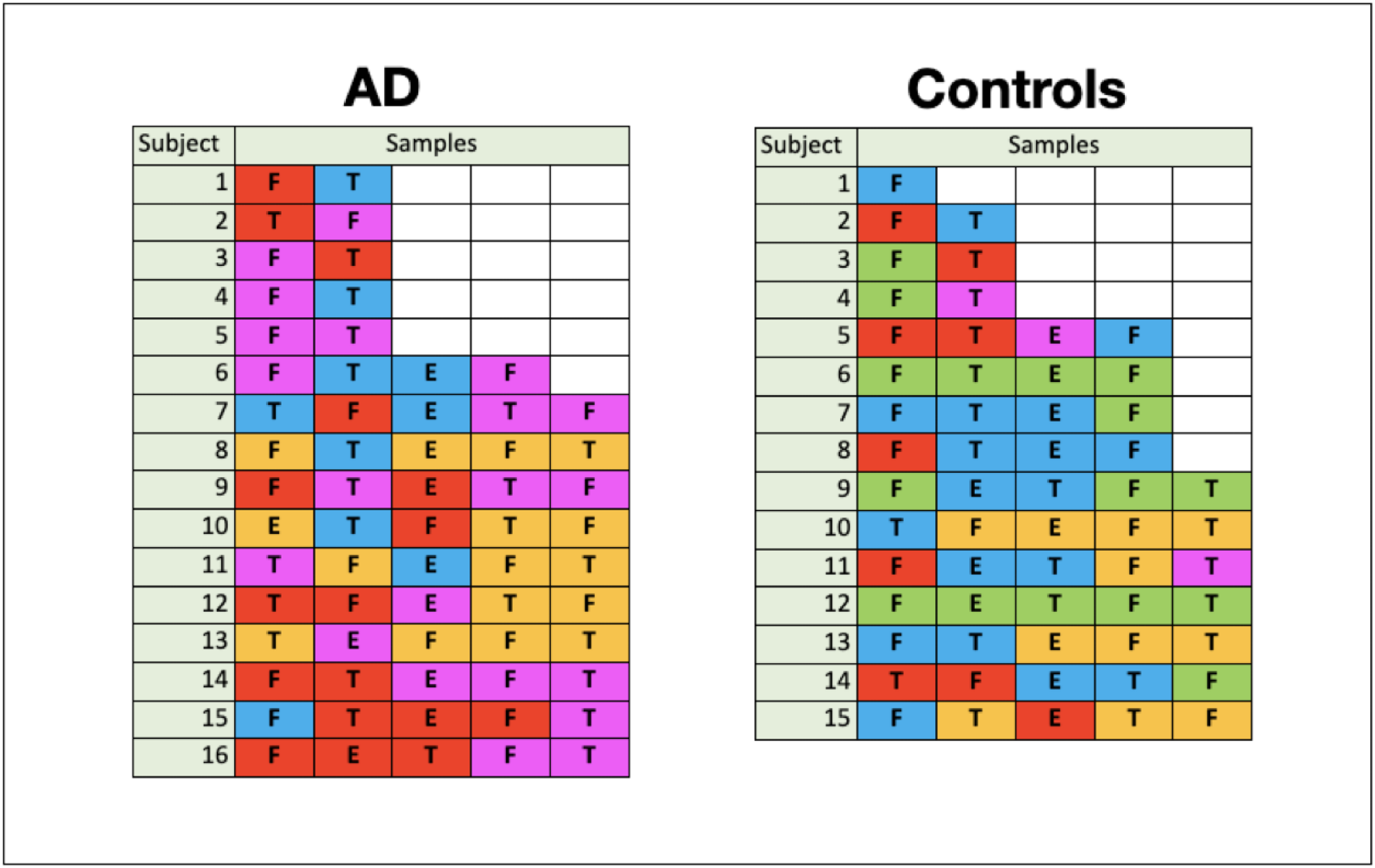
Color Class of Samples by Subject.

##### A glance at this figure indicates that the occurrence of a single magenta sample accounts for a subject’s AD in almost all cases

This suggests that the ecosystems underlying the other classes are not pathogenic even though many of their samples come from subjects who have AD. In order to do a more rigorous computation, we defined a subject level classification as the normalized sum of the classifications of each of its samples. The sum was unweighted as we had no *a priori* way to assign weights. Using the resultant mixture vectors as independent variables in a logit regression with a cutoff of 0.5 [87], we were able to obtain a high accuracy prediction (about 88%) of AD or lack of AD. True Positive, False Positive, True Negative, False Negative rates were found to be 88%, 13%, 87% and 13% respectively.

The values of coefficients for magenta and green are large and opposite in sign and the others are much smaller, again supporting what can be seen in Figure 16, that the ecosystems underlying all but the magenta class are likely not pathogenic. The coefficients were: (20.0, −3.86, 2.21, – 14.9, −31.4) for (green, orange, blue, red, magenta). This raises an issue that will have to be addressed with more research and that is whether the AD statistics for the orange, blue and red classes mainly reflect how large the regions of the AD subject are that are dominated by non-pathogenic ecosystems. A new set of measurements with a similar N could yield quite different AD statistics for the orange, blue and red classes because under-sampled statistics have very large variances.

###### Green-Orange Anomaly. (subject-level effect)

There are no subjects that have samples in the green class that also have samples in the orange class. In other words, green and orange samples do not appear together in the same subject. This is true both for AD and control subjects. We are going to assume that this is not a statistical fluke due to under-sampling of subjects and look for explanations. On the other **hand, for other control subjects** where orange is not present, we find green occurring with blue, red, and magenta samples. In all these cases but one, however, there is only one green sample in the subject. We will explore the meaning of these results in the discussion section where we utilize arguments based on ecosystem structure and infection mechanisms.

###### The *A. junii-14* to *A. junii*-13 Orange Class Transition

In Figure 17, it can be observed that as the *A. junii* objects are varied from 14 to 13, enlarged nodes are seen that are mainly from control samples at 14 and mainly from AD samples at 13. This suggests that there is a dynamic that relates *A. junii* abundance to AD that may not be consistent with our Theme 3 results that indicate that orange is not pathogenic. We will attempt to resolve this inconsistency after we discuss the structure of large-scale spatial correlations in the discussion section. Interestingly, the A. *tjernbergiae* species does not have this dynamic. In going from low to high abundance, this species jumps from green to orange around level 12 but unlike *A. junii,* there are roughly the same number of samples with each disease state at each level over 12.

**Figure 17:**
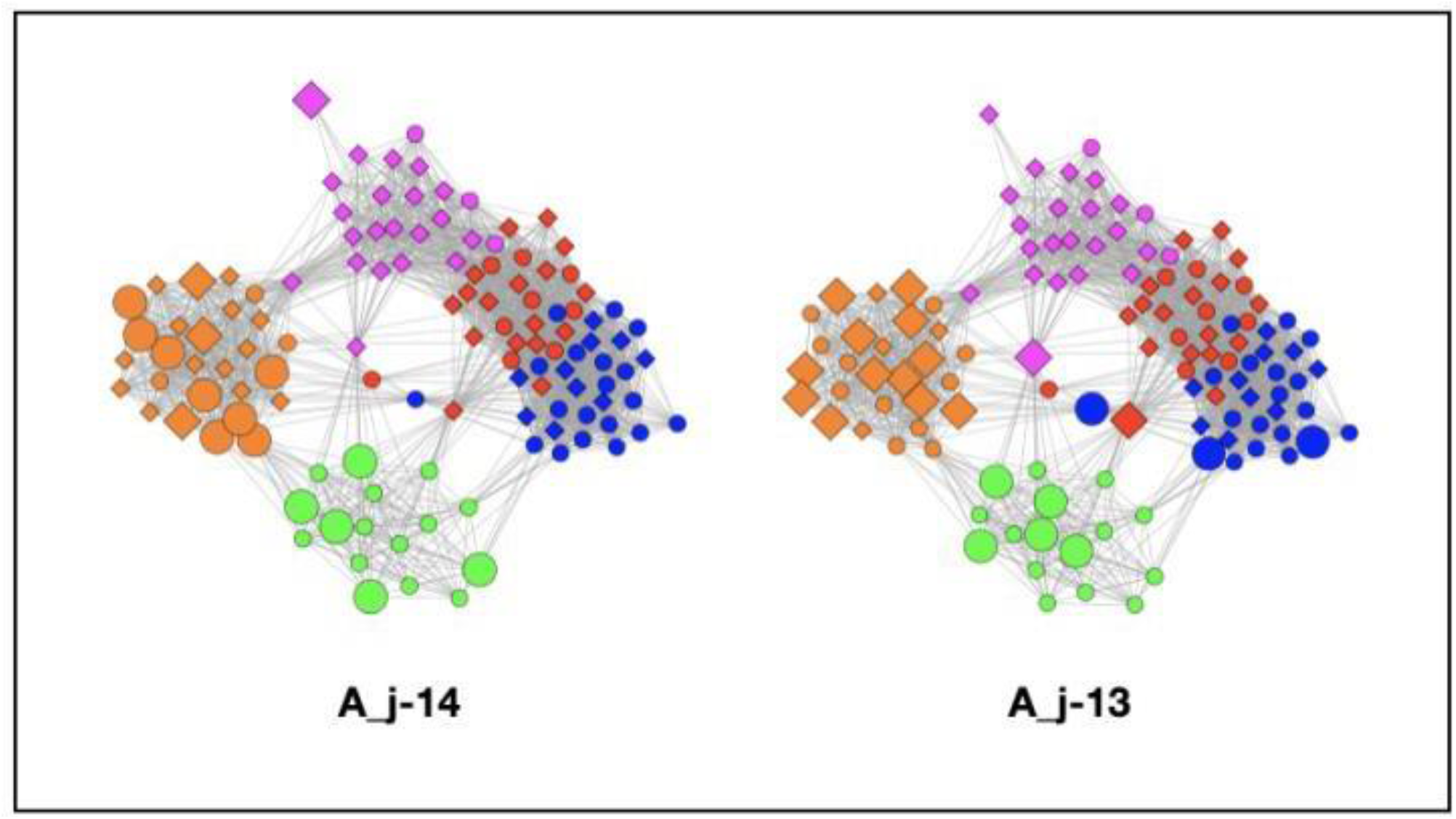
A-14 to A-13 transition.

###### Location

There is insufficient data to say anything definitive about location, but we offer one set of results. There is a hint that when a magenta infection occurs in an AD subject, it is likely to be found at least in the frontal lobe. Out of 13 AD subjects with magenta infections, 9 have at least one frontal lobe magenta sample, while only 2 have temporal lobe magenta samples without a frontal lobe magenta sample. Of these 13 AD subjects, 5 have temporal lobe magenta samples that also have magenta frontal lobe infections. For the 3 subjects that had samples in the entorhinal cortex, only 1 had samples in the temporal or frontal cortex and, in this case, it was both. The other 2 had neither.

***So, at least with this small set of data, there is a suggestion that subjects with the magenta class in their frontal lobe are more likely to have the diminished cognitive abilities that result in AD diagnoses.***

## DISCUSSION

The existence of a human brain microbiome has been suggested recently and a dysbiotic brain microbiome could contribute to AD pathogenesis. The use of long-read sequencing of full-length 16S rRNA genes allowed us to profile the bacterial composition of human brain tissue samples from AD and non-demented control subjects as well as to evidence potential complex polymicrobial interactions. The MLDA computations provide us with a rich set of results that allow us to construct a simple microscopic and macroscopic theory of the ecosystems underlying the bacterial abundance classes and their temporal and pathogenic relationships to AD. In this section, we will interpret the results of the computations and speculate how Alzheimer’s disease may develop because of dynamic and evolving bacterial ecosystems in the brain. We do not assume that there are no other microbial factors or even non-microbial factors that are simultaneously involved.

The MLDA computations allow us to: (1) characterize a macroscopic spatial distribution of the ecosystems, (2) propose a microscopic ecosystem structure on the cellular scale, (3) relate these structures to the temporal ordering of classes, and (4) speculate how the magenta ecosystem structure could be pathogenic. We further suggest ways that bacteria may have invaded and spread in ways that are qualitatively consistent with our statistical results. Together, these will comprise a rudimentary bacterial etiology of AD.

### Ecosystem Mixtures - From the Microscopic to Macroscopic

Since the sampling process sums and averages the bacterial load over the sample volume, the sum does not tell us about microscopic structure within a single sample. There could be ecosystems inside human cells, between the cells, within capillaries, etc., each with different bacterial abundances. The abundances could be dynamic and have different abundance equilibria. The same bacteria may be in more than one ecosystem. In other words, there are lots of possibilities, with some even persisting and changing postmortem. The sampling process, however, just sums them all giving us a set of total abundances for each sample.

In other words, a class microbiome comes from a sum of the ecosystem mixtures. These classified mixtures are what we have at this point. While we do not have enough data to determine the underlying ecosystems precisely, these results give us enough to identify several important features. Given that MLDA has identified 5 sets of samples with similar microbiomes, we will try to use the differences among samples *within* a class to infer features of the underlying ecosystem structure.

These sample differences are manifested in the results in many ways. We will focus on the abundance distributions of the principal bacteria and their occurrence within each class to reveal the underlying spatial structure. We also postulate several scenarios and then use the results to infer the likeliest among them.

Since the results already tell us that the magenta class is associated with AD and its lack mainly not associated with AD, the underlying structure of this class should provide additional information in regard to how its bacterial ecosystems could be causing AD.

At the other end of the spatial scale, we wish to understand how the classes of ecosystem mixtures are spatially distributed in the brain. We know that the brains of AD patients manifest large spatial scale changes accompanied by amyloid plaques and neurofibrillary tangles. So, we wish to determine if the color clusters have a relationship to this physiological structure. We will examine color class occurrence within subjects to look for such patterns.

### Useful Statistics and Their Meaning

In all the following arguments, we wish to emphasize that we are making qualitative and not rigorous statistical arguments as our sample size does not permit it at this point.

Despite not directly observing ecosystems at the microscopic scale, the sample abundance average and distribution statistics within each class can reveal characteristics of the microscopic structure. It is useful to recall that the number of samples in each class are: green: 16, orange: 27, blue: 29, red: 26, and magenta: 24. The computational results effectively break samples into many buckets defined by class, dominant microbes, subject disease state, etc. We use the counts of samples in various *ad hoc* buckets to make our arguments. Generally, these buckets do not exceed 15 counts and are often less than five.

Consequently, no matter what the argument, we are faced with the challenge of making arguments on statistically shaky grounds. Sometimes, we may sum over abundance or bacteria to improve significance, understanding that it is at the expense of bacterial specificity or abundance resolution. Despite the lack of statistical rigor, we nonetheless believe that the number of patterns we are able to discern fit together into a simple framework that is not likely to have occurred by chance and which suggests many novel hypotheses.

#### Cellular Scale Microscopic Structure

##### First Steps

The first step toward understanding the microscopic structures at the root of class is to summarize which objects appear the most often in samples, grouped by their largest class component (i.e. color). These are listed in Table S3, Table S4, Table S5 and Table S8. in various levels of detail. The classes are dominated by three genera: *Propionibacterium (P), Acinetobacter (A)* and *Comamonas (C)* and, in particular, three species, *P. acnes, A. junii* and *C. jiangduensis.* By ‘dominate’, we mean that a microbe has both high average abundance among samples of the class that contain it and that it occurs with high frequency within the samples of a class. We may also say that it has a high prevalence within the samples of a class.

We need to determine if the three dominant microbes are part of one or more ecosystems to understand the dynamics of the system. If two microbes are in the same ecosystem, then we can assume that their average distance is on the scale of the microbe size. On the other hand, if they are in separate ecosystems, we can assume that they are separated by distances on the scale of human cells because different ecosystems may occupy niches defined by different nutrients and environments that likely vary on the cellular scale. Overall, we are trying to determine the microscopic and macroscopic spatial distributions of these principal bacteria, but we have to remember that MLDA does not directly say anything about spatial distributions. We will be hypothesizing various spatial distribution scenarios and evaluating whether they are consistent with the MLDA results.

##### Spatial Sampling - Microscopic Point of View

To begin with, we will assume that classes represent different mixtures of ecosystems that the MLDA algorithm separates and that it is reasonable to apply the sampling principle of Theme 2 on the results. A physical sample is the weighted sum of these class mixtures where the weights are the class components. The principle allows us to model a single physical sample as if it were the separate sampling of each ecosystem mixture. We will call this virtual sampling. Even so, the output of the MLDA is still in terms of the measured microbial objects which are sample averages with *no* spatial ecosystem information. (See for example Table S3, Table S4, Table S5, and Table S8).

The grouping of the raw data of the principal microbes by maximum class component (color) in Table 5 provides a way of testing several ecosystem scenarios. Our arguments will assume that it is good enough to model the sample as if it contained a single component defined by its color (maximum component) and that the color’s microbiome is defined by the set of microbial objects contained in samples of that color. This is reasonable given that the maximum class components in most samples are high (see Figure 9).

Three possible ecosystem spatial scenarios are discussed below. We will make qualitative statisticalarguments about the reasonableness of each one. Rigorous statistics with the small amount of data would be questionable. We will evaluate each scenario in terms of the sample microbial averages and qualitative widths of the logarithmic microbial abundance distributions.

##### Scenario 1 - Microscopically Interleaved Ecosystems

This scenario assumes that there are distinct ecosystems that exist at the human cellular scale, each being dominated by one of the principal/dominant bacteria. Refer to Figure 18. One can visualize them as pixels that aggregate to a sample. Each pixel represents a *Propionibacterium, Acinetobacter* or a *Comamonas* dominated ecosystem. Each class could have different densities and homogeneities of each of these ecosystems within the brain. We have in mind a random marble in a bucket type of model. In some cases, one pixel may be spread over the sample area with a random scattering of one or two of the others. In other cases, there may be two with similar densities and a scattering of the third. The scattering densities vary by class. In general, low densities imply a larger average distance between ecosystems and high densities imply a smaller distance (Figure 19). Add up the number of ecosystem pixels within a black circled sample as an estimate of the measured abundance of the raw data in a particular class. In the end, the densities and homogeneities of each ecosystem pixel will determine the abundance averages and distributions within the class. Lower density pixels will tend to be less homogeneous and have wider abundance distributions than higher density pixels. *Note our use of ‘density’ to describe actual biological distributions as opposed to ‘abundance’ which we use to describe the fractional amount of a bacterium within a macroscopic sample.*

**Figure 18:**
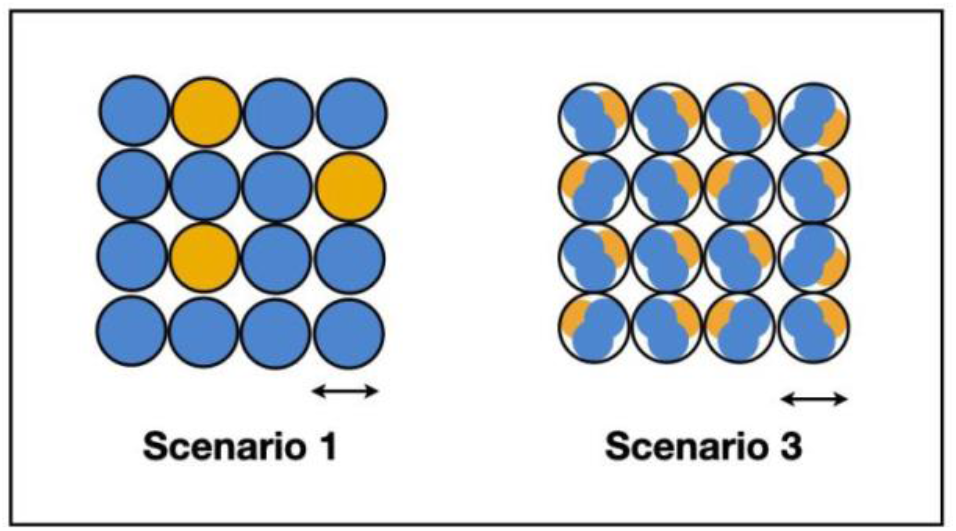
This figure shows possible ecosystem structure at the microscopic scale. The arrows roughly indicate the human cellular scale. Scenario 1 suggests ecosystems dominated by one principal bacterium predominate around a particular cell while Scenario 3 suggests that ecosystems comprised by multiple principal bacteria predominate around a particular cell. A physical sample would comprise all or large fractions of the above arrays.

**Figure 19:**
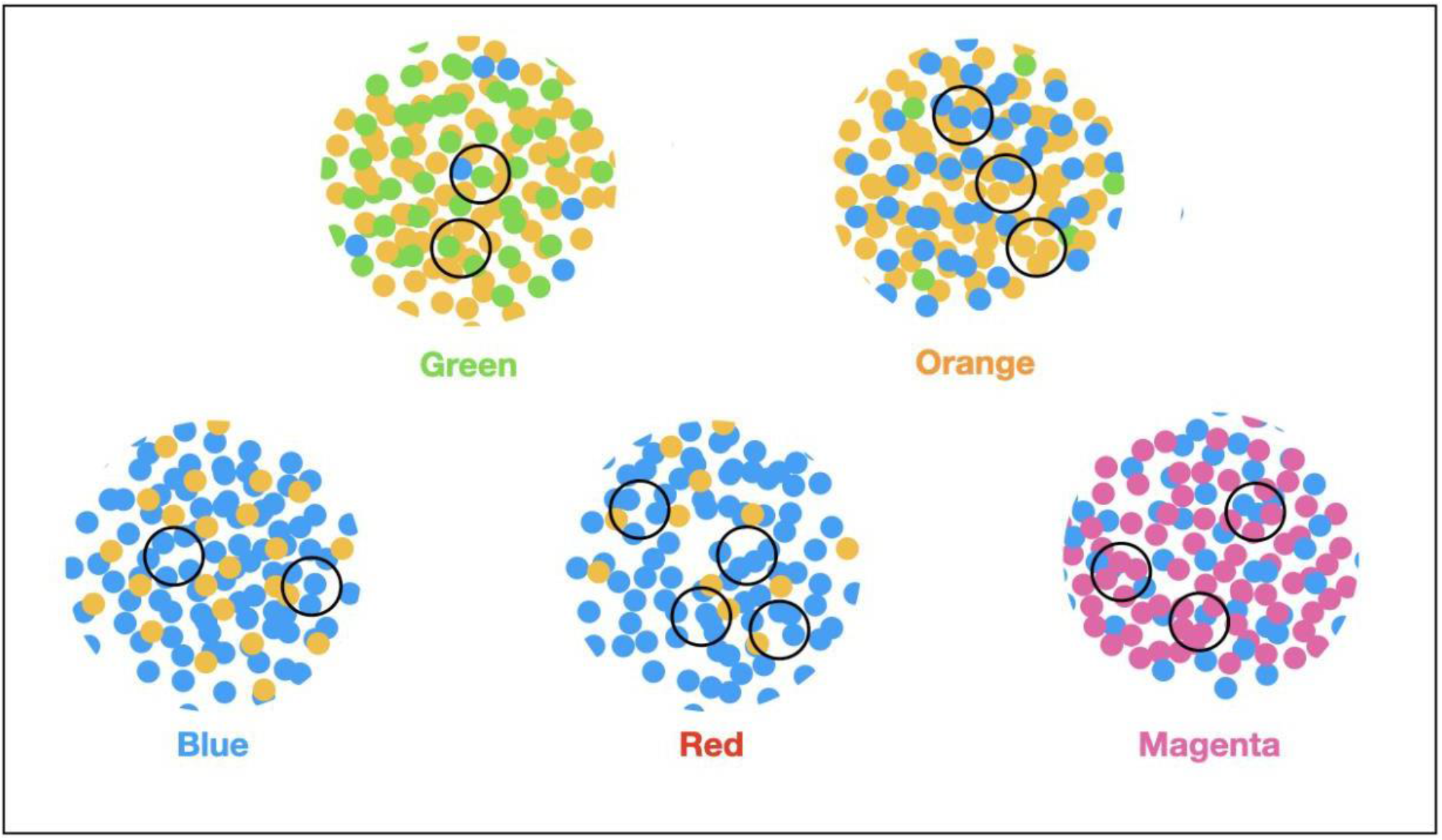
Idealized depiction of distribution of ecosystems. Each dot is an ecosystem dominated by a particular species: blue for *P. acnes,* orange for *A. junii*, green for *C. jiangduensis* and magenta for M+. The large circles are class mixtures also labeled by colors. The small black circles depict samples. Green is dominated by *C. jiangduensis*; blue is dominated by *P. acnes*; orange by *A. junii*, and magenta by the M+ set.

Given that the sample to human cell size ratio is on the order of 50X, it is possible to qualitatively reproduce the class raw data, by experimenting with different densities and homogeneities of the pixel distributions.

##### Scenario 2 - Sample Area Structure

Scenario 2 differs from Scenario 1 in that the ecosystems are not randomly mixed but fall within relatively homogeneous areas. For this scenario to work, samples have to be likely to straddle regions which would only happen if the regions are on the scale of the sample or smaller. The smaller the regions are, the more this scenario approximates Scenario 1 so it does not seem like a good candidate.

##### Scenario 3 - Single Ecosystem Model

The third scenario assumes that P, A, C and M+ microbes comprise a single ecosystem of interacting bacteria on a bacterial scale. We have visualized this in Figure 18 as multicolored dots of P(blue) and A(orange) within the human cellular scale to suggest the bacterial scale. Also see Supplemental Table S6.

Each class has different P, A, C and M+ objects in its samples. There is P and A in most of the classes and there is P, A and C in the orange and green classes. Some of the M+ can be found in all classes. If this scenario is the case, then this ecosystem would be comprised of a competitive system where the system sometimes has one bacterium dominating and sometimes another. Physical samples would simply average over these random competition phase differences. The existence of the clusters and temporal class order makes this unlikely. Clusters occur when you have mainly high levels of one class component within most samples. This scenario would have all levels within the pixels averaging to a medium level over the sample. For this not to be the case, the pixels would have to be spatially correlated so that far away pixels always had the same species dominate at the same time within a pixel. Even if we forget about the pixels, it is hard to come up with random processes that account for some of the major features we see such as the green-yellow exclusion and the *P. acnes* dynamics we have described and which we explain below using Scenario 1.

Further, we do not think that this model can easily account for the abundance variances in the easy way that the first scenario does. Another problem is that it cannot account for the absence or near absence of *Comamonas* in everywhere but the green class which is more fully discussed below.

##### Idealized Model of Ecosystem Mixtures by Class

In the specific class descriptions below in terms of Scenario 1, we use the logarithmic abundance bin values to discuss average abundances and their widths. It is possible to argue that these bins are too coarse. We will show, however, that it does suffice, though, to get a qualitative sense for the likely ecosystem structure for each of the classes. At the beginning of this paper, we argued that the precision of the sequencing was probably beyond what was needed to understand the biology. Being able to describe Scenario 1 by using the reduced abundance resolution of the logarithmic bins supports this argument.

In Figure 19, each large circular area is an idealized area of the brain representing a pure ecosystem mixture class. Each smaller black circle is a physical sample. Each dot is an ecosystem with a particular dominating microbe: blue for *P. acnes*, orange for *A. junii*, green for *C. jiangduensis* and magenta for M+. The number of dots within a sample is a qualitative way of estimating the abundance of the principal microbe. By showing gaps in some of the spatial distributions, we are trying to create a visualization of inhomogeneity. Below, we describe each of the elements of the figure primarily in the context of scenario 1 (see Table 5).

###### Green

The green class has *C. jiangduensis* and *A. junii* as its dominant microbes. There is a lot of *C. jiangduensis* in the green class but almost none in the orange class which also shares both species of *Acinetobacter* with green. Green presents with a narrow *C. jiangduensis* distribution of 14s and 13s. *A. junii* has a narrow distribution of 13s and 14s. *P. acnes* on the other hand, has a wider distribution peaking at 11 with a width of several abundance units and is only present in half of the samples at levels ≥ 7. These results suggest an ecosystem structure where *C. jiangduensis* and *Acinetobacter spp.* are interleaved with one another with a random scattering of *P. acnes* at lower density than either *C. jiangduensis or A. junii*. This density should be low enough so that there is a high probability that some samples from a green region will not contain *P. acnes* as observed.

###### Orange

The orange class has a narrow distribution of *A. junii* in 13s and 14s that it shares with green. In comparison to green, the *P. acnes* distribution is narrower with a pronounced peak at 13 and a presence in nearly all samples. *C. jiangduensis* has a wide distribution but is present in less than one third of the samples in this class and has no samples with abundances at 13 or 14. Compared to the green class, it has largely disappeared.

These results suggest a density distribution dominated by *A. junii* with a significant density of *P. acnes* but with *C. jiangduensis* present only at a very low density. The fact that the *P. acnes* distribution has a width >2 and the abundance level is not the highest suggests some inhomogeneity in its spatial distribution.

###### Blue

The blue class has a very narrow distribution of *P. acnes* with a high average abundance at 14 that is representative of almost every sample of the class. *A. junii* has a wide distribution, peaking at 10, suggesting a competition where *P. acnes* has become dominant. *C. jiangduensis* is not present at all in abundances ≥ 7. Because of the narrowness of the *P. acnes* distribution, its spatial distribution is homogeneous with a light random scattering of *A. junii.*

###### Red

The red class has a somewhat wider *P. acnes* distribution than the blue class with some 13s in addition to 14s. There is again no *C. jiangduensis*. The larger width of the *P. acnes* distribution compared to the blue class indicates that the underlying spatial distribution of its ecosystems is not as homogeneous as the blue class. The *A. junii* distribution is wide but it does not occur in most of the samples suggesting it is widely spaced with significantly lower densities than in the *A. junii-*predominant green and orange classes.

###### Magenta

The magenta class has a wider *P. acnes* distribution compared to blue with a mix of 13 s, 12s and a few 11s leading to lower average abundances of this species. The *P. acnes* ecosystems are therefore not homogeneously distributed. Again, there is no *C. jiangduensis* ≥ level 7 abundance. As mentioned in the results, the M+ microbes appear along with the *P. acnes* within individual samples, but at far higher abundances. Looking back at the M+ microbes in earlier classes, we see that they have been present, however at far lower abundances with wider distributions, roughly in the 8-12 range. Importantly, in the red and magenta classes, they jump up into the 13-14 range where in most samples they overtake the *P. acnes* abundances. In overtaking *P. acnes* in the magenta class, the M+ microbes could either have increased their abundances in microscopic niches where *P. acnes* was not located, or they may have outcompeted *P. acnes* in their own microscopic niches. Alternatively, the M+ could have been part of the *P. acnes* ecosystem as suggested in Scenario 3 and somehow pushed down its density, but we believe this to be less likely for the reasons already expressed. In Figure 19, we have portrayed the M+ as separate ecosystems but additional studies will be required to test these alternative hypotheses.

##### Explanation of Green-Orange Anomaly

The green and orange classes are on the ascendant part of the *P. acnes* dynamic. Once *P. acnes* begins to spread, either within an AD subject or a control, it is widespread enough so that the *P. acnes* vs. *C. jiangduensis* anti-correlation kicks in, making it unlikely that *C. jiangduensis* exists at a level where a sample would be classified green. There is some amount of data that supports this assertion (see Supplemental Table S6). In the green class, there are 8 samples where *P. acnes*, *A. junii and C. jiangduensis* coexist. Seven of these have a *C. jiangduensis* abundance ≥ 13 and only 2 of these have a level *P. acnes* abundance > 11 with 5 having abundances ≤ 11.

At the microscopic level, when there is enough *P. acnes* to be proximate to most of the *C. jiangduensis*, there is some kind of interaction that makes *C. jiangduensis* populations not possible. The lack of samples that have microbiomes with object abundances between green and orange suggests a rapid diminishment of *C. jiangduensis* once *P. acnes* reaches a critical level. Alternatively, it could also be possible that this situation was not sampled.

Thus overall, we now have the beginnings of a theory. There seems to be consistency between the ecosystem structures we describe and the MLDA statistics. The remaining parts of the theory will tie these ideas to deeper concepts of pathogenicity, possible relationships of the ecosystems to NFTs and plaques, and entry mechanisms. We undertake that below with more details.

#### Large Scale Macroscopic Structure

Previously, we indicated that there is more than one model that could explain how the class mixtures are spatially distributed on a large scale. We know several things. First, there is a high probability of sampling a magenta mixture from an AD subject. Second, we know that the chance of sampling another color is lower among AD subjects (Figure 16). Third, we know that we have under-sampled the brain as the volume of the samples is very small compared to the entire brain or a particular part of it. Specifically, we have 1 or 2 samples per lobe per subject and the samples are very far apart. In the following analysis, we rely on the virtual brain assumption which we mentioned earlier. This assumes that we are sampling two subjects, one with AD patterns and one with control patterns. The justification for this is the clustering of samples from many subjects into the same color class. In the analysis below, we will use only the AD virtual brain as our conclusions extend to the control brain too.

We are going to look at three scenarios. The first assumes lots of small regions of homogeneous class where small means small compared to the distance between physical samples. The second assumes large regions of homogeneous class. The third scenario assumes some of both.

By looking at the statistics of class occurrence by subject, we can evaluate alternatives for how the class mixtures may be spatially distributed. We constructed distributions of the number of class occurrences by subject for each class. In other words, we counted the number of subjects that have 1 occurrence of a class, the number of subjects that have 2, etc. See Table 7.

**Table 7:** Number of class occurrences by subjects for each class for comparison with simulation of macroscopic distribution scenarios.

|  | AD |  |  |  |  |  |  | Controls |  |  |  |  |  |
| --- | --- | --- | --- | --- | --- | --- | --- | --- | --- | --- | --- | --- | --- |
|  | 1 | 2 | 3 | 4 | 5 | Tot |  | 1 | 2 | 3 | 4 | 5 | Tot |
| green | 0 | 0 | 0 | 0 | 0 | 0 | green | 4 | 0 | 1 | 1 | 1 | 7 |
| orange | 0 | 1 | 2 | 2 | 0 | 5 | orange | 1 | 0 | 2 | 1 | 0 | 4 |
| blue | 6 | 2 | 0 | 0 | 0 | 8 | blue | 5 | 4 | 2 | 0 | 0 | 11 |
| red | 5 | 3 | 2 | 0 | 0 | 10 | red | 5 | 2 | 0 | 0 | 0 | 7 |
| magenta | 7 | 4 | 2 | 0 | 0 | 13 | magenta | 3 | 0 | 0 | 0 | 0 | 3 |

In all colors and disease states but one, orange, we see a skewing towards the occurrence of one class. In the orange case, both the AD and control distributions are skewed towards a flatter broader distribution that does not include 1s.

To understand this statistic, we constructed a virtual brain to model the AD subject. Then, in a simulation, we under-sampled it like we did with the real brains, sampling 4 at a time. We compared two underlying class distributions by constructing the same statistic as we did from the real results. Keep in mind the sampled elements are spatial distributions of class mixtures not individual ecosystems so the colors we are referring to are the colors of the mixtures as in Figure 19 and the previous paragraph. To see the following models, refer to Figure 20. The first was random, like well -mixed colored marbles dropped on a surface. The second was clumped where there was a regional structure like countries on a world map where many like colored elements were next to one another. We created one large clumped structure (magenta class) and the rest of the space was randomly filled with the other colors. We did two simulations to compare these scenarios.

**Figure 20:**
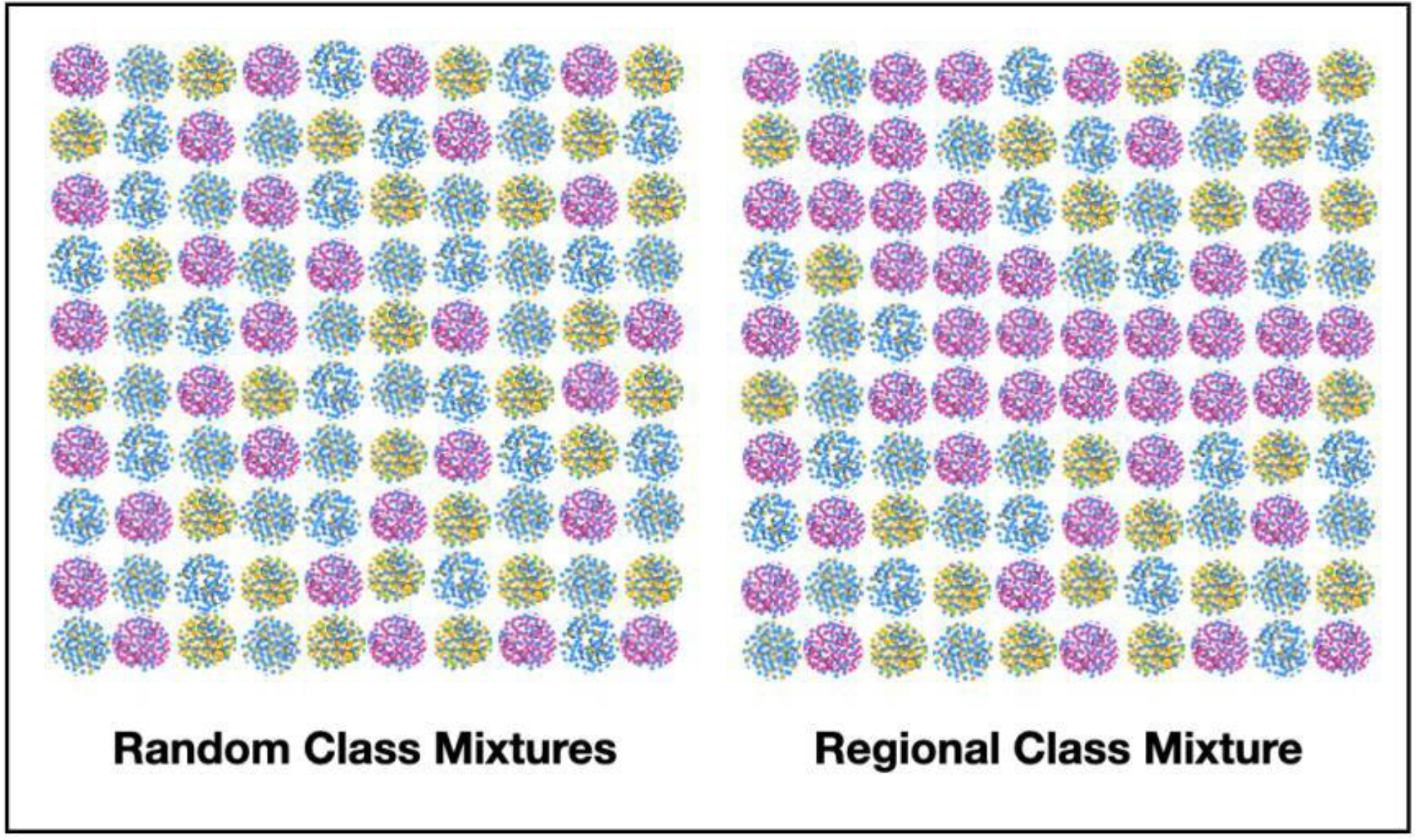
Each array represents a large area of the brain. Each element is a single class mixture like the ones from Figure 19. The size of the element could be from centimeters to several centimeters. The left-hand side produces statistics like Table 7, (except orange). The right side produces flatter statistics without 1s, like orange.

The random scenario produced results skewed towards 1 occurrence among four colors. The regional structure, once it is large enough, produces a flatter distribution that is missing 1’s and distributed over 2s, 3s and 4s. Our results, except for orange are therefore consistent with the *lack* of a regional structure. Given the lack of data, it is hard to say much more than the orange data suggests more of a regional structure.

So, given the high occurrence of magenta in the AD subjects, we can say that there is a high density of small areas in these subjects that contain the magenta microbiome interspersed with the others. In addition, if our comment about orange is right, some of the subjects might have large regions that are orange with scatterings of the other colored microbiomes.

This structure is thought to be repeated over the brain with the marbles referred to above being the color of the elements as labeled in Figure 19, not the dots. The brain might be covered by hundreds of these randomly interleaved elements.

The explanation of the occurrence of the magenta class mixtures in AD subjects is that it stems from their spatial density. As long as density of magenta class elements is high enough, there will be a high probability of sampling at least one magenta element when four samples are taken per subject, whereas the probabilities will be smaller for the other class mixtures if they have a smaller density.

There are two other important points. The first involves assuming elements are made of a single class mixture. Because the math can separate out real samples into classes (or at least samples that have >40% of one class), we can approximate the samples as coming from a particular class mixture. The second point brings us back to the origin of the abundance distributions. When eachelement is sampled, the sample size is far smaller than the element. So, the repeated samplings of the elements will reveal the inhomogeneities of the class mixtures on the microscopic scale as shown in Figure 19.

##### Explanation of A-14 to A-13 AD Transition

It has been noted that in the orange class, samples from controls mainly have *A. junii*-14 objects and samples from AD subjects almost all have *A. junii-*13 objects. In other words, controls in the orange class tend to have higher levels of *A. junii* than samples from AD subjects. This phenomenon is not likely to be causal because the orange class is not particularly associated with AD on a subject level. A more reasonable explanation could be that since AD takes time to develop, we could be seeing earlier times associated with high levels of *A. junii* and later times with lower levels of *A. junii*. For the other prevalent *Acinetobacter* species of the orange class, *A*. *tjernbergiae,* this dynamic is absent, so it does not seem to have the same time correlation with AD subjects as *A. junii*. With an abundance level < 12, *A. tjernbergiae* is green and therefore all controls. At levels > 12, this species is just as likely to come from an AD subject as a control. If the time correlation explanation is correct, perhaps the low levels of *A. tjernbergiae* comes from earlier stages and the higher levels wax and wane removing the time correlation of *A. junii*. It is hard to further investigate this hypothesis with the data available, but it is worth noting that this is another fascinating dynamic, identified by the methodology, that could be used to understand underlying mechanisms of bacterial spreading and the emergence of AD.

###### Thoughts on the Etiology of AD

The following involves a fair amount of speculation but given the correspondence with our findings, we felt that it was worth hypothesizing as an inspiration to focus future research.

###### The ubiquity of *P. acnes* and Amyloid Plaques and Neurofibrillary Ranges and Time

*P. acnes* occurs at some level in 83% of the samples, both the AD and controls, and in all classes. It occurs in over 88% of the samples not including the green class. These observations suggest that it may be interacting with all the ecosystems in each class and through these interactions plays a primary role in defining class by determining which microbes ultimately predominate. If the temporal order of classes we have argued is correct, *P. acnes* begins at low abundance as seen in the green class, it then increases in abundance in the orange class, peaking in the blue class and falling in abundance throughout the red and magenta classes as disease emerges. The fact that orange and magenta have similar average abundances strongly suggests that, as time passes, something changes, perhaps physiologically to allow the M+ species complex to dominate in the magenta rather than the ecosystems evolving back to orange. Perhaps the brain’s immune protection is diminished or a failing blood brain barrier gradually increases the microbes it lets in over time. This may be an example of multiple meanings of one object mentioned earlier. The same *P. acnes* objects in the orange and magenta classes seem to be involved in different processes.

*P. acne’* ubiquity also corresponds to another well-known observation, the ubiquity of plaques and NFTs in the brain tissue of AD subjects. The details of their spatial distribution is not within the scope of this work. Even so, we would like to suggest that the ubiquity of both is not happenstance and perhaps the plaques and tangles are some type of response to the *P. acnes*. While the two observations seem to be related, we emphasize that our results suggest that the presence of *P. acnes* alone is NOT evidence of damaged tissue that results in the observed cognitive impairment of the subjects. In other words, we are suggesting that the presence of *P. acnes* could cause plaques and NFTs but *not* AD.

The evolution of the microscopic structure of *P. acnes* ecosystems suggest that it is a driving factor in the emergence of AD, even if it does not directly cause it. Even though the *P. acnes* ecosystems are a little closer together in the orange than in the green, it is enough to eliminate the ability of *C. jiangduensis* to survive. As the concentration of *P. acnes* in ecosystems increases, the ability of *A. junii* to survive diminishes. Clearly, in the magenta class, something dramatic changes as the homogeneity of *P. acnes* declines along with its abundance.

###### Pathogenicity - Prediction of Disease State

We have argued that the only class that has a strong relationship to pathogenicity is the magenta class. Now that we have a theory for the microscopic ecosystem structure, we can speculate on how the magenta class ecosystem structure might relate to pathogenicity. There are two parts to the argument. First, given that there are many M+ bacteria from many species and genera, it is hard to argue that these are all pathogens. On the other hand, given their presence in the magenta class and their pairing with *P. acnes*-(11-13) in individual samples, this suggests that the presence of both is related to pathogenicity. From a biological point of view, this pairing suggests some type of interaction between the *P. acnes* and M+. It is not outlandish to presume that the M+ share how they communicate or compete even though they are demonstrably different species [88,89]. So, it may be that the biochemical mode of communication or other interaction between M+ and *P. acnes* directly causes AD.

###### Microbiology of Principal Bacteria

As we stated at the beginning of the paper, it is difficult to ascertain the properties of all the bacteria observed including the principal ones. We will nonetheless try to point out how some of these properties may be consistent with what our results show. We will focus on their motility and preferred pH. We will not comment on the M+ set.

*P. acnes* is not motile [90,91], while *A. junii* has twitching motility [92,93] and *C. jiangduensis* is motile [94]. *P. acnes*’ lack of motility suggests that there must be a mechanism for its ubiquity other than the ability to move. Perhaps it gains access through the capillaries of the blood-brain-barrier or another system like the glymphatic system. *A. junii* has limited motility suggesting a somewhat similar mechanism.

*C. jiangduensis*’ association with control subjects suggests that it might be part of a healthy microbiome or at least the microbiome of elderly subjects without AD. As it is motile, perhaps it functions efficiently in the inter-cellular medium as a waste processor. While it is not a major gut bacterium, its biofilms proliferate in human wastewater treatment facilities [95,96] suggesting it may prefer the pH found there of 7-9. *P. acnes*’ ability to emit propionic and acetic acids [90] may make it difficult for *C. jiangduensis* to thrive or live. This type of mechanism provides for a long-range mechanism to reduce *C. jiangduensis* if the *P. acnes* and *C. jiangduensis* ecosystems are further apart. Further, if *C. jiangduensis* has a waste treatment function, its elimination could result in pathologies. *A. junii* also prefers a pH of 7-9, again suggesting a mechanism for anti-correlation with *P. acnes* [92].

It is not clear if the lower levels of *P. acnes* in the magenta class is due to competition with M+, an increase of M+ in niches not occupied by *P. acnes*, or other factors.

Last, given our suggestion that *P. acnes*, *A. junii*, *C. jiangduensis* and M+ may occupy distinct spatial niches, the question arises as to where these niches are. One group imaged brain tissue from ALS patients and found inter- and intra-cellular bacteria as well as fungi which is consistent with our prediction of distinct spatial niches [97]. Most important, this group observed bacterial abundance profiles in these subjects that had key similarities to the magenta microbiome. Specifically, many samples had levels of *Methylobacterium* that were several times higher than the *Propionibacterium* they found also. This, of course, suggests a multi-disease effect for bacterial infection, although the common theme reported among degenerative brain diseases is build-up of toxic protein breakdown products.

###### Points of Entry - Blood Brain Barrier and Statistics

Much has been written about the possibility of AD being a vascular disease involving the failure of the blood brain barrier (BBB) [98]. While less is known, other distribution systems like the glymphatic system could also be candidates [99].

The major reason why fluidic distribution systems could be behind our results is that there needs to be a mechanism for the random microscopic distribution of ecosystems and macroscopic distribution of class mixtures. The BBB could provide such a mechanism because it could deliver bacteria anywhere in the brain — if and as it fails. This process could be essentially a pseudo-random scattering of bacteria into the capillaries and across them when possible. On the other hand, if the BBB fails in only some localized places, we would need a spreading mechanism inside the brain. Spreading, however is more likely to cause a regional structure of ecosystems and classmixtures of them which is contradicted by our results. There are reports, however, that it happens [100]. It may be though that it is the failure of the BBB that spreads making it look like a spreading infection [101].

The hints at diversity between lobes that we have mentioned where the frontal lobe seems somewhat more associated with AD than the temporal lobe further suggest that we are observing a gradual failure of the BBB by lobe. A worsening of the failure could also account for both the rise and fall of *P. acnes* abundances. Perhaps, on the ascendant side of the *P. acnes* abundance curve, the BBB is not in as bad shape and lets through the *P.* acnes and *A. junii* ecosystems. As the pathology worsens, maybe the BBB allows more of the M+ set in and they can either outcompete the *P. acnes* or occupy a different niche outside of *P. acnes* influence where they can increase in abundance and do the damage associated with AD.

Regarding the vascular residence of observed bacteria, Emery et. al have measured the blood microbiome of AD subjects and found some of the same genera we observe in the orange class, particularly *Acidovorax* ([102] and private communication). They also prepared their samples by removing the larger vascular structures whereas we did not. This could skew the results if there is a large difference between what is in or crossed over from the capillaries and what is in larger diameter blood vessels.

If large-scale BBB failure happens in AD, bacterial introduction to the brain through a failure in the BBB could be at the root of other neurological diseases but involve failure in other parts of the brain.

On the other hand, bacteria may not be the main pathogenic factor and just be markers for the progressive failure of the BBB that is allowing other pathogenic agents in. This interpretation would allow for the patterns we have observed to only be correlative and not causal. This certainly is always worth keeping in mind, especially with the increasing evidence of the presence of fungi and viruses in the brain [23,103]. Even so, it is quite hard to conclude that all of the bacterial patterns are unrelated to the cause of AD.

## CONCLUSIONS

We detected many different species of bacteria both in the brains of AD subjects and in controls, particularly *P. acnes* which was present in 83% of samples and all but one subject. It was difficult to see major differences between the AD cases and controls because the abundance variances were large. Consequently, we undertook a different type of statistical analysis.

The first attempt, utilizing a Dirichlet-Multinomial modelling approach that utilized the disease state, found that the abundance of *P. acnes* was somewhat predictive of AD with weaker indications for other bacterial taxa. The second approach was a hybrid approach that first clustered similar samples together and then related these clusters to the disease state. Essentially, we were able to classify the samples into 5 clusters. We labeled the classes with five colors: green, orange, blue, red, and magenta. The combined data from each cluster created a class microbiome which was defined in terms of the occurrence of specific bacterial genera and species within various abundance bins. Within these microbiomes, we found that 3 principal genera: *Propionibacterium, Acinetobacter, Comamonas,* dominated 4 of 5 classes. *Comamonas* dominates a class with samples only from control subjects. The fifth class was characterized by the combined presence of *Propionibacterium* in the range of 3-30% abundance and at least one of a small set of other bacteria at levels over 30% (called M+). Various statistics suggested competitions between *P. acnes* and both *Acinetobacter spp.* and *C. jiangduensis,* with *P. acnes* outcompeting the other two taxa. In the competition between *P. acnes* and the M+ taxa, M+ appears to predominate although there are multiple explanations as described above and below.

When we looked at the class microbiome of the 2-4 samples from each subject, we were able to see that the magenta class (M+ predominant) was found in almost all the subjects with AD and occurred infrequently in the controls, suggesting that that this microbiome is pathogenic. The other classes were not strongly associated with either disease state.

We were able to compute a possible etiology for AD in terms of these microbiome classes. First, ve assumed that the microbiomes found in different samples may represent infections that began at different times. We also observed that one of these microbiomes never occurred in subjects without AD and one almost always came from AD subjects. Then, assuming that health precedes disease and by computing statistical relationships among pairs of classes, we were able to time order the appearance of the bacterial classes: green, orange, blue, red, and magenta. Green samples never came from AD subjects and magenta samples almost always came from AD subjects. This ordering also served to reveal the temporal abundance dynamics of the principal bacteria and enabled us to understand that *P. acnes* rises and falls over time but that the pathogenic magenta class does not occur where *P. acnes* peaks. Rather, it is associated with the waning of the *P. acnes* abundance, which is peculiar behavior for a single pathogen.

We obtained samples from three parts of the brain: the frontal lobe, temporal lobe, and entorhinal cortex. We were able to determine that, for the AD subjects, most of their frontal lobes contained the pathogenic class; about 1/3 contained this class in both the frontal and temporal lobes and only a couple of AD subjects contained this class only in the temporal lobe. This is consistent with medical observations that report symptoms that appear to result from damage in both lobes.We were not able to reach conclusions about the entorhinal cortex which has been suggested previously as a possible entry point for bacteria into the brain [104]. At least, for this set of subjects, this finding suggests that AD symptoms may appear first from frontal lobe damage.

The results of the LDA computations also enabled us to infer findings on the microscopic structure of the ecosystems underlying the samples and suggest how the different classes of ecosystem mixtures may be spatially distributed in the brain macroscopically. By looking at the abundance statistics of the principal microbes within a class, we concluded that they were likely part of spatially distinct ecosystems, dominated by the principal bacteria at the microscopic scale and randomly interleaved. They may be at inter- or intra-cellular locations, near or not near capillaries and so forth. Our findings on the macroscopic scale are a bit more speculative. We estimated how likely it was to sample one or more of the same classes within a subject and used these statistics to compare two possibilities. The first involved widespread homogeneous regions dominated by single classes and the second involved smaller scale randomly interleaved regions of one class (~ a few cm). These may correspond respectively to infections that enter in a limited number of locations, and then multiply and spread or, alternatively, infections that enter in a large number of locations such as when the blood brain barrier or glymphatic systems fail randomly on a microscopic scale, perhaps at the capillaries, but where the failure extends over macroscopic scales. Our results suggested the latter for 4 of the 5 classes, including the pathogenic class. So, at the microscopic scale we hypothesize interleaved ecosystems dominated by the principal bacteria and at the macroscopic scale we have mixtures of these ecosystems on sub-lobe scales, also interleaved, consistent with the class computations. So, the class temporal dynamics which we compute could be due to an evolution of the underlying ecosystems or due to changes in the entry mechanism or a combination.

These overall findings together with the *P. acnes* dynamics allow us to suggest some possibilities for the biology. Given that that the ecosystems of the principal bacteria seem to occupy spatially distinct locations on microscopic scales and that some type of competition exists between *P. acnes* and both *C. jiangduensis* and *A. junii*, these interactions need to be somewhat long range, i.e. much larger than a bacterium. Such an interaction could occur, for example, if the acidic output of *P. acnes* (propionic and acetic acids) creates an environment whose pH is too low for either the *C. jiangduensis* or *A. junii* to thrive but there could be other mechanisms, for example, involving molecular communication. The ubiquity of *P. acnes*, even as it waxes and wanes, should also prompt a reaction from the host and this could be a partial explanation of the amyloid deposits and neurofibrillary tangles, a histopathology that occurs not only with AD but other neurologicalillnesses as well - most notably general paresis associated with tertiary syphilis, a known microbially-induced neurodegenerative condition that is also characterized by amyloid deposits and neurofibrillary tangles. While we do not have direct temporal measurements in this study, it may be that *C. jiangduensis* is a critical part of a healthy brain microbiome that is involved in waste clearance. While it is not a common gut bacterium, it is found to flourish in waste-water treatment facilities in biofilm form. Its destruction by an advancing *P. acnes* infection could deprive the brain of a critical function.

What happens last in the pathogenic magenta class is the most curious. *P. acnes* falls and the M+ taxa increase. This could be because of competition where M+ drives out *P. acnes.* Alternatively, the M+ taxa may move in and proliferate in unoccupied spatial niches. These niches might not have been occupied before or could have been produced by a significant amount of damage from *P. acnes* activity allowing the M+ taxa to infiltrate into the wound. Either way, clearly there is something different going on as *P. acnes* abundances in orange are similar to magenta but the M+ did not take over then, even though some were present in low abundance. So perhaps *P. acnes* cleared the way for the M+ by removing the other principal bacteria (or hadn’t yet had time to create the M+ environment), the M+ being either late entrants or always there but not able to thrive in the earlier competitive environment. Both explanations are consistent with the sample abundance data, the former involving relative abundance changes of both and the latter involving an increase in absolute numbers of M+ but not *P. acnes.* Since we are only looking at relative abundances, it is not possible to distinguish between them. We hope that our findings prompt a reexamination of the *P. acnes* role in other infections, such as spinal cord discs, implants, and acne [51,105,106].

There is also an alternative to this bacterio-centric picture, or at least another pathogenic mechanism that might coexist with it. The microscopic and macroscopic structures that we have described suggest that there exists some sort of mechanism that introduces pathogens randomly over large areas of the brain rather than at a limited number of entry points followed by spreading of the entrant. Others have suggested that the failure of the blood brain barrier or glymphatic system could be such a model. We suggest that the failure of these systems at the capillary level with entry points, “holes”, randomly distributed on a very small scale could be consistent with our results. The evolution of the disease could be modeled by changes in porosity and location within the brain. Such a model could work with both a bacterio-centric model of pathogenicity or a model where another pathogenic microbe such as a virus or fungus or some other molecule enters the brain concurrent with the development of the magenta stage. The latter would indicate that the complex dynamics that we report are actually temporal markers for the gradual failure of a brain blood or lymph distribution system. Further research could illuminate whether it is one, the other, or both and whether the location and type of infection explains other neurological diseases.

In summary, the process described by the data is that of an evolving microbiome in the human brain that begins, perhaps as a healthy microbiome and then gradually changes until it is unquestionably associated with Alzheimer’s disease. This microbiome dynamic, however, calls out for explanation in more fundamental terms. It is a complex dynamic that likely involves the time dependence of multiple interacting systems: including the microbial ecosystems, a changing immune reaction with genetic constraints, and dynamic delivery networks driven by external factors that could have happened once or are ongoing. This work has only begun to uncover how this works. Understanding AD apparently will involve a program of discovering the workings of these fundamental components, how they affect each other and ultimately how they affect the function of the mind.

## LIST OF ABBREVIATIONS

AD: Alzheimer’s disease
AMC: Age-matched controls
CCS: Circular consensus sequence
Clr: centered log ratio
DMM: Dirichlet-multinomial model
LDA: Latent Dirichlet allocation
MLDA: Modified latent Dirichlet allocation
MCSMRT: Microbiome Classifier using Single Molecule Real-time Sequencing
OTU: Operational taxonomic unit
PacBio: Pacific Biosciences
PCA: Principal component analysis

## DECLARATIONS

### Ethics approval and consent to participate

The human biospecimens used in this study conformed to relevant ethics regulatory standards. Institutional Review Board (IRB) of Drexel University College of Medicine approved this study (IRB approval # 1410003161).

### Availability of data and material

The full-length 16S sequences have been deposited at the NIH NCBI SRA repository (BioProject PRJNA822777). A temporary reviewer’s link is: https://dataview.ncbi.nlm.nih.qov/obiect/PRJNA822777?reviewer=1d51bhv25cibkbsdaiqOdaqlfd.

Additional code, input and output files related to each analytical method are shown below.

### Additional Files

We have provided additional files relating to both analytical methods in two repositories at GitHub. Each contains a readme file with additional details.

**METHOD 1**: Files for the differential abundance analysis (Method for Individual Bacteria) using using the Dirichlet-multinomial model (DMM) are available at https://github.com/ilapides/alzheimers_method_1

**Additional file 1**: Sample information and OTU tables (multiple worksheet xlsx file, 672 KB). **Tab 1**: Sample information. **Tab 2**: Read counts after each CCS filtering step (SMRT Link v7 - MCSMRT). **Tab 3**: Read counts and taxonomic assignment with confidences for each OTU (SMRT Link v7 - MCSMRT). **Tab 4**: Read counts after each CCS filtering step (SMRT Link v9 - MCSMRT). **Tab 5**: Read counts and taxonomic assignment with confidences for each OTU (SMRT Link v9 - MCSMRT).

**Additional file 2**: Differential abundance analysis script (R Markdown, 22.1 KB).

**Additional file 3**: Brain microbiome phyloseq object (RDS file for R, 26.6 KB).

**METHOD 2**: Files for the Modified LDA analysis (Method for Multiple Bacteria) is available at https://github.com/jlapides/alzheimers_method_2. In this repository, we have included the program code for the Modified Latent Dirichlet Allocation (MLDA) analysis organized with their input files. The code is written in Wolfram Mathematica and will work for versions 12.2 and above.

**Additional_file_4**_alzheimers_input_data_and_results.xlsx (multiple worksheets, 877 KB). **Tab 1**: Readme(redo). **Tab 2**: raw count data at OTU level. **Tab 3**: raw count data at OTU level, background removed using negative controls. **Tab 4**: raw count data with background removed summed to species or genus level. **Tab 5**: raw count data with background removed at species or genus level, filtered for low counts, 4 genera removed. **Tab 6**: abundance data after background removal, etc., normalized from counts by sample, also included as Additional file 5. **Tab 7**: tk2 variable in code, each row is a sample with microbial object list, also included as Additional file 6. **Tab 8**: metadata by sample with subject, disease state, sample location, also included as Additional file 7. **Tab 9**: microbial objects after rebinning abundances to Hi and Lo. **Tab 10**: microbial objects that were renamed for MLDA. **Tab 11**: final microbial object list with occurrence counts after rebidding and renaming. **Tab 12**: sample, top classes, normalized class components, combination of 5 runs, used as input to graph, also included as Additional file 8. **Tab 13**: Approximate microbiome, list of object with class component in counts, also shown in Table S8, also shown as Additional file 9. **Tab 14**: Approximate microbiome, list of object with normalized class components, also shown as Additional file 10.

**Additional_file_5**_binning_and_object_computation_paper_v1.nb (Mathematica notebook, 35 KB).

**Additional_file_6**_biomedataFilt.csv (comma-separated values, 110 KB).

**Additional_file_7.1**_microbiome_alzheimer’s_input_11_s1.tsv (tab-separated values, 30 KB).

**Additional_file_7.2**_microbiome_alzheimer’s_input_11a_s1.tsv (tab-separated values, 32 KB).

**Additional_file_8**_classification_model_7_alzheimers_paper_v1.nb (Mathematica notebook, 1.0 MB).

**Additional_file_9**_md1UAMS.csv (tab-separated values, 6 KB). Metadata.

**Additional_file_10**_Alzheimers_AK_5_topics_a_0P3_b_0P25_cut_1_12.15rrtst_docTopicSum.csv (comma-separated values, 13 KB)).

**Additional_file_11**_graph_alzheimers_paper.nb (Mathematica code, 1.5 MB)

**Additional_file_12**_microbiome_obj_approx_cnts.csv (comma-separated values, 4 KB).

**Additional_file_13**_microbiome_obj_approx_norm.csv (comma-separated values, 7 KB).

**Additional_file_14**_allTable7.tsv (tab-separated values, 247 KB)

### Authors’ contributions

GDE and YM conceived and designed the study. AA, BS, GDE, JEK, JPE, and YM provided the materials and reagents and carried out the experiments. JRL coded and wrote the software for the method to analyze combinations of bacteria. GDE, JRL and YM analyzed and interpreted the data. GDE, JRL and YM wrote the manuscript. All authors read and approved the final manuscript.

## Acknowledgements

We are very grateful to Dr W. Sue T. Griffin (Department of Geriatrics, University of Arkansas for Medical Sciences, Little Rock, AR, USA) for kindly providing the brain tissue samples used in this study. We thank Dr Joshua Chang Mell for fruitful discussions. We also thank Dr. James Truchard for his insight and many discussions. Dr. Jeffrey Lapides would like to thank Hiroshi Inoue (former CEO of Canon U.S. Life Sciences) for many early discussions on the microbiome and Owen Dall for suggesting learning about LDA for many discussions over the years. Last, but not least, he would also like to acknowledge inspiration from Dr. David Emery and his colleagues at the University of Bristol, Dr. Shelley Allen-Birt, Dr. Maria Davies, and Dr. Nicola West. Sadly, Dr. Emery passed away as this manuscript was nearing completion.

## SUPPLEMENTARY INFORMATION

**Table S1:** OTU identified as potential contaminants OTU identifiers (column 1), species name (column 2), Genus-level confidence values (column 3), Species-level confidence values (column4), Mean of relative abundance (RAb) in negative controls (column 5), Mean of relative abundance in biological samples (column 6), Prevalence-based score statistic P (column 7). OTU are identified as contaminant (TRUE) when they show a score statistic P > 0.5 or when RAb in negative controls is higher than RAb in biological samples (column 8).

| OTU_ID | species | genus<br>conf | species<br>conf | RAb in<br>negative<br>controls | RAb in<br>samples | decontam<br>statistic<br>score $P$ | contaminant |
| --- | --- | --- | --- | --- | --- | --- | --- |
| OTU_771 | *Achromobacter denitrificans* | 0.4089 | 0.0341 | 0.0373 | 0 | 0.9996 | TRUE |
| OTU_2 | *Achromobacter xylosoxidans* | 0.8292 | 0.494 | 16.6884 | 1.8438 | 0.9483 | TRUE |
| OTU_555 | *Achromobacter xylosoxidans* | 0.4696 | 0.0441 | 0.0117 | 0 | 0.9996 | TRUE |
| OTU_566 | *Achromobacter xylosoxidans* | 0.3483 | 0.0241 | 0.0024 | 0 | NA | TRUE |
| OTU_650 | *Achromobacter xylosoxidans* | 0.5302 | 0.0441 | 0.0222 | 0.0002 | 0.9688 | TRUE |
| OTU_9 | *Acinetobacter junii* | 0.973 | 0.9346 | 0.0007 | 11.9637 | NA | FALSE |
| OTU_3 | *Anabaena cylindrica* | 0.0019 | 0 | 14.1167 | 0.9935 | 0.8908 | TRUE |
| OTU_549 | *Anabaena cylindrica* | 0.0017 | 0 | 0.0016 | 0 | NA | TRUE |
| OTU_295 | *Bordetella petrii* | 0.8292 | 0.7194 | 0.0016 | 0.0347 | 0.9688 | TRUE |
| OTU_916 | *Corynebacterium mucifaciens* | 0.9531 | 0.0541 | 0.0036 | 0 | NA | TRUE |
| OTU_103 | *Corynebacterium ureicelerivorans* | 0.9862 | 0.3055 | 0.3105 | 0.121 | 0.7531 | TRUE |
| OTU_8 | *Delftia acidovorans* | 0.9796 | 0.5993 | 0.0012 | 1.8643 | 0.6502 | TRUE |
| OTU_467 | *Hydrotalea flava* | 0.8763 | 0.3055 | 0.0089 | 0.0012 | 0.9925 | TRUE |
| OTU_6 | *Hydrotalea flava* | 0.9796 | 0.8396 | 23.9855 | 15.1389 | NA | TRUE |
| OTU_729 | *Hydrotalea flava* | 0.2877 | 0.0141 | 0.028 | 0.0322 | 0.8291 | TRUE |
| OTU_1060 | *Hydrotalea sandarakina* | 0.0497 | 0.0026 | 0.0086 | 0.022 | 0.8291 | TRUE |
| OTU_1099 | *Pelomonas puraquae* | 0.4696 | 0.0441 | 0.0022 | 0 | NA | TRUE |
| OTU_1054 | *Pelomonas saccharophila* | 0.3483 | 0.0541 | 0.0007 | 0 | NA | TRUE |
| OTU_7 | *Pelomonas saccharophila* | 0.7349 | 0.3997 | 3.5897 | 0.6594 | 0.8137 | TRUE |
| OTU_262 | *Peredibacter starrii* | 0.0014 | 0 | 0.0198 | 0.0013 | 0.9964 | TRUE |
| OTU_5 | *Peredibacter starrii* | 0.0014 | 0 | 21.5241 | 3.8676 | 0.6401 | TRUE |
| OTU_1 | *Propionibacterium acnes* | 0.973 | 0.9575 | 0.0005 | 20.6146 | NA | FALSE |
| OTU_835 | *Pseudomonas hibiscicola* | 0.3483 | 0.0341 | 0.1144 | 0 | 0.9996 | TRUE |
| OTU_65 | *Sediminibacterium ginsengisoli* | 0.0497 | 0.0043 | 0.5176 | 0.4599 | 0.7381 | TRUE |
| OTU_1082 | *Sphingomonas sanxanigenens* | 0.0005 | 0 | 0.001 | 0 | NA | TRUE |
| OTU_36 | *Sphingomonas sanxanigenens* | 0.0006 | 0 | 0.6718 | 0.0138 | 0.9688 | TRUE |
| OTU_436 | *Sphingomonas sanxanigenens* | 0.0008 | 0 | 0.0063 | 0.0002 | 0.9688 | TRUE |
| OTU_1250 | *Stenotrophomonas humi* | 0.0019 | 0 | 0.0005 | 0 | NA | TRUE |
| OTU_4 | *Stenotrophomonas maltophilia* | 0.6878 | 0.3055 | 18.3224 | 0.3844 | 0.9426 | TRUE |

**Table S2:**
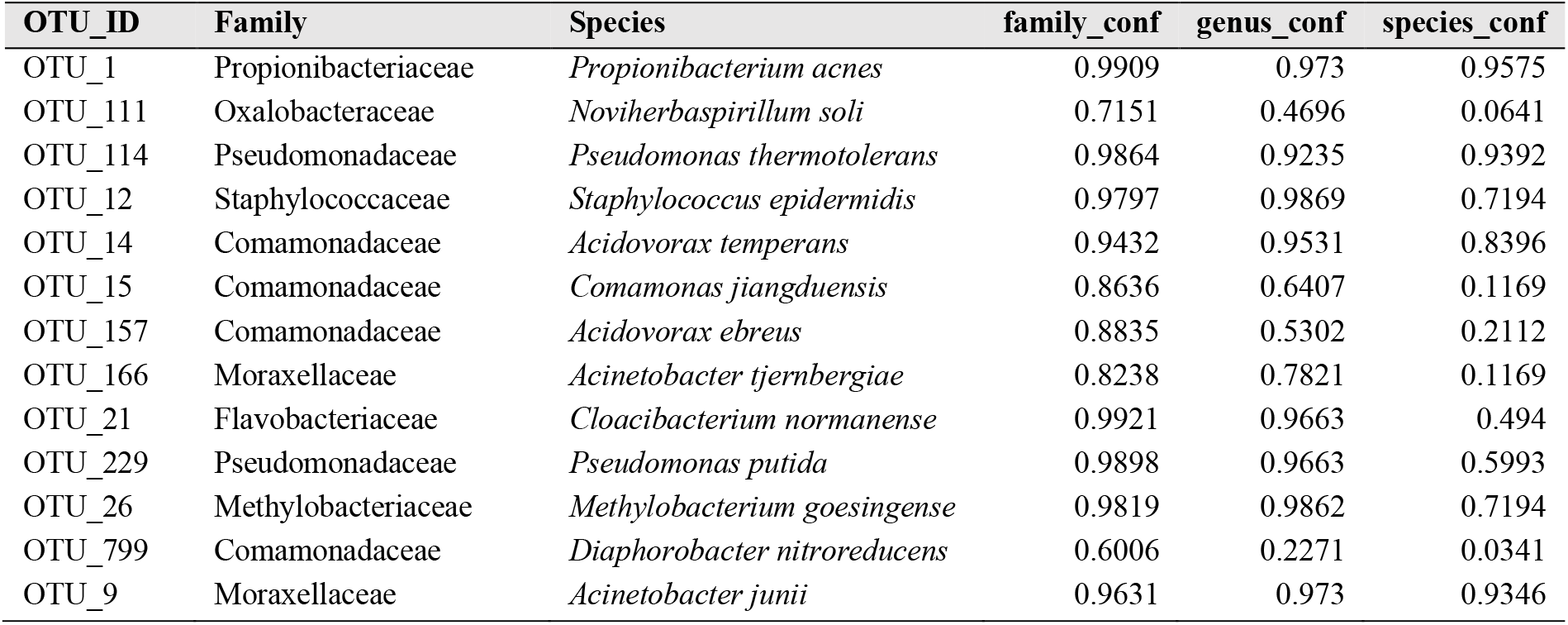
Taxonomic classification of OTU that shift in abundance between the Alzheimer’s disease group and the age-matched control group. For each OTU of interest, the taxonomic assignment and the family-, genus- and species-confidence values are reported.

**Table S3:**
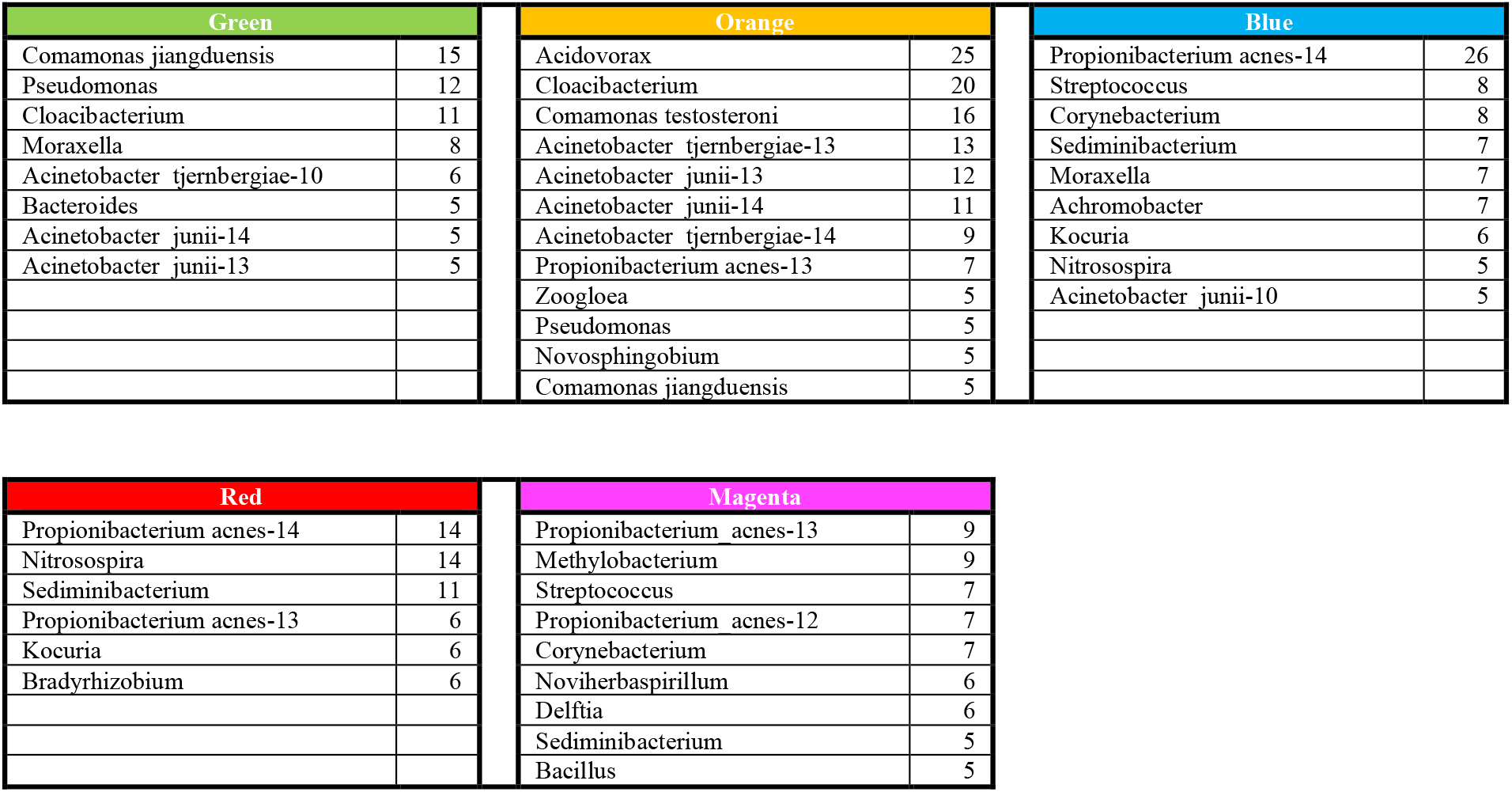
Classification results with lower occurring objects summed over abundance bin.

**Table S4:**
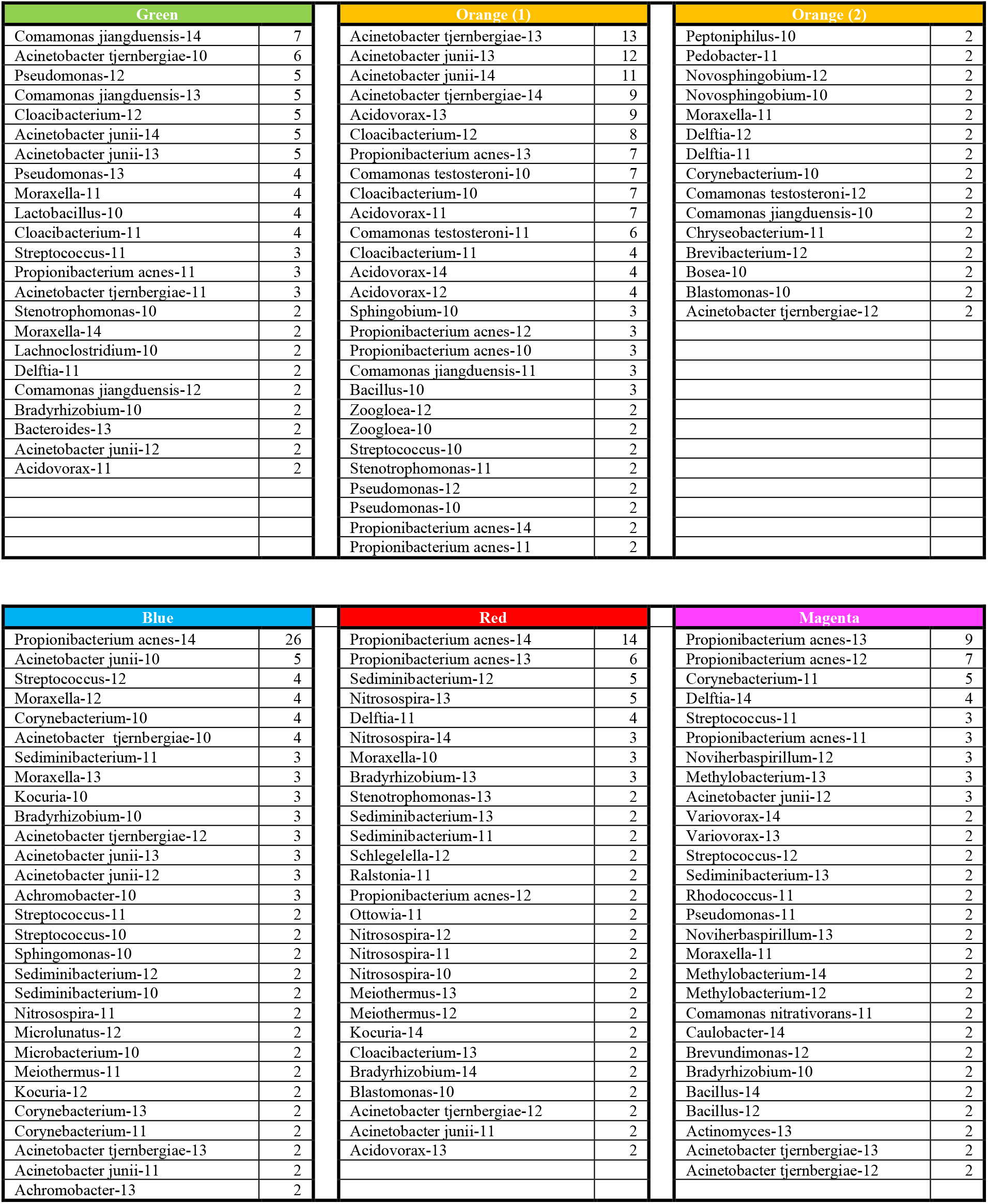
Classification results without summing lower occurring objects over abundance bin.

**Table S5:** Classification results without summing lower occurring objects over abundance bin after object merging.

| Green |  | Orange |  | Blue |  |
| --- | --- | --- | --- | --- | --- |
| Comamonas jiangduensis-hi | 14 | Acidovorax-hi | 17 | Propionibacterium acnes-14 | 26 |
| Pseudomonas-hi | 10 | Acinetobacter tjernbergiae-13 | 13 | Moraxella-hi | 7 |
| Cloacibacterium-hi | 7 | LoCnt-hi | 12 | Streptococcus-hi | 4 |
| Acinetobacter junii-14 | 5 | Acinetobacter junii-13 | 12 | LoCnt-hi | 3 |
| Acinetobacter junii-13 | 5 | Acinetobacter junii-14 | 11 | Kocuria-hi | 3 |
| Moraxella-hi | 3 | Cloacibacterium-hi | 9 | Acinetobacter tjernbergiae-12 | 3 |
| Acinetobacter junii-12 | 2 | Acinetobacter tjernbergiae-14 | 9 | Acinetobacter junii-13 | 3 |
|  |  | Propionibacterium acnes-13 | 7 | Acinetobacter junii-12 | 3 |
|  |  | Propionibacterium acnes-12 | 3 | Achromobacter-hi | 3 |
|  |  | Sediminibacterium-hi | 2 | Sediminibacterium-hi | 2 |
|  |  | Pseudomonas-hi | 2 | Nitrosospira-hi | 2 |
|  |  | Propionibacterium acnes-14 | 2 | Acinetobacter tjernbergiae-13 | 2 |
|  |  | Delftia-hi | 2 |  |  |
|  |  | Acinetobacter tjernbergiae-12 | 2 |  |  |

| Red |  | Magenta |  |
| --- | --- | --- | --- |
| Propionibacterium acnes-14 | 14 | Propionibacterium acnes-13 | 9 |
| Nitrosospira-hi | 10 | LoCnt-hi | 9 |
| Sediminibacterium-hi | 8 | Propionibacterium acnes-12 | 7 |
| Propionibacterium acnes-13 | 6 | Methylobacterium-hi | 7 |
| Bradyrhizobium-hi | 6 | LoCnt-14 | 6 |
| LoCnt-hi | 5 | Delftia-hi | 6 |
| Meiothermus-hi | 4 | Noviherbaspirillum-hi | 5 |
| Kocuria-hi | 4 | Streptococcus-hi | 4 |
| Stenotrophomonas-hi | 3 | Bacillus-hi | 4 |
| Cloacibacterium-hi | 3 | Sediminibacterium-hi | 3 |
| Propionibacterium acnes-12 | 2 | Acinetobacter junii-12 | 3 |
| Acinetobacter tjernbergiae-12 | 2 | Acinetobacter tjernbergiae-13 | 2 |
| Acidovorax-hi | 2 | Acinetobacter tjernbergiae-12 | 2 |
|  |  | Acidovorax-hi | 2 |
|  |  | Achromobacter-hi | 2 |

**Table S6:** Main Microbial Objects By Class and Disease State - Individual Samples

| Green |  |  |  | Orange |  |  |  |
| --- | --- | --- | --- | --- | --- | --- | --- |
| Propionibacterium | Acinetobacter | Comamonas | Disease | Propionibacterium | Acinetobacter | Comamonas | Disease |
| P. acnes-13 | A. junii-13 | none | AD | none | none | C. jiangduensis-14 | C |
| P. acnes-14 | none | none | C | none | A. junii-13 | C. jiangduensis-12 | C |
| P. acnes-14 | A. junii-13 | none | AD | none | A. junii-12 | C. jiangduensis-14 | C |
| P. acnes-13 | A. junii-13 | none | AD | none | A. junii-14 | C. jiangduensis-13 | C |
| P. acnes-11 | A. junii-13 | none | AD | none | A. junii-13 | C. jiangduensis-14 | C |
| P. acnes-9 | A. junii-13 | none | AD | P. acnes-10 | A. junii-13 | C. jiangduensis-14 | C |
| P. acnes-13 | A. junii-14 | none | C | P. acnes-14 | A. junii-14 | C. jiangduensis-13 | C |
| P. acnes-10 | A. junii-13 | C. jiangduensis-7 | AD | P. acnes-11 | A. junii-13 | C. jiangduensis-14 | C |
| P. acnes-9 | A. junii-14 | C. jiangduensis-11 | C | P. acnes-7 | A. junii-14 | C. jiangduensis-13 | C |
| P. acnes-7 | A. junii-14 | C. jiangduensis-10 | C | P. acnes-8 | A. junii-14 | C. jiangduensis-13 | C |
| P. acnes-12 | A. junii-13 | none | AD | P. acnes-11 | A. junii-12 | none | C |
| P. acnes-13 | A. junii-13 | none | AD | none | A. junii-10 | C. jiangduensis-14 | C |
| P. acnes-10 | A. junii-13 | C. jiangduensis-9 | AD | P. acnes-11 | A. junii-13 | C. jiangduensis-14 | C |
| P. acnes-8 | A. junii-13 | none | AD | P. acnes-13 | A. junii-14 | C. jiangduensis-11 | C |
| P. acnes-10 | A. junii-14 | C. jiangduensis-11 | C | none | none | C. jiangduensis-13 | C |
| none | A. junii-14 | C. jiangduensis-11 | C | none | none | C. jiangduensis-12 | C |
| none | A. junii-14 | none | AD |  |  |  |  |
| P. acnes-12 | A. junii-13 | none | AD |  |  |  |  |
| none | A. junii-13 | none | AD |  |  |  |  |
| P. acnes-13 | A. junii-14 | none | AD |  |  |  |  |
| P. acnes-13 | A. junii-14 | none | C |  |  |  |  |
| P. acnes-11 | A. junii-14 | none | AD |  |  |  |  |
| P. acnes-12 | none | C. jiangduensis-7 | C |  |  |  |  |
| P. acnes-9 | A. junii-14 | C. jiangduensis-10 | C |  |  |  |  |
| P. acnes-7 | A. junii-14 | none | C |  |  |  |  |
| P. acnes-13 | none | none | C |  |  |  |  |
| none | A. junii-11 | none | AD |  |  |  |  |

| Blue |  |  |  | Red |  |  |  |
| --- | --- | --- | --- | --- | --- | --- | --- |
| Propionibacterium | Acinetobacter | Comamonas | Disease | Propionibacterium | Acinetobacter | Comamonas | Disease |
| P. acnes-14 | none | none | C | P. acnes-14 | none | none | AD |
| P. acnes-14 | A. junii-11 | none | AD | P. acnes-14 | none | none | AD |
| P. acnes-14 | none | none | AD | P. acnes-14 | none | none | C |
| P. acnes-14 | none | none | C | P. acnes-13 | none | none | AD |
| P. acnes-14 | none | none | C | P. acnes-11 | none | none | AD |
| P. acnes-14 | A. junii-10 | none | C | P. acnes-12 | none | none | C |
| P. acnes-14 | A. junii-9 | none | C | P. acnes-14 | A. junii-13 | none | AD |
| P. acnes-14 | A. junii-9 | none | C | P. acnes-13 | none | none | AD |
| P. acnes-14 | A. junii-12 | none | C | none | none | none | C |
| P. acnes-14 | A. junii-10 | none | C | P. acnes-13 | none | none | AD |
| P. acnes-14 | A. junii-13 | none | C | P. acnes-14 | none | none | C |
| P. acnes-14 | none | none | AD | P. acnes-12 | none | none | AD |
| P. acnes-14 | A. junii-10 | none | C | none | none | none | C |
| P. acnes-14 | none | none | AD | P. acnes-14 | none | none | C |
| P. acnes-14 | A. junii-13 | none | C | P. acnes-14 | none | none | AD |
| P. acnes-14 | A. junii-12 | none | AD | P. acnes-14 | A. junii-11 | none | C |
| P. acnes-14 | A. junii-13 | none | C | P. acnes-14 | none | none | AD |
| P. acnes-14 | none | none | AD | P. acnes-13 | A. junii-11 | none | AD |
| none | none | none | AD | P. acnes-14 | none | none | AD |
| P. acnes-14 | A. junii-10 | none | AD | P. acnes-14 | none | none | AD |
| P. acnes-14 | none | none | C | P. acnes-14 | none | none | AD |
| none | A. junii-10 | none | C | P. acnes-14 | A. junii-10 | none | AD |
| P. acnes-14 | A. junii-11 | none | AD | P. acnes-13 | none | none | C |
| P. acnes-14 | A. junii-12 | none | C | none | none | none | C |
| P. acnes-12 | A. junii-9 | none | C | P. acnes-13 | none | none | AD |
| P. acnes-14 | none | none | C | P. acnes-14 | none | none | AD |
| P. acnes-14 | none | none | C |  |  |  |  |
| P. acnes-14 | none | none | C |  |  |  |  |
| P. acnes-14 | none | none | AD |  |  |  |  |

| Magenta |  |  |  |
| --- | --- | --- | --- |
| Propionibacterium | Acinetobacter | Comamonas | Disease |
| P. acnes-13 | A. junii-12 | none | AD |
| none | none | none | AD |
| none | none | none | AD |
| P. acnes-13 | none | none | C |
| P. acnes-12 | none | none | AD |
| P. acnes-12 | none | none | AD |
| P. acnes-12 | A. junii-14 | none | AD |
| P. acnes-12 | A. junii-13 | none | AD |
| P. acnes-13 | none | none | AD |
| P. acnes-13 | none | none | AD |
| P. acnes-13 | none | none | AD |
| none | none | none | AD |
| P. acnes-12 | none | none | AD |
| P. acnes-11 | none | none | AD |
| P. acnes-13 | none | none | AD |
| P. acnes-10 | A. junii-12 | none | C |
| P. acnes-13 | none | none | AD |
| P. acnes-13 | none | none | AD |
| P. acnes-13 | none | none | AD |
| P. acnes-11 | none | none | AD |
| none | none | none | AD |
| P. acnes-12 | none | none | C |
| P. acnes-12 | A. junii-12 | none | AD |
| P. acnes-11 | none | none | AD |

**Figure S7:**
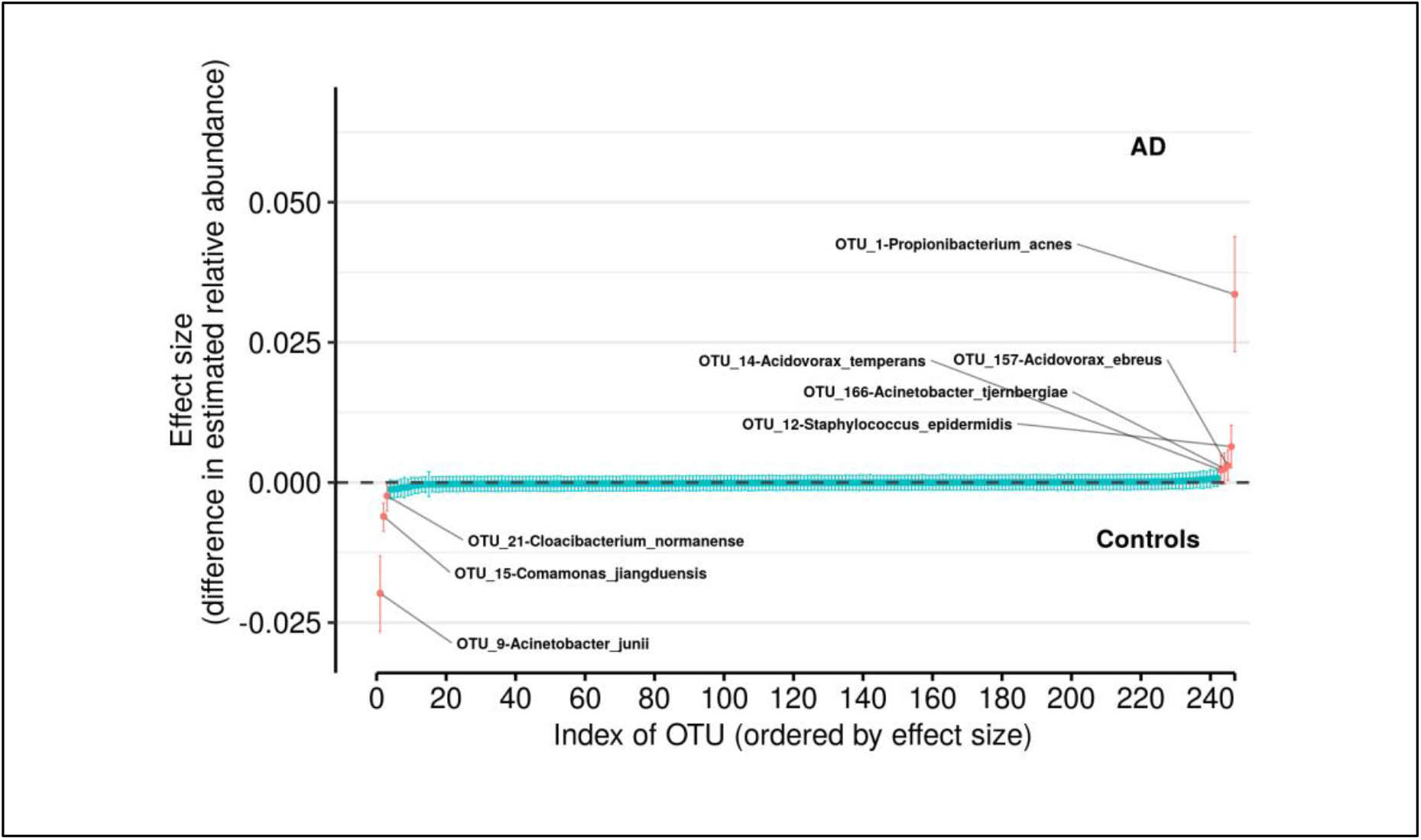
Differences in relative abundance between the Alzheimer’s disease (AD) group and the age-matched control group (controls). The relative abundances were estimated for each OTU from each group through hierarchical Bayesian modeling while ignoring the non-independence of the samples. The vertical axis shows the difference for the estimated relative abundance of OTU between the AD and control groups. Points are the means of PPD and the whiskers show the 95% equal tail probability intervals of PPD (see Materials and methods).:

**Table S8:**
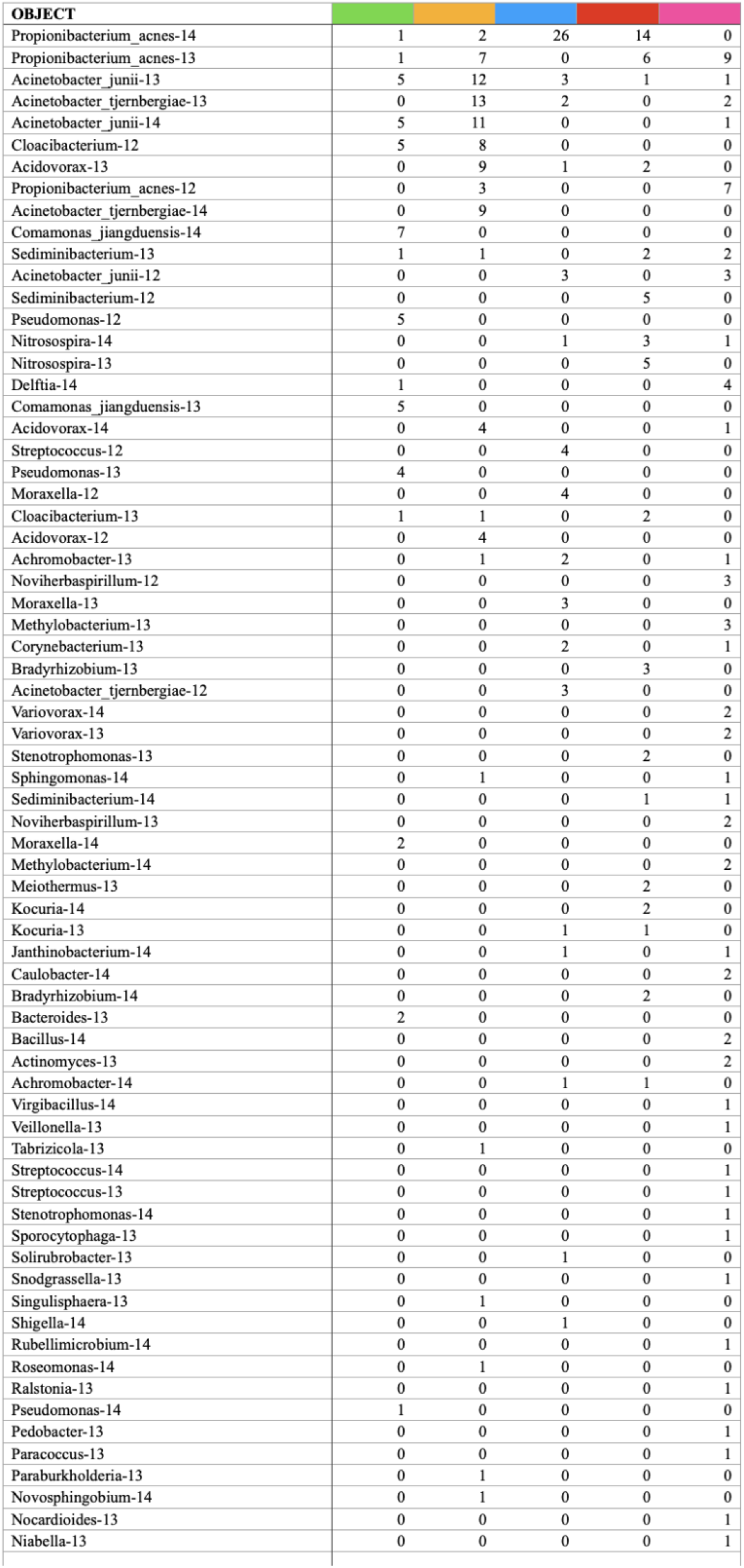
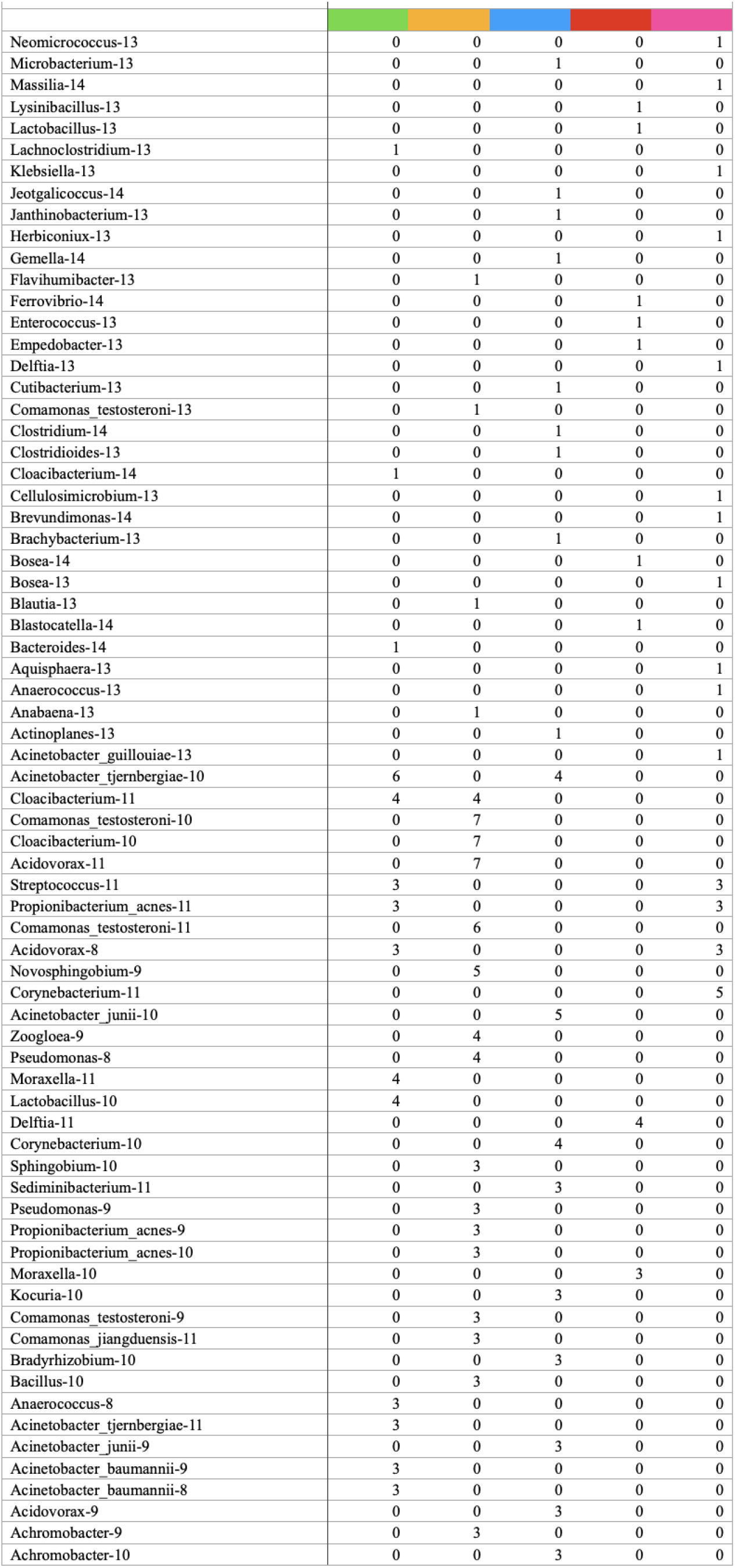
microbiome objects approximated from sample input data of a given color - counts.

## REFERENCES

1. Fischer O. Miliaere Nekrosen mit drusigen Wucherungen der Neurofibrillen, eine regelmassige Veraenderung der Hirnrinde bei seniler Demenz. Monatsschr Psychiat Neurol. 1907;22: 361–372.

2. Fischer O. Die presbyophrene Demenz, deren anatomische Grundlage und klinische Abgrenzung. Zeitschrift für die gesamte Neurol und Psychiatr. 1910;3: 371–471.

3. Alzheimer A. Über eine eigenartige Erkrankung der Hirnrinde. Allg Z Psych Psych-gerich Med. 1907;64: 146–148.

4. Goedert M. Oskar Fischer and the study of dementia. Brain. 2009;132: 1102–1111. doi:10.1093/brain/awn256

5. 2020 Alzheimer’s disease facts and figures. Alzheimer’s Dement. 2020;16: 391–460. doi:10.1002/alz.12068

6. Mendez MF. Early-Onset Alzheimer Disease. Neurol Clin. 2017;35: 263–281. doi:10.1016/j.ncl.2017.01.005

7. Selkoe DJ, Hardy J. The amyloid hypothesis of Alzheimer’s disease at 25 years. EMBO Mol Med. 2016;8: 595–608. doi:10.15252/emmm.201606210

8. Hardy J, Allsop D. Amyloid deposition as the central event in the aetiology of Alzheimer’s disease. Trends Pharmacol Sci. 1991;12: 383–388. doi:10.1016/0165-6147(91)90609-V

9. Hardy J, Selkoe DJ. The amyloid hypothesis of Alzheimer’s disease: Progress and problems on the road to therapeutics. Science (80-). 2002;297: 353–356. doi:10.1126/science.1072994

10. Bolduc DM, Montagna DR, Seghers MC, Wolfe MS, Selkoe DJ. The amyloid-beta forming tripeptide cleavage mechanism of γ-secretase. Elife. 2016;5. doi:10.7554/eLife.17578

11. Gu L, Guo Z. Alzheimer’s Aβ42 and Aβ40 peptides form interlaced amyloid fibrils. J Neurochem. 2013;126: 305–311. doi:10.1111/jnc.12202

12. Terrill-Usery SE, Colvin BA, Davenport RE, Nichols MR. Aβ40 has a subtle effect on Aβ42 protofibril formation, but to a lesser degree than Aβ42 concentration, in Aβ42/Aβ40 mixtures. Arch Biochem Biophys. 2016;597: 1–11. doi:10.1016/j.abb.2016.03.017

13. Dunys J, Valverde A, Checler F. Are N- And C-terminally truncated A β species key pathological triggers in Alzheimer’s disease? J Biol Chem. 2018;293: 15419–15428. doi:10.1074/jbc.R118.003999

14. Li C, Götz J. Somatodendritic accumulation of Tau in Alzheimer’s disease is promoted by Fyn-mediated local protein translation. EMBO J. 2017;36: 3120–3138. doi:10.15252/embj.201797724

15. Larson M, Sherman MA, Amar F, Nuvolone M, Schneider JA, Bennett DA, et al. The complex PrPc-Fyn couples human oligomeric Aβ with pathological tau changes in Alzheimer’s disease. J Neurosci. 2012;32: 16857–16871. doi:10.1523/JNEUROSCI.1858-12.2012

16. Nisbet RM, Götz J. Amyloid-β and Tau in Alzheimer’s Disease: Novel Pathomechanisms and Non-Pharmacological Treatment Strategies. Perry G, Avila J, Moreira PI, Sorensen AA, Tabaton M, editors. J Alzheimer’s Dis. 2018;64: S517–S527. doi:10.3233/JAD-179907

17. Vergara C, Houben S, Suain V, Yilmaz Z, De Decker R, Vanden Dries V, et al. Amyloid-β pathology enhances pathological fibrillary tau seeding induced by Alzheimer PHF in vivo. Acta Neuropathol. 2019;137: 397–412. doi:10.1007/s00401-018-1953-5

18. Higashi T, Nishii R, Kagawa S, Kishibe Y, Takahashi M, Okina T, et al. 18F-FPYBF-2, a new F-18-labelled amyloid imaging PET tracer: first experience in 61 volunteers and 55 patients with dementia. Ann Nucl Med. 2018;32: 206–216. doi:10.1007/s12149-018-1236-1

19. Cohen AD, Rabinovici GD, Mathis CA, Jagust WJ, Klunk WE, Ikonomovic MD. Using Pittsburgh Compound B for In Vivo PET Imaging of Fibrillar Amyloid-Beta. Advances in Pharmacology. Academic Press Inc.; 2012. pp. 27–81. doi:10.1016/B978-0-12-394816-8.00002-7

20. Osorio C, Kanukuntla T, Diaz E, Jafri N, Cummings M, Sfera A. The post-amyloid era in Alzheimer’s disease: Trust your gut feeling. Front Aging Neurosci. 2019;11. doi:10.3389/fnagi.2019.00143

21. Sochocka M, Zwolińska K, Leszek J. The Infectious Etiology of Alzheimer’s Disease. Curr Neuropharmacol. 2017;15. doi:10.2174/1570159x15666170313122937

22. Itzhaki RF, Lathe R, Balin BJ, Ball MJ, Bearer EL, Braak H, et al. Microbes and Alzheimer’s disease. J Alzheimer’s Dis. 2016;51: 979–984. doi:10.3233/JAD-160152

23. Itzhaki RF, Golde TE, Heneka MT, Readhead B. Do infections have a role in the pathogenesis of Alzheimer disease? Nat Rev Neurol. 2020;16: 193–197. doi:10.1038/s41582-020-0323-9

24. Fülöp T, Munawara U, Larbi A, Desroches M, Rodrigues S, Catanzaro M, et al. Targeting Infectious Agents as a Therapeutic Strategy in Alzheimer’s Disease. CNS Drugs. 2020;34: 673–695. doi:10.1007/s40263-020-00737-1

25. Moir RD, Lathe R, Tanzi RE. The antimicrobial protection hypothesis of Alzheimer’s disease. Alzheimer’s Dement. 2018;14: 1602–1614. doi:10.1016/j.jalz.2018.06.3040

26. Itzhaki RF. Corroboration of a major role for herpes simplex virus type 1 in Alzheimer’s disease. Front Aging Neurosci. 2018;10. doi:10.3389/fnagi.2018.00324

27. Wozniak M, Mee A, Itzhaki R. Herpes simplex virus type 1 DNA is located within Alzheimer’s disease amyloid plaques. J Pathol. 2009;217: 131–138. doi:10.1002/path.2449

28. Tzeng NS, Chung CH, Lin FH, Chiang CP, Yeh C Bin, Huang SY, et al. Anti-herpetic Medications and Reduced Risk of Dementia in Patients with Herpes Simplex Virus Infections—a Nationwide, Population-Based Cohort Study in Taiwan. Neurotherapeutics. 2018;15: 417–429. doi:10.1007/s13311-018-0611-x

29. Eimer WA, Vijaya Kumar DK, Navalpur Shanmugam NK, Rodriguez AS, Mitchell T, Washicosky KJ, et al. Alzheimer’s Disease-Associated β-Amyloid Is Rapidly Seeded by Herpesviridae to Protect against Brain Infection. Neuron. 2018;99: 56–63.e3. doi:10.1016/j.neuron.2018.06.030

30. Ezzat K, Pernemalm M, Pålsson S, Roberts TC, Järver P, Dondalska A, et al. The viral protein corona directs viral pathogenesis and amyloid aggregation. Nat Commun. 2019;10: 2331. doi:10.1038/s41467-019-10192-2

31. Readhead B, Haure-Mirande J-VV, Funk CC, Richards MA, Shannon P, Haroutunian V, et al. Multiscale Analysis of Independent Alzheimer’s Cohorts Finds Disruption of Molecular, Genetic, and Clinical Networks by Human Herpesvirus. Neuron. 2018;99: 64–82.e7. doi:10.1016/j.neuron.2018.05.023

32. Jeong HH, Liu Z. Are HHV-6A and HHV-7 Really More Abundant in Alzheimer’s Disease? Neuron. 2019;104: 1034–1035. doi:10.1016/j.neuron.2019.11.009

33. Readhead B, Haure-Mirande JV, Ehrlich ME, Gandy S, Dudley JT. Clarifying the Potential Role of Microbes in Alzheimer’s Disease. Neuron. 2019;104: 1036–1037. doi:10.1016/j.neuron.2019.11.008

34. Rizzo R. Controversial role of herpesviruses in Alzheimer’s disease. Spindler KR, editor. PLoS Pathog. 2020;16: e1008575. doi:10.1371/journal.ppat.1008575

35. Chorlton SD. Reanalysis of Alzheimer’s brain sequencing data reveals absence of purported HHV6A and HHV7. J Bioinform Comput Biol. 2020;18: 2050012. doi:10.1142/S0219720020500122

36. Allnutt MA, Johnson K, Bennett DA, Connor SM, Troncoso JC, Pletnikova O, et al. Human Herpesvirus 6 Detection in Alzheimer’s Disease Cases and Controls across Multiple Cohorts. Neuron. 2020; 105: 1027–1035.e2. doi:10.1016/j.neuron.2019.12.031

37. Miklossy J. Emerging roles of pathogens in Alzheimer disease. Expert Rev Mol Med. 2011;13: e30. doi:10.1017/s1462399411002006

38. Miklossy J. Bacterial Amyloid and DNA are Important Constituents of Senile Plaques: Further Evidence of the Spirochetal and Biofilm Nature of Senile Plaques. J Alzheimer’s Dis. 2016;53: 1459–1473. doi:10.3233/JAD-160451

39. Carter CJ. Genetic, Transcriptome, Proteomic, and Epidemiological Evidence for Blood-Brain Barrier Disruption and Polymicrobial Brain Invasion as Determinant Factors in Alzheimer’s Disease. J Alzheimer’s Dis Reports. 2017;1: 125–157. doi:10.3233/adr-170017

40. Woods JJ, Skelding KA, Martin KL, Aryal R, Sontag E, Johnstone DM, et al. Assessment of evidence for or against contributions of Chlamydia pneumoniae infections to Alzheimer’s disease etiology. Brain Behav Immun. 2020;83: 22–32. doi:10.1016/j.bbi.2019.10.014

41. Al-Atrache Z, Lopez DB, Hingley ST, Appelt DM. Astrocytes infected with Chlamydia pneumoniae demonstrate altered expression and activity of secretases involved in the generation of ?-amyloid found in Alzheimer disease. BMC Neurosci. 2019;20: 6. doi:10.1186/s12868-019-0489-5

42. Little CS, Hammond CJ, MacIntyre A, Balin BJ, Appelt DM. Chlamydia pneumoniae induces Alzheimer-like amyloid plaques in brains of BALB/c mice. Neurobiol Aging. 2004;25: 419–429. doi:10.1016/S0197-4580(03)00127-1

43. Chen CK, Wu YT, Chang YC. Association between chronic periodontitis and the risk of Alzheimer’s disease: A retrospective, population-based, matched-cohort study. Alzheimer’s Res Ther. 2017;9: 56. doi:10.1186/s13195-017-0282-6

44. Dominy SS, Lynch C, Ermini F, Benedyk M, Marczyk A, Konradi A, et al. Porphyromonas gingivalis in Alzheimer’s disease brains: Evidence for disease causation and treatment with small-molecule inhibitors. Sci Adv. 2019;5: eaau3333. doi:10.1126/sciadv.aau3333

45. Haditsch U, Roth T, Rodriguez L, Hancock S, Cecere T, Nguyen M, et al. Alzheimer’s Disease-Like Neurodegeneration in Porphyromonas gingivalis Infected Neurons with Persistent Expression of Active Gingipains. J Alzheimer’s Dis. 2020;75: 1301–1317. doi:10.3233/JAD-200393

46. Balin BJ, Gérard HC, Arking EJ, Appelt DM, Branigan PJ, Abrams JT, et al. Identification and localization of Chlamydia pneumoniae in the Alzheimer’s brain. Med Microbiol Immunol. 1998;187: 23–42. doi:10.1007/s004300050071

47. Post JC, Preston RA, Aul JJ, Larkins-Pettigrew M, Rydquist-White J, Anderson KW, et al. Molecular analysis of bacterial pathogens in otitis media with effusion. JAMA. 273: 1598–604. Available: http://www.ncbi.nlm.nih.gov/pubmed/7745773

48. Costerton W, Veeh R, Shirtliff M, Pasmore M, Post C, Ehrlich G. The application of biofilm science to the study and control of chronic bacterial infections. J Clin Invest. 2003;112: 1466–77. doi:10.1172/JCI20365

49. Ehrlich GD, Hu FZ, Shen K, Stoodley P, Post JC. Bacterial plurality as a general mechanism driving persistence in chronic infections. Clin Orthop Relat Res. 2005; 20-4. doi:10.1097/00003086-200508000-00005

50. Ehrlich GD, Ahmed A, Earl J, Hiller NL, Costerton JW, Stoodley P, et al. The distributed genome hypothesis as a rubric for understanding evolution in situ during chronic bacterial biofilm infectious processes. FEMS Immunol Med Microbiol. 2010;59: 269–79. doi:10.1111/j.1574-695X.2010.00704.x

51. Ehrlich GD, DeMeo PJ, Costerton JW, Winkler H, editors. Culture Negative Orthopedic Biofilm Infections. Berlin, Heidelberg: Springer Berlin Heidelberg; 2012. doi:10.1007/978-3-642-29554-6

52. Stoodley P, Ehrlich GD, Sedghizadeh PP, Hall-Stoodley L, Baratz ME, Altman DT, et al. Orthopaedic biofilm infections. Curr Orthop Pract. 2011;22: 558–563. doi:10.1097/BCO.0b013e318230efcf

53. Post JC, Aul JJ, White GJ, Wadowsky RM, Zavoral T, Tabari R, et al. PCR-based detection of bacterial DNA after antimicrobial treatment is indicative of persistent, viable bacteria in the chinchilla model of otitis media. Am J Otolaryngol. 17: 106–11. doi:10.1016/s0196-0709(96)90005-8

54. Aul JJ, Anderson KW, Wadowsky RM, Doyle WJ, Kingsley LA, Post JC, et al. Comparative evaluation of culture and PCR for the detection and determination of persistence of bacterial strains and DNAs in the Chinchilla laniger model of otitis media. Ann Otol Rhinol Laryngol. 1998;107: 508–13. doi:10.1177/000348949810700609

55. Dingman JR, Rayner MG, Mishra S, Zhang Y, Ehrlich MD, Post JC, et al. Correlation between presence of viable bacteria and presence of endotoxin in middle-ear effusions. J Clin Microbiol. 1998;36: 3417–9. doi:10.1128/JCM.36.11.3417-3419.1998

56. Tuttle MS, Mostow E, Mukherjee P, Hu FZ, Melton-Kreft R, Ehrlich GD, et al. Characterization of bacterial communities in venous insufficiency wounds by use of conventional culture and molecular diagnostic methods. J Clin Microbiol. 2011;49: 3812–9. doi:10.1128/JCM.00847-11

57. Nickel JC, Stephens A, Landis JR, Chen J, Mullins C, van Bokhoven A, et al. Search for Microorganisms in Men with Urologic Chronic Pelvic Pain Syndrome: A Culture-Independent Analysis in the MAPP Research Network. J Urol. 2015;194: 127–35. doi:10.1016/j.juro.2015.01.037

58. Nickel JC, Stephens A, Landis JR, Mullins C, van Bokhoven A, Lucia MS, et al. Assessment of the Lower Urinary Tract Microbiota during Symptom Flare in Women with Urologic Chronic Pelvic Pain Syndrome: A MAPP Network Study. J Urol. 2016;195: 356–62. doi:10.1016/j.juro.2015.09.075

59. Earl JP, Adappa ND, Krol J, Bhat AS, Balashov S, Ehrlich RL, et al. Species-level bacterial community profiling of the healthy sinonasal microbiome using Pacific Biosciences sequencing of full-length 16S rRNA genes 06 Biological Sciences 0604 Genetics 06 Biological Sciences 0605 Microbiology. Microbiome. 2018;6: 190. doi:10.1186/s40168-018-0569-2

60. Socarras KM, Earl JP, Krol JE, Bhat A, Pabilonia M, Harrison MH, et al. Species-Level Profiling of Ixodes pacificus Bacterial Microbiomes Reveals High Variability Across Short Spatial Scales at Different Taxonomic Resolutions. Genet Test Mol Biomarkers. 2021;25: 551–562. doi:10.1089/gtmb.2021.0088

61. Emery DC, Shoemark DK, Batstone TE, Waterfall CM, Coghill JA, Cerajewska TL, et al. 16S rRNA next generation sequencing analysis shows bacteria in Alzheimer’s Post-Mortem Brain. Front Aging Neurosci. 2017;9. doi:10.3389/fnagi.2017.00195

62. Westfall S, Dinh DM, Pasinetti GM. Investigation of Potential Brain Microbiome in Alzheimer’s Disease: Implications of Study Bias. J Alzheimer’s Dis. 2020;75: 559–570. doi:10.3233/JAD-191328

63. Alonso R, Pisa D, Fernández-Fernández AM, Carrasco L. Infection of fungi and bacteria in brain tissue from elderly persons and patients with Alzheimer’s disease. Front Aging Neurosci. 2018;10. doi:10.3389/fnagi.2018.00159

64. Soscia SJ, Kirby JE, Washicosky KJ, Tucker SM, Ingelsson M, Hyman B, et al. The Alzheimer’s disease-associated amyloid β-protein is an antimicrobial peptide. Bush AI, editor. PLoS One. 2010;5: e9505. doi:10.1371/journal.pone.0009505

65. Kumar DKV, Choi HS, Washicosky KJ, Eimer WA, Tucker S, Ghofrani J, et al. Amyloid-β peptide protects against microbial infection in mouse and worm models of Alzheimer’s disease. Sci Transl Med. 2016;8: 340ra72–340ra72. doi:10.1126/scitranslmed.aaf1059

66. Greathouse KL, White JR, Vargas AJ, Bliskovsky V V., Beck JA, von Muhlinen N, et al. Interaction between the microbiome and TP53 in human lung cancer. Genome Biol. 2018;19: 123. doi:10.1186/s13059-018-1501-6

67. Amplification of Full-Length 16S Gene with Barcoded Primers for Multiplexed SMRTbell® Library Preparation and Sequencing. 2019. Report No.: Part Number 101-599-700 version 02 (June 2019). Available: https://www.pacb.com/wp-content/uploads/Procedure-Checklist-Amplification-of-Full-Length-16S-Gene-with-Barcoded-Primers-for-Multiplexed-SMRTbell-Library-Preparation-and-Sequencing.pdf

68. Griffiths TL, Steyvers M. Finding scientific topics. Proc Natl Acad Sci U S A. 2004;101 Suppl: 5228–35. doi:10.1073/pnas.0307752101

69. Davis NM, Proctor DM, Holmes SP, Relman DA, Callahan BJ. Simple statistical identification and removal of contaminant sequences in marker-gene and metagenomics data. Microbiome. 2018;6: 226. doi:10.1186/s40168-018-0605-2

70. McMurdie PJ, Holmes S. phyloseq: an R package for reproducible interactive analysis and graphics of microbiome census data. PLoS One. 2013;8: e61217. doi:10.1371/journal.pone.0061217

71. Gloor GB, Macklaim JM, Pawlowsky-Glahn V, Egozcue JJ. Microbiome Datasets Are Compositional: And This Is Not Optional. Front Microbiol. 2017;8. doi:10.3389/fmicb.2017.02224

72. Harrison JG, Calder WJ, Shastry V, Buerkle CA. Dirichlet-multinomial modelling outperforms alternatives for analysis of microbiome and other ecological count data. Mol Ecol Resour. 2020;20: 481–497. doi:10.1111/1755-0998.13128

73. Holmes I, Harris K, Quince C. Dirichlet Multinomial Mixtures: Generative Models for Microbial Metagenomics. Gilbert JA, editor. PLoS One. 2012;7: e30126. doi:10.1371/journal.pone.0030126

74. Fordyce JA, Gompert Z, Forister ML, Nice CC. A Hierarchical Bayesian Approach to Ecological Count Data: A Flexible Tool for Ecologists. Scalas E, editor. PLoS One. 2011;6: e26785. doi:10.1371/journal.pone.0026785

75. Blei DM, Ng AY, Jordan MI. Latent Dirichlet Allocation. J Mach Learn Res. 2003;3: 993–1022.

76. van der Maaten L, Hinton G. Visualizing Data using t-SNE. J Mach Learn Res. 2008;9: 2579–2605. Available: http://jmlr.org/papers/v9/vandermaaten08a.html

77. McInnes L, Healy J, Saul N, Großberger L. UMAP: Uniform Manifold Approximation and Projection. J Open Source Softw. 2018;3: 861. doi:10.21105/joss.00861

78. Blei DM. Probabilistic topic models. Commun ACM. 2012;55: 77–84. doi:10.1145/2133806.2133826

79. Hofmann T. Unsupervised Learning by Probabilistic Latent Semantic Analysis. Mach Learn. 2001;42: 177–196. doi:10.1023/A:1007617005950

80. Geman S, Geman D. Stochastic relaxation, gibbs distributions, and the bayesian restoration of images. IEEE Trans Pattern Anal Mach Intell. 1984;6: 721–41. doi:10.1109/tpami.1984.4767596

81. Heinrich G. A Generic Approach to Topic Models. Proceedings of the 2009th European Conference on Machine Learning and Knowledge Discovery in Databases - Volume Part I. Berlin, Heidelberg: Springer-Verlag; 2009. pp. 517–532.

82. Heinrich G. Parameter estimation for text analysis. Leipzig, Germany; 2008. Available: https://www.arbylon.net/publications/text-est2.pdf

83. Newman MEJ, Barkema GT. Monte Carlo Methods in Statistical Physics. Clarendon Press; 1999. Available: https://books.google.com/books?id=J5aLdDN4uFwC

84. Pritchard JK, Stephens M, Donnelly P. Inference of population structure using multilocus genotype data. Genetics. 2000;155: 945–59. doi:10.1093/genetics/155.2.945

85. Wolfram Research Inc. Graph. 2010 [cited 3 Mar 2022]. Available: https://reference.wolfram.com/language/ref/Graph.html

86. Wolfram Research Inc. GraphLayout. 2021 [cited 3 Mar 2022]. Available: https://reference.wolfram.com/language/ref/GraphLayout.html

87. Wolfram Research Inc. LogitModelFit. 2008 [cited 3 Mar 2022]. Available: https://reference.wolfram.com/language/ref/LogitModelFit.html

88. Prindle A, Liu J, Asally M, Ly S, Garcia-Ojalvo J, Süel GM. Ion channels enable electrical communication in bacterial communities. Nature. 2015;527: 59–63. doi:10.1038/nature15709

89. Andrew SC, Dumoux M, Hayward RD. Chlamydia Uses K+ Electrical Signalling to Orchestrate Host Sensing, Inter-Bacterial Communication and Differentiation. Microorganisms. 2021;9: 173. doi:10.3390/microorganisms9010173

90. Mayslich C, Grange PA, Dupin N. Cutibacterium acnes as an Opportunistic Pathogen: An Update of Its Virulence-Associated Factors. Microorganisms. 2021;9: 303. doi:10.3390/microorganisms9020303

91. Vorobjeva LI. Propionibacteria. Springer Netherlands; 1999.

92. Jung J, Park W. Acinetobacter species as model microorganisms in environmental microbiology: current state and perspectives. Appl Microbiol Biotechnol. 2015;99: 2533–48. doi:10.1007/s00253-015-6439-y

93. Bitrian M, González RH, Paris G, Hellingwerf KJ, Nudel CB. Blue-light-dependent inhibition of twitching motility in Acinetobacter baylyi ADP1: additive involvement of three BLUF-domain-containing proteins. Microbiology. 2013;159: 1828–1841. doi:10.1099/mic.0.069153-0

94. Steinberg JP, Burd EM. Other Gram-Negative and Gram-Variable Bacilli. Mandell, Douglas, and Bennett’s Principles and Practice of Infectious Diseases. Elsevier; 2015. pp. 2667–2683.e4. doi:10.1016/B978-1-4557-4801-3.00238-1

95. Wu Y, Arumugam K, Tay MQX, Seshan H, Mohanty A, Cao B. Comparative genome analysis reveals genetic adaptation to versatile environmental conditions and importance of biofilm lifestyle in Comamonas testosteroni. Appl Microbiol Biotechnol. 2015;99: 3519–3532. doi:10.1007/s00253-015-6519-z

96. Wu Y, Zaiden N, Cao B. The Core-and Pan-Genomic Analyses of the Genus Comamonas: From Environmental Adaptation to Potential Virulence. Front Microbiol. 2018;9. doi:10.3389/fmicb.2018.03096

97. Alonso R, Pisa D, Carrasco L. Searching for Bacteria in Neural Tissue From Amyotrophic Lateral Sclerosis. Front Neurosci. 2019;13. doi:10.3389/fnins.2019.00171

98. Sweeney MD, Sagare AP, Zlokovic B V. Blood–brain barrier breakdown in Alzheimer disease and other neurodegenerative disorders. Nat Rev Neurol. 2018;14: 133–150. doi:10.1038/nrneurol.2017.188

99. Iliff JJ, Wang M, Liao Y, Plogg BA, Peng W, Gundersen GA, et al. A Paravascular Pathway Facilitates CSF Flow Through the Brain Parenchyma and the Clearance of Interstitial Solutes, Including Amyloid β. Sci Transl Med. 2012;4. doi:10.1126/scitranslmed.3003748

100. Stopschinski BE, Del Tredici K, Estill-Terpack S-J, Ghebremdehin E, Yu FF, Braak H, et al. Anatomic survey of seeding in Alzheimer’s disease brains reveals unexpected patterns. Acta Neuropathol Commun. 2021;9: 164. doi:10.1186/s40478-021-01255-x

101. Pritchard AB, Fabian Z, Lawrence CL, Morton G, Crean S, Alder JE. An Investigation into the Effects of Outer Membrane Vesicles and Lipopolysaccharide of Porphyromonas gingivalis on Blood-Brain Barrier Integrity, Permeability, and Disruption of Scaffolding Proteins in a Human in vitro Model. J Alzheimers Dis. 2022. doi:10.3233/JAD-215054

102. Emery DC, Cerajewska TL, Seong J, Davies M, Paterson A, Allen-Birt SJ, et al. Comparison of Blood Bacterial Communities in Periodontal Health and Periodontal Disease. Front Cell Infect Microbiol. 2021;10. doi:10.3389/fcimb.2020.577485

103. Li F, Hearn M, Bennett LE. The role of microbial infection in the pathogenesis of Alzheimer’s disease and the opportunity for protection by anti-microbial peptides. Crit Rev Microbiol. 2021;47: 240–253. doi:10.1080/1040841X.2021.1876630

104. Doty RL. The olfactory vector hypothesis of neurodegenerative disease: is it viable? Ann Neurol. 2008;63: 7–15. doi:10.1002/ana.21327

105. Capoor MN, Ruzicka F, Machackova T, Jancalek R, Smrcka M, Schmitz JE, et al. Prevalence of Propionibacterium acnes in Intervertebral Discs of Patients Undergoing Lumbar Microdiscectomy: A Prospective Cross-Sectional Study. Brüggemann H, editor. PLoS One. 2016;11: e0161676. doi:10.1371/journal.pone.0161676

106. Jacovides CL, Kreft R, Adeli B, Hozack B, Ehrlich GD, Parvizi J. Successful identification of pathogens by polymerase chain reaction (PCR)-based electron spray ionization time-of-flight mass spectrometry (ESI-TOF-MS) in culture-negative periprosthetic joint infection. J Bone Joint Surg Am. 2012;94: 2247–54. doi:10.2106/JBJS.L.00210

